# The SmARTR pipeline: a modular workflow for the cinematic rendering of 3D scientific imaging data

**DOI:** 10.1101/2024.07.03.601651

**Authors:** Simone Macrì, Nicolas Di-Poï

## Abstract

Advancements in non-invasive surface and internal imaging techniques, along with computational methods, have revolutionized 3D visualization of organismal morphology. These breakthroughs not only enhance research and medical anatomical analysis, but also facilitate the preservation and digital archiving of scientific specimens. We introduce the SmARTR pipeline (Small Animal Realistic Three-dimensional Rendering), a comprehensive workflow integrating wet lab procedures, 3D data acquisition, and processing to produce photorealistic 3D scientific data through cinematic rendering. This versatile pipeline supports multi-scale visualizations, from tissue-level to whole-organism details across diverse living organisms and is adaptable to various imaging sources and platforms. Its modular design and customizable rendering scenarios, provided by detailed SmARTR networks in a free software environment, make it a powerful tool for 3D data analysis. Accessible to a broad audience, SmARTR serves as a valuable resource not only for multiple fields of research in life sciences but also for education, diagnosis, outreach, and artistic endeavors.

## INTRODUCTION

The study of morphology is crucial to understand the basic connections linking organismal form, function, and environment. Indeed, a comprehensive understanding of both the external characteristics (eidonomy) and internal structure (anatomy) of animals is foundational to a wide range of scientific fields, which include but are not limited to zoology, embryology, physiology, pathology, and evolution, helping to unravel the core processes of an organism’s life and behavior. Scientific morphological studies, tracing back to ancient civilizations, recognize Aristotle (384– 322 BC) as a pioneering figure. He laid the groundwork for comparative anatomy through dissections of over 500 vertebrate and invertebrate species. His meticulous observations, documented with systematic terminology in comprehensive treatises^1,2^, likely included diagrams and pictorial tables alongside textual descriptions^1,3^. Although not preserved today, his illustrative opus, referred to as *Anatomai*, probably featured detailed drawings of both animal’s eidonomy and anatomy^1,4^. Andreas Vesalius (1514–1564) further advanced anatomy with his work *De humani corporis fabrica* (1543). His detailed illustrations, combined with scientific rigor, paved the way for modern biomedical sciences^5–9^ and underscored the importance of images in comprehending and communicating organismal morphology^10^. Following Vesalius’ work, artists were drawn to create accurate anatomical depictions, interweaving art and science. This integration, fueled by technical innovations, evolved over time. Scientists like Maria Sibylla Merian (1647-1717)^11,12^, Robert Hooke (1635-1703)^13–15^, Ernst Haeckel (1834-1919)^16,17^, Johan Erik Vesti Boas (1855-1935), and Simon Paulli (1865-1933)^18,19^ exemplify how this synergy enhances our understanding of nature’s diversity and life’s complexity^20,21^.

Nowadays, dramatic technological breakthroughs persist in advancing the study and visualization of organism morphology to levels of detail unprecedented in previous eras, surpassing past anatomistś expectations. Three-dimensional (3D) imaging technologies have revolutionized medical, veterinary, and biological disciplines, profoundly impacting diagnostics, pathology, and various areas of life sciences research. Additionally, they have fostered the conservation and expanded the impact of scientific and natural history museum collections by enabling the creation of digital copies of precious and delicate specimens^22–24^. Non-invasive techniques like photogrammetry and surface scanning capture detailed morphological information of the outer surface of samples^25^, while X-ray microcomputed tomography (µCT) and magnetic resonance imaging (MRI) reveal their intricate internal structure^25,26^. These techniques are increasingly employed to support qualitative and quantitative morphological analyses in living organisms^27–36^. Recently, µCT digital imaging has gained momentum due to improved scanner availability, cost-effectiveness, and superior micrometer-level spatial resolution compared to ordinary MRI^25^. Moreover, contrast-enhancing agents have expanded µCT’s capability beyond mineralized animal tissues like bones and teeth, making it more appealing for morphological studies^37–39^. Commonly used contrast agents, including Iodine (I_2_) (either dissolved in ethanol (I_2_E) or methanol (I_2_M), or in the form of aqueous Lugol solution (I_2_KI)), phosphotungstic acid (PTA), phosphomolybdic acid (PMA), and osmium tetroxide (OsO_4_), exhibit distinct diffusive potential, toxicity profiles, and varying degrees of affinity for enhancing soft tissue contrast^39–43^. Owing to their low toxicity, high diffusion rate, contrast-enhancing potential, and the ease of removal from specimens, iodine solutions are widely used as soft tissue staining compounds for large biological sample µCT imaging^28,30,44–50^. Larger molecules such as PTA or PMA provide excellent soft tissue contrast but permeate samples at a slow rate^37,39,43^, making them suitable for staining smaller or poorly keratinized specimens such as early-stage vertebrate embryos or exposed internal organs^51–57^. More recently, phase-contrast X-ray imaging^58–61^ has emerged as a valuable modality for soft tissue imaging without contrast agents. However, achieving detailed resolution with this technique necessitates either very small biological samples or advanced and expensive equipment (e.g., synchrotrons) to match the spatial resolution of contrast enhanced µCT.

Accurate visualization of 3D morphology with minimal deformation is critical for basic research in life sciences, educational purposes, and veterinary medicine. Additionally, as 3D imaging data becomes increasingly relevant across diverse scientific domains, ensuring its accuracy and reliability remains essential. Recent advancements in computational power and rendering algorithms^62–69^ have significantly advanced the visualization of complex 3D datasets. In particular, direct volume rendering (DVR) algorithms, which utilize global illumination models and Monte Carlo integration^70^, accurately simulate light-ray paths through methods like ray tracing^65^ or path tracing^71^. These algorithms can reproduce effects such as indirect lighting, light reflection/refraction from curved surfaces (caustics), blurry reflections, and soft shadows, dramatically enhancing the realism of generated 3D volumes^65,71–75^. Referred to as cinematic or photorealistic rendering^74,76^, these DVR techniques—largely implemented through commercial platforms usually bundled with expensive imaging hardware—are gaining momentum in the human medical field to support diagnostics, surgery, and anatomy teaching^73,77–84^, while revitalizing the connection between science and impactful anatomical illustrations. In contrast to this dynamic landscape, the potential impact of cinematic rendering remains largely unexplored across most biological fields, yet significant benefits are anticipated from incorporating this method into the experimental routine of various disciplines. In this study, we developed the SmARTR pipeline (Small Animal Realistic Three-dimensional Rendering), a scalable, modular, and versatile workflow that seamlessly integrates wet lab procedures, imaging, and advanced volumetric data processing. By leveraging the state-of-the-art Monte Carlo path tracing framework embedded in free MeVisLab SDK (MeVis Medical Solutions AG and Fraunhofer MEVIS, Germany), an image processing platform distributed for non-commercial use, our pipeline generates photorealistic 3D scientific data through cinematic rendering. Several modular networks have been meticulously designed and prototyped for use in a diverse array of living organisms and tissues. Additionally, a detailed step-by-step procedure has been developed and made accessible to a broad biological audience. The potential applications of the SmARTR pipeline are far-reaching, spanning diverse fields in life sciences research such as evolutionary and developmental biology (evo-devo), comparative anatomy, physiology, paleontology, zoology, diagnostics, and veterinary medicine, as well as extending to education, outreach, and art.

## RESULTS AND DISCUSSION

### Experimental design

The informative power of 3D non-invasive imaging techniques such as µCT, MRI, phase-contrast imaging, and optical coherence tomography (OCT) heavily relies on post-processing software. This software utilizes algorithms like filtered convolution back-projection and inverse fast Fourier transform to interpret the raw images acquired from scanners and faithfully reconstruct the sample’s morphology into a stack of grayscale cross-sectional images^37,85,86^. To visualize the sample as a full 3D volume, rendering techniques like indirect volume rendering (IVR) and DVR are crucial^87–93^. IVR generates an intermediate, sharp, and opaque polygonal mesh that approximates the sample’s outer surface, while DVR directly visualizes the entire volumetric data, preserving all 3D scan-derived information and offering comprehensive insights into sample thickness, texture, and internal microarchitecture^90,91,94,95^. Enhanced by segmentation of anatomical regions of interest and virtual volume sectioning operations, DVR algorithms facilitate a detailed appreciation of complex anatomical structures and spatial relationships within biological samples. Indeed, volume rendering is crucial not only in medical and veterinary diagnostics^96–107^, pre-operative investigations^108–116^, and post-operative follow-up^114,117–120^ but also in diverse qualitative and quantitative analyses widely employed in scientific disciplines such as paleontology^121–125^, evo-devo^29,126,127^, ecology^47,128–132^, and zoology^28,30,36,48,49,133–139^. However, although traditional DVR implemented through ray casting technology can generate aesthetically appealing images, it relies on simple lighting and local illumination techniques that hinder the production of realistic representation of 3D data^140–142^. In contrast, Monte Carlo path tracing method^70^ excels in modeling global illumination, recreating lifelike shadows and reflections by simulating thousands of light rays, thus enhancing depth perception and creating cinematic 3D renderings^143,144^. Yet, the stochastic nature of light path approximation requires a variable number of iterations for convergence, causing volume interactions to lag on low-end graphics processing unit (GPU) systems. To explore the potential advantages offered by Monte Carlo path tracing algorithms in 3D visualizations of biological data beyond the human medical domain, and to address the fidelity of the increasing amount and importance of 3D imaging data in various scientific fields^23,24^, we developed a modular and scalable workflow—the SmARTR pipeline—that encompasses all steps from small animal or organ sample collection to final generation of cinematic 3D renderings (Figure 1).

**Figure 1.**
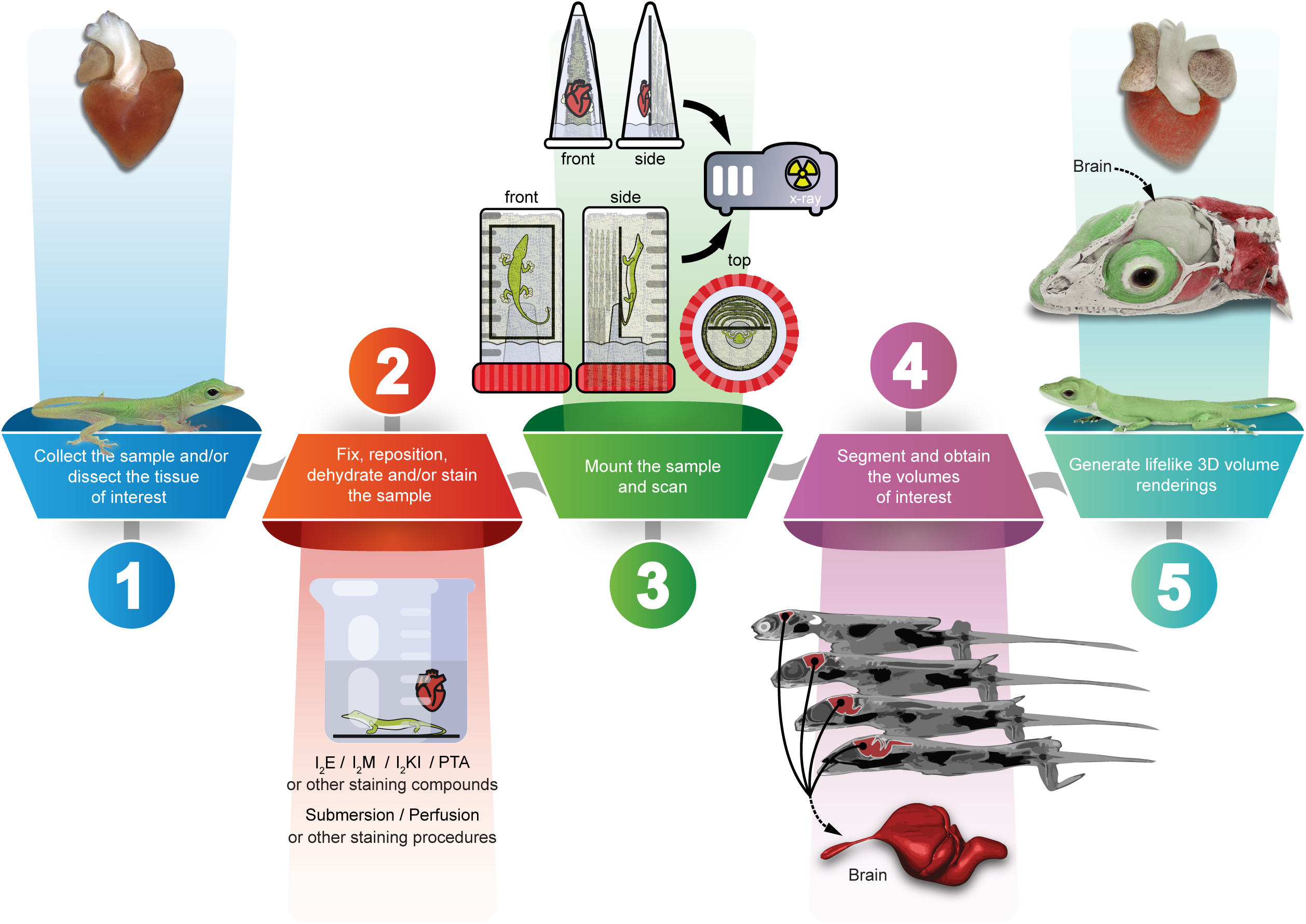
Overview of the SmARTR pipeline adapted for µCT imaging. Graphical representation depicting the multi-step workflow (steps 1-5) of the SmARTR pipeline applied to µCT imaging. Images on the left (step 1) show a newborn *A. carolinensis* (bottom) and a *P. vitticeps* dissected heart (top). The 3D rendered versions of these processed specimens, including a cutaway view of the lizard’s head highlighting the skull and brain, are displayed on the right (step 5). The final rendering framework is versatile, capable of processing 3D data derived from any imaging modality that aligns with specific research objectives. While optimized for µCT, the modular design of the SmARTR pipeline offers flexibility to accommodate other imaging modalities, such as MRI, phase-contrast X-ray imaging, and even 3D microscopy. Abbreviations: I_2_E, iodine dissolved in ethanol; I_2_M, iodine dissolved in methanol; I_2_KI, iodine-potassium iodine or Lugoĺs solution; PTA, phosphotungstic acid.

### Sample preparation and volumetric data rendering

We designed each step of the pipeline to enhance its applicability across a broad spectrum of animal samples, ranging from vertebrate embryos and postnatal stages to echinoderms and insects, as well as a wide selection of organs and tissues derived from dissected specimens (Figure 1, step 1). Additionally, this approach is expected to work for non-animal biological samples, including plants and fungi, or fossilized specimens, utilizing similar 3D imaging methodologies. Importantly, the pipeline enables multi-scale visualizations, spanning from tissue to whole-organism details, and is versatile in terms of 3D imaging sources and platforms, as well as soft tissue staining agents (Figure 1, steps 2 and 3). In this study, we used μCT for the pipeline example, and our samples were stained with either 1% I_2_E, 0.7% PTA, or a combination of both compounds for dissected organs. However, alternative staining agents or methods beyond submersion^30,37–39,145–148^, known to maximize soft tissue contrast in particular samples or to enhance specific organ differentiation, may be employed (Figure 1, step 2). Regarding specimen mounting, we devised a simple methodology (Figure 1, step 3) to position samples, enabling optimal imaging and rendering conditions while preserving sample integrity in terms of both morphology and anatomy. Instead of wrapping the sample in tissue paper moistened with contrast agent solvent to maintain humidity and prevent dehydration, we created a custom chamber utilizing common lab equipment (Supplemental Figure S1), which allows the sample to be surrounded by air. This prevents staining agent transfer to the wrapping tissues and facilitates sample isolation during the rendering step. This setup consistently preserved sample integrity and position even during prolonged imaging sessions. Moreover, although we cannot guarantee this for all specimens, mounting with flexible cyanoacrylate glue has proven to be reversible. After scanning, samples could be easily detached from the mounting slide without any tissue damage and processed further, if required, or stored. Importantly, while sample preparation and μCT imaging yielded excellent results across the wide range of sample types and sizes used in the study, specific contexts may benefit from alternative mounting or imaging acquisition to better fit particular aims.

Versatility was a key criterion guiding the design of the pipeline steps involving volumetric data manipulation and rendering (Figure 1, steps 4 and 5). Utilizing the broadly customizable visualization framework offered by free MeVisLab SDK, the software platform used for the final rendering step, six main modular networks (SmARTR networks, Table 1) were specifically designed, developed, and prototyped for a wide range of biologists. The SmARTR_Pattern_Drawing network can be used to design simple color schemes on volumes recreating sample natural coloration for illustrative purposes, while the SmARTR_Single_Volume, SmARTR_Multi-Mask_Volume, and SmARTR_Multi-Independent_Volume networks allow visualization and modification—by performing volume cutting operations—of either single or multiple volumes in a scene. The SmARTR_Nested_Multi-Volume and SmARTR_Advanced_Multi-Volume networks support elaborate visualizations, suitable to illustrate the complex relationships between animal organs within body walls or display single organ internal structure. A detailed procedure for all networks has been developed and made accessible to a broad audience (Supplemental Methods). Although designed with various visualization scenarios in mind, the SmARTR networks inherently possess broader applications and can be easily expanded, combined, or modified to adapt to particular needs. Furthermore, pre-visualization operations such as segmentation of structures of interest, data resampling, and image registration, if required, can be performed using any 3D imaging software platform that supports these processes. Examples include Amira, Avizo, 3D Slicer, VGStudio Max, Dragonfly, Fiji, ITK-Snap, and Mimics, all of which support data conversion to numerous formats compatible with MeVisLab (Supplemental Methods, section 2.2, panel d2). Given the routine nature of these tasks, we thought that researchers or scientists could use their preferred software to perform such steps rather than detailing these operations in MeVisLab, thus avoiding unnecessary additional complexity in the pipeline. In addition, while the full pipeline has been conceived and optimized for µCT imaging, volumetric data obtained with different methodologies, like MRI or phase-contrast imaging, and even 3D fluorescence microscopy can also be processed using the SmARTR visualization framework.

**Table 1.**
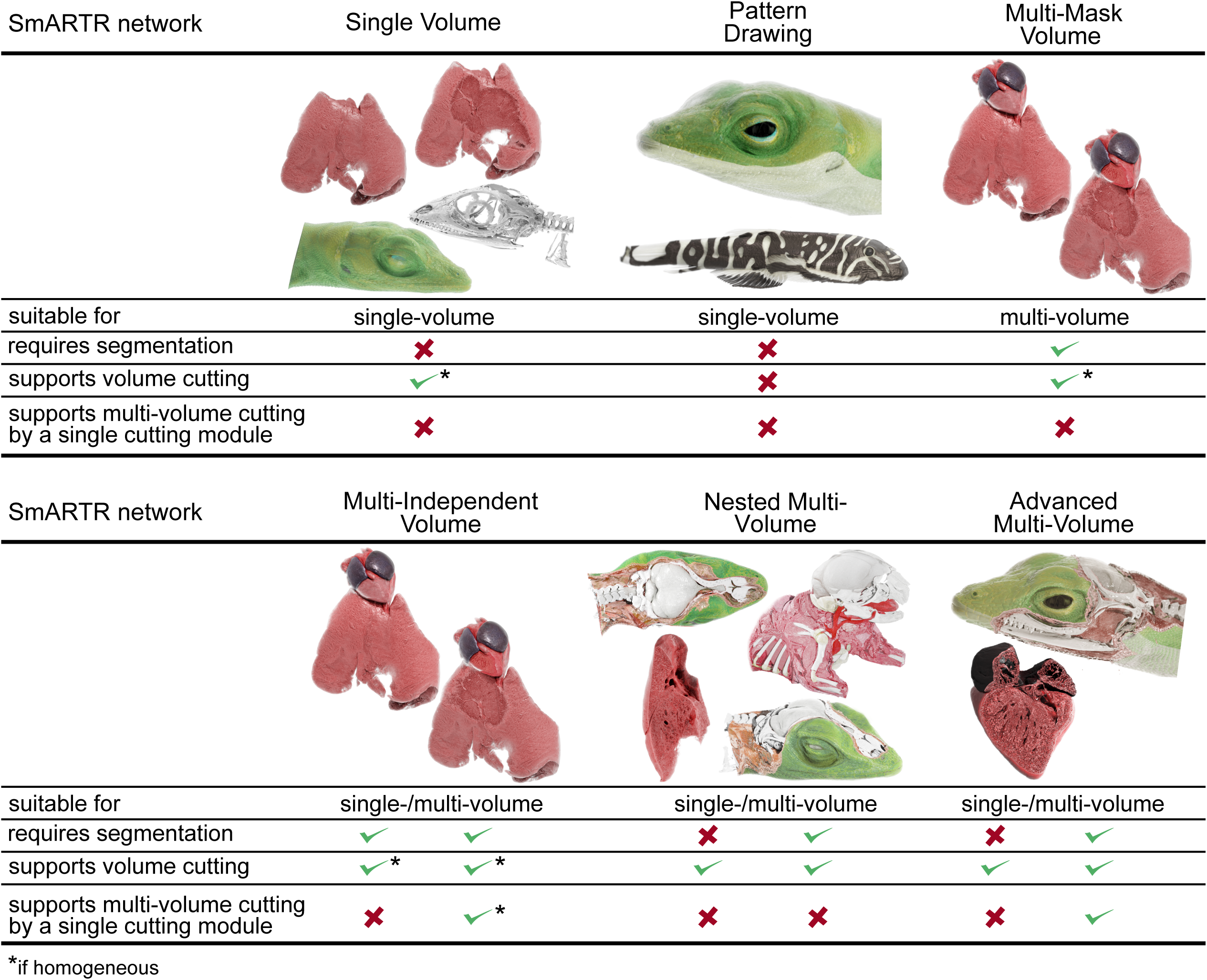
Properties of the SmARTR networks. Table highlighting the requirements, volume operations, and rendering purposes of each SmARTR network.

| SmARTR network | Single Volume |  | Pattern Drawing |  | Multi-Mask Volume |
| --- | --- | --- | --- | --- | --- |
| suitable for | single-volume |  | single-volume |  | multi-volume |
| requires segmentation | ✗ |  | ✗ |  | ✓ |
| supports volume cutting | ✓* |  | ✗ |  | ✓* |
| supports multi-volume cutting by a single cutting module | ✗ |  | ✗ |  | ✗ |

| SmARTR network | Multi-Independent Volume |  | Nested Multi-Volume |  | Advanced Multi-Volume |  |
| --- | --- | --- | --- | --- | --- | --- |
| suitable for | single-/multi-volume |  | single-/multi-volume |  | single-/multi-volume |  |
| requires segmentation | ✓ | ✓ | ✗ | ✓ | ✗ | ✓ |
| supports volume cutting | ✓* | ✓* | ✓ | ✓ | ✓ | ✓ |
| supports multi-volume cutting by a single cutting module | ✗ | ✓* | ✗ | ✗ | ✗ | ✓ |

\*:if homogeneous

### SmARTR networks and pipeline applications for full-body and organ-level views

Taking advantage of the global light simulation implemented by the MeVis path tracer, the SmARTR visualization pipeline allows the accurate reproduction of small animal gross and fine anatomical features, making it a valuable and multi-purpose tool in various life science fields (Figure 2). The increasing use and importance of 3D imaging techniques as investigative tools in different domains of basic, medical, and veterinary research in life sciences, coupled to recent collaborative efforts and initiatives between scientists, natural history museums, and zoological gardens to digitize^23^ and publicly share a wide variety of specimens—like the oVert (openVertebrate) network^24^ and the Morphosource repository (http://www.MorphoSource.org/)— and the creation of specimen data aggregators—like VertNet, GBIF, DiSSCO, and iDigBio^22^—are collectively increasing the availability of anatomical data from diverse and numerous biological samples. In this context, our SmARTR visualization framework can effectively exploit these extensive collections, amplifying the educational scope and scientific outreach, while also improving various research fields (Figure 2). Photorealistic renderings and illustrations of intact animals featuring uniform coloration can be realized using the SmARTR_Single_Volume network (Table 1, Figure 3A and section 2 of Supplemental Methods). For specimens displaying patterns or skin markings, a specifically designed network, the SmARTR_Pattern_Drawing_network (Table 1, Figure 3B-D and section 6 of Supplemental Methods), is employed to generate chromatic motifs on volumes, approximating their natural coloration. While complex color schemes cannot be achieved, as they are typically the prerogative of methodologies other than internal imaging and DVR techniques, the network allows multiple patterns to be generated and rendered as independent structures, enhancing the realism and lifelikeness of the samples. This ability to reproduce simple skin patterns featured by the selected specimens on the volumetric data could represent an advanced tool for creating and showcasing virtual museum collections, taxonomic identification, enhancing understanding of biological processes, as well as producing educational material. This could take the form of physical or digital flashcards/flyers or interactive models, highlighting species’ main characteristics or distinctive features. Moreover, by leveraging the growing availability, usability, and reliability of generative AI tools that create images from user text prompts, rich and intricate natural habitats that faithfully reproduce a species’ environment can be generated, further enhancing the educational value of such illustrations (Figure 3).

**Figure 2.**
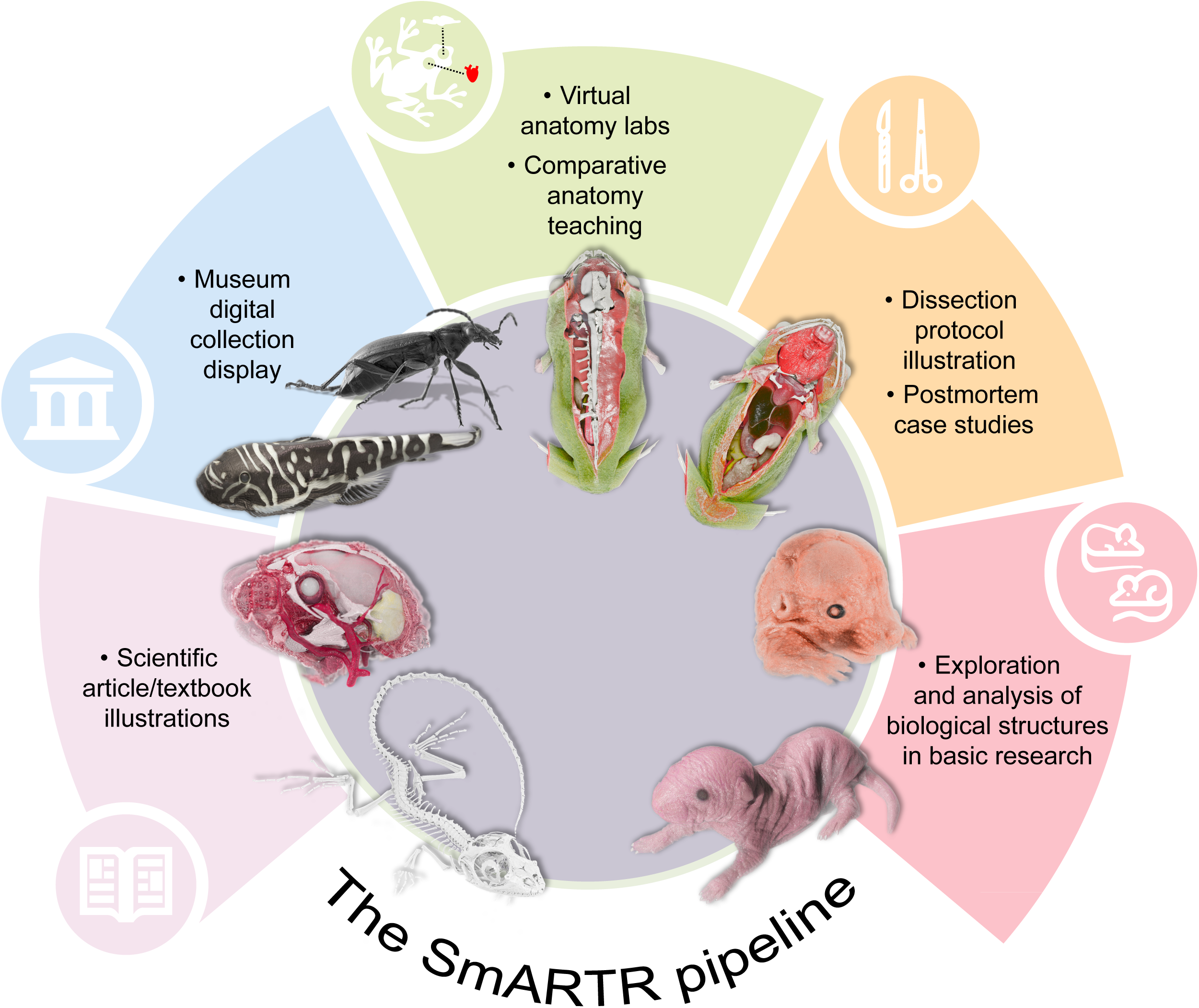
Visualization capabilities and applications of the SmARTR pipeline. Schematic summarizing the extensive opportunities for visualization offered by the SmARTR pipeline and its potential contributions to various biological and biomedical fields, scientific outreach, and education. Silhouettes in circles are from Iconduck (https://iconduck.com/), courtesy of the following sources (clockwise order starting from the bottom left): Windows Icons (used without modification, CC-BY 3.0 https://creativecommons.org/licenses/by/3.0/), OSM Map Icons (used without modification, CC0 1.0 https://creativecommons.org/publicdomain/zero/1.0/), Luan Himmlisch (modified from original CC-BY 4.0 https://creativecommons.org/licenses/by/4.0/), Resolve To Save lives (used without modification, CC0 1.0 https://creativecommons.org/publicdomain/zero/1.0/), Vectopus (modified from original, MIT license https://choosealicense.com/licenses/mit/).

**Figure 3.**
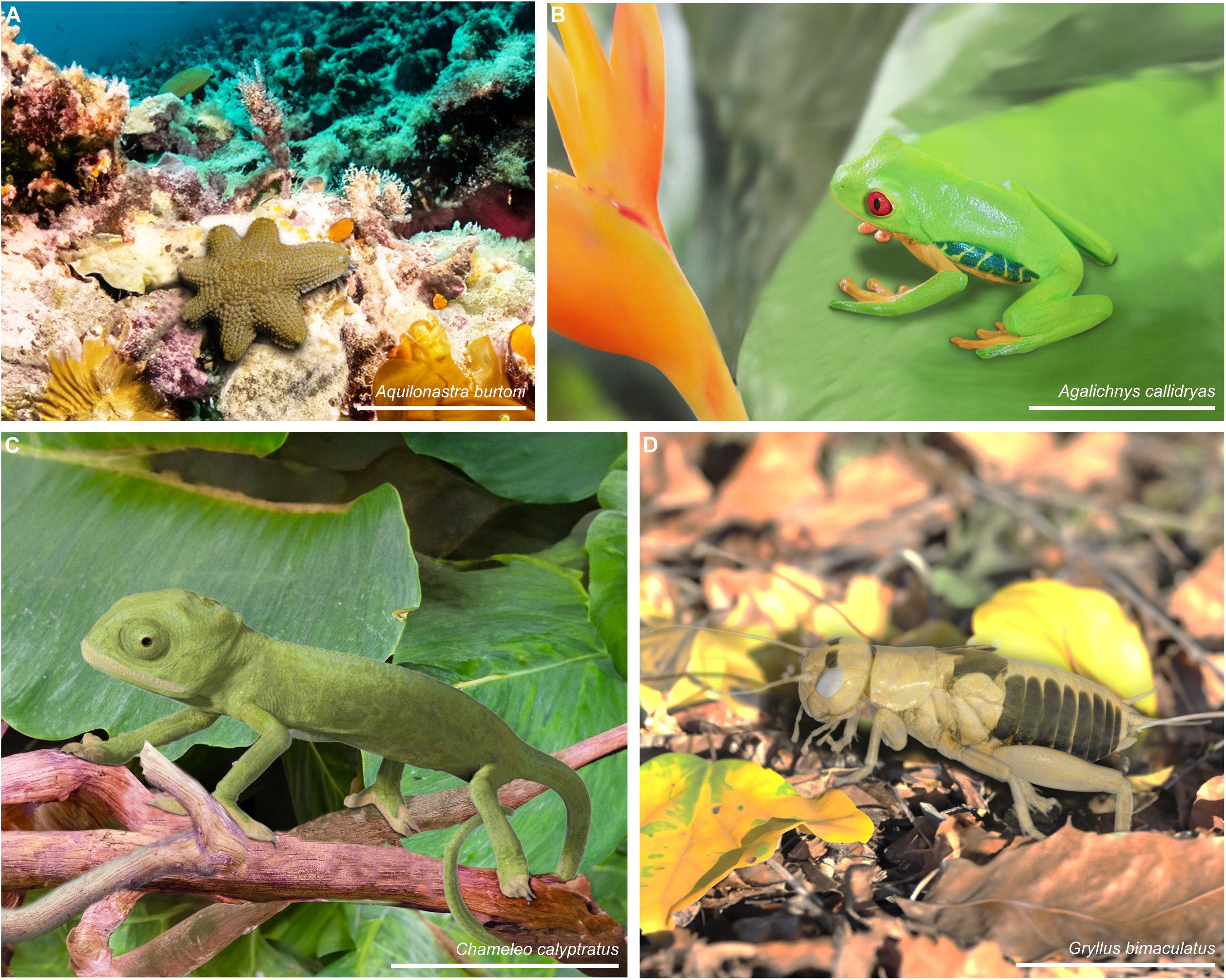
SmARTR-enhanced renderings of intact specimens. Cinematic rendering reproducing the natural coloration of invertebrate (*A. burtoni* (A); *G. bimaculatus* (D)) and vertebrate (*A. callidryas* (B); *C. calypratus* (C)) species against AI-generated backgrounds that recreate their natural environments. The eyes of *A. callidryas* and *C. calyptratus* specimens were enhanced by overlaying photographs using photo editing software. Scale bars, 1 cm.

Yet, the strength of the SmARTR networks is their capability to extract the full potential residing within contrast-enhanced samples obtained from various 3D imaging technologies such as μCT, providing detailed views of small animals’ internal anatomy and organ structure. Indeed, the different SmARTR networks offer a wide array of visualization options, each with numerous potential applications, designed to cover a wide spectrum of specific user needs (Table 1 and Supplemental Methods). Single- or multi-volume/-organ visualization is essential for understanding biological complexity, improving qualitative descriptions and/or complementing quantitative analyses of species-specific anatomical traits in both extant and extinct species^30,33,47,122,124,134,135,137,149–153^. Especially, detailed visualizations of animal visceral organs, their reciprocal relationships, and their distribution within skeletal elements, as well as isolated organs or animal parts (Figure 4) can be easily obtained using either the SmARTR_Multi-Mask_Volume or SmARTR_Multi-Independent_Volume network (Table 1 and sections 3-4 of Supplemental Methods). These two prototyped networks represent alternative ways to render multiple subvolumes deriving from the same scan data—such as multiple organs and skeletal elements (Figure 4)—in the same 3D scene. Both networks use segmentation masks to either “label” the corresponding subvolumes on the whole scan volume or extract these subvolumes to represent them as structures independent from the original volume. The first strategy (section 3 of Supplemental Methods) relies on the possibility to generate multiple instances (copies) of the whole scan volume, each displaying only the subvolume specified by the corresponding segmentation mask. The alternative multi-volume rendering strategy (section 4 of Supplemental Methods) uses a separate accessory network (SmARTR_Volume_Extraction_Network; section 4 of Supplemental Methods) to extract the subvolumes of interest from the whole scan volume, rendering them as individual structures in the SmARTR_Multi-Independent_Volume network. Though both strategies can lead to the same results, they allow different types of operations on volumes (Table 1 and sections 3-4 of Supplemental Methods). Of note, although the focus of the SmARTR pipeline is on photorealism, artificial colors can be selected to mark specific anatomical structures while keeping other material properties unaltered (section 2.2 (step 2c) of Supplemental Methods). Beyond their integration into educational materials, these multi-volume representations have far-reaching implications across various domains, enhancing scientific articles, supporting textual descriptions in scientific books, and improving the understanding of anatomy and pathology, thereby increasing the impact of such resources. Similarly, these networks could provide valuable insights for basic research in life sciences, including developmental processes and abnormalities, potentially leading to advances in regenerative medicine, while facilitating detailed study of the biomechanics of biological systems or fossil specimens in unprecedented depth.

**Figure 4.**
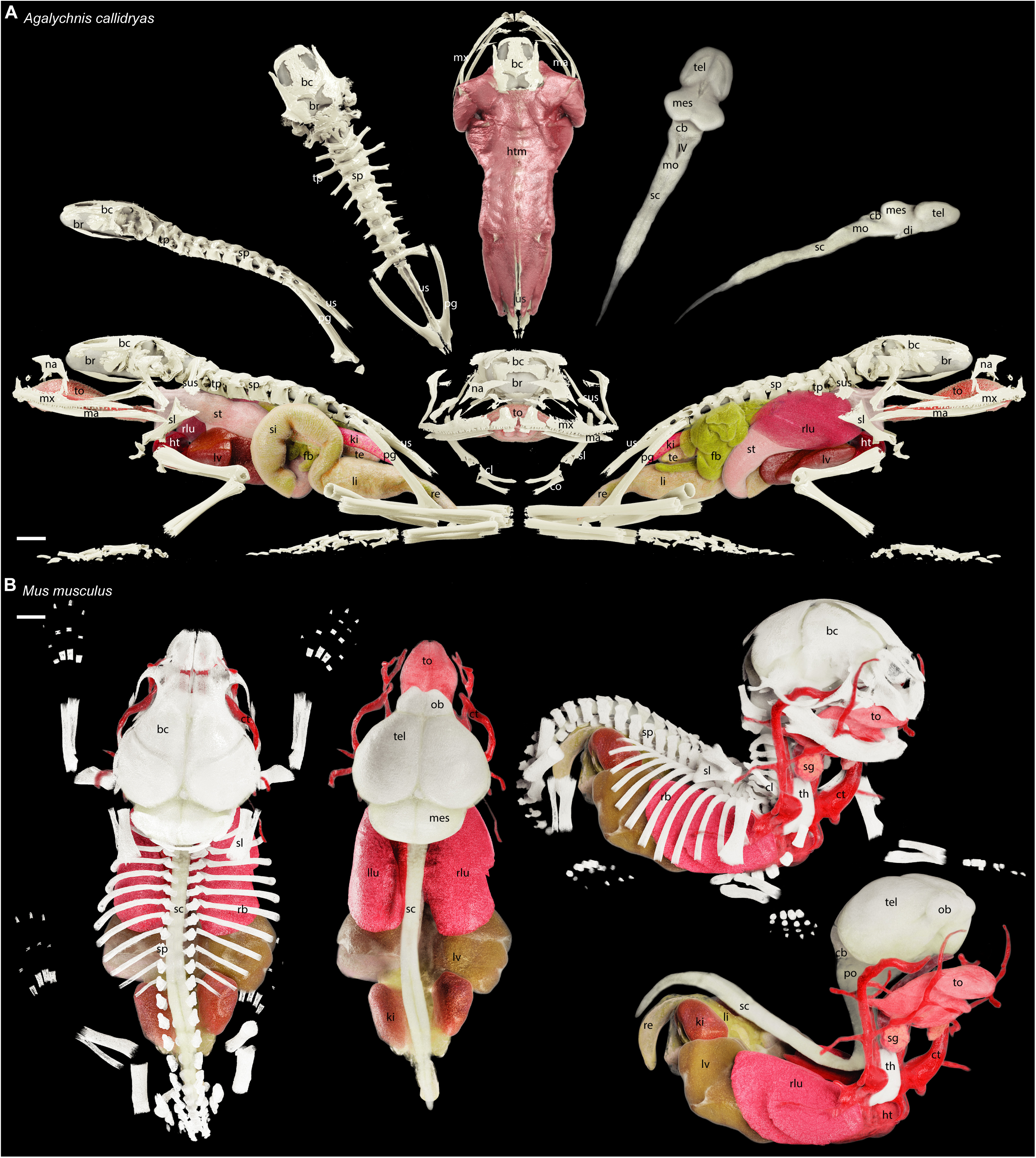
Cinematic rendering of vertebrate bony skeleton and internal organs enabled by the SmARTR pipeline. (A) Detailed renderings of the skeletal elements and major internal organs of *A. callidryas*. Descriptions of the panels follow a clockwise arrangement starting from the bottom left: left lateral view of the bony skeleton and major internal organs (left lung removed for visibility of medial structures); lateral view of the braincase, spine, and pelvic girdle; dorsal view of the braincase, spine, and pelvic girdle; dorsal view of the axial musculature integrated with head, spine, and pelvic girdle bones; dorsal view of the brain; lateral view of the brain; right lateral view of the bony skeleton and internal organs. A frontal view of the head skeleton and major internal organs is shown in the middle panel. (B) Detailed renderings of the skeletal elements and major internal organs of *M. musculus*. Descriptions include from left to right: dorsal view of major internal organs within the bony skeleton; dorsal view of the major internal organs; right lateral view of major internal organs within the bony skeleton; right lateral view of major internal organs. Abbreviations: bc, braincase; br, brain; cb, cerebellum; cl, clavicle; co, coracoid; ct, carotid; di, diencephalon; fb, fat body; ht, heart; htm, head and trunk muscles; ki, kidney; IV, fourth ventricle; li, large intestine; llu, left lung; lv, liver; ma, mandible; mes, mesencephalon; mo, medulla oblongata; mx, maxilla; na, nasal bone; ob, olfactory bulb; pg, pelvic girdle; po, pons; rb, rib; re, rectum; rlu, right lung; sc, spinal cord; sg, salivary gland; si, small intestine; sl, scapula; sp, spine; st, stomach; sus, suprascapular; te, testis; tel, telencephalon; th, thymus; to, tongue; tp, transverse process; us, urostyle. Scale bars, 1mm.

### SmARTR networks and pipeline applications for cutaway views

Another key feature of most of the main SmARTR networks is the possibility to perform cuts— either within a selected volume only or simultaneously across multiple volumes in the scene (Table 1)—using freehand contours drawn directly in the software viewer window. Such an implementation allows for cutting along complex, curved paths and, depending on the sample, can significantly facilitate the appreciation of the intricate relationships between the various organs and systems of an organism (Figure 5) or aid in fully exploring organ internal structure (Figure 6), often difficult to visualize in detail when using flat cutting planes. Cutaway views of single, non-segmented, and homogeneous volumes, such as structures characterized by an external and internal composition that can be well approximated by a single appearance profile (color, light interaction, etc.), like an unlayered single organ or animal part, can be easily obtained with the SmARTR_Single_Volume_network (Table 1 and section 2 of Supplemental Methods). In some cases, the volumes of interest can be nested within each other, such as the compartments of an organ, the head with its skull and brain, and other head soft tissues like the gills, or the body and its visceral organs (Figure 5). In these scenarios, unless each structure exposed by cutting operations is individually segmented and rendered (Figure 5A, top panel), the realism of the scene is significantly limited. This is because only one appearance profile can be assigned to each volume, while the external surface of the outermost volume (e.g., the skin of a specimen’s head) often does not match the appearance of its internal structures (e.g., the head and neck muscles). By leveraging a specifically designed integration of modules, the SmARTR_Nested_Multi-Volume network (section 5 of Supplemental Methods) produces photorealistic cutaway views to reveal inner body structures, like distinct tissues of an organ (Figure 5A, bottom panel), the skull, the brain, and other soft tissues within the head (Figure 5B), or the skeleton and the visceral organs inside the body walls (Figure 5C). Importantly, the strategy developed in this network saves time by bypassing the need for extensive segmentation of intermediate structures when creating photorealistic multi-organ cutaway views. In fact, volume cutting operations performed in this network generate a supplementary volume at the interface between the exposed surface and the overlaying space. This new volume can be rendered independently to match the internal appearance of the sectioned volume, thereby enhancing scene realism. Of note, the SmARTR_Nested_Multi-Volume can also be selected to provide an additional level of refinement when generating cuts through a single homogeneous volume (Figure 6A, left panel) or when performing such operations on single non-homogeneous volumes (Figure 6A, right panel). Finally, enhanced cutaway views of multiple volumes that share the same internal structure but differ, even slightly, in surface color, such as the atrial and ventricular heart regions (Figure 6B) or varying skin patterns of a fish (see Figure 5B, top panel), can be achieved with the SmARTR_Advanced_Multi-Volume network (Table 1 and section 7 of Supplemental Methods), which extends the strategy implemented in the SmARTR_Nested_Multi-Volume network to multiple volumes.

**Figure 5.**
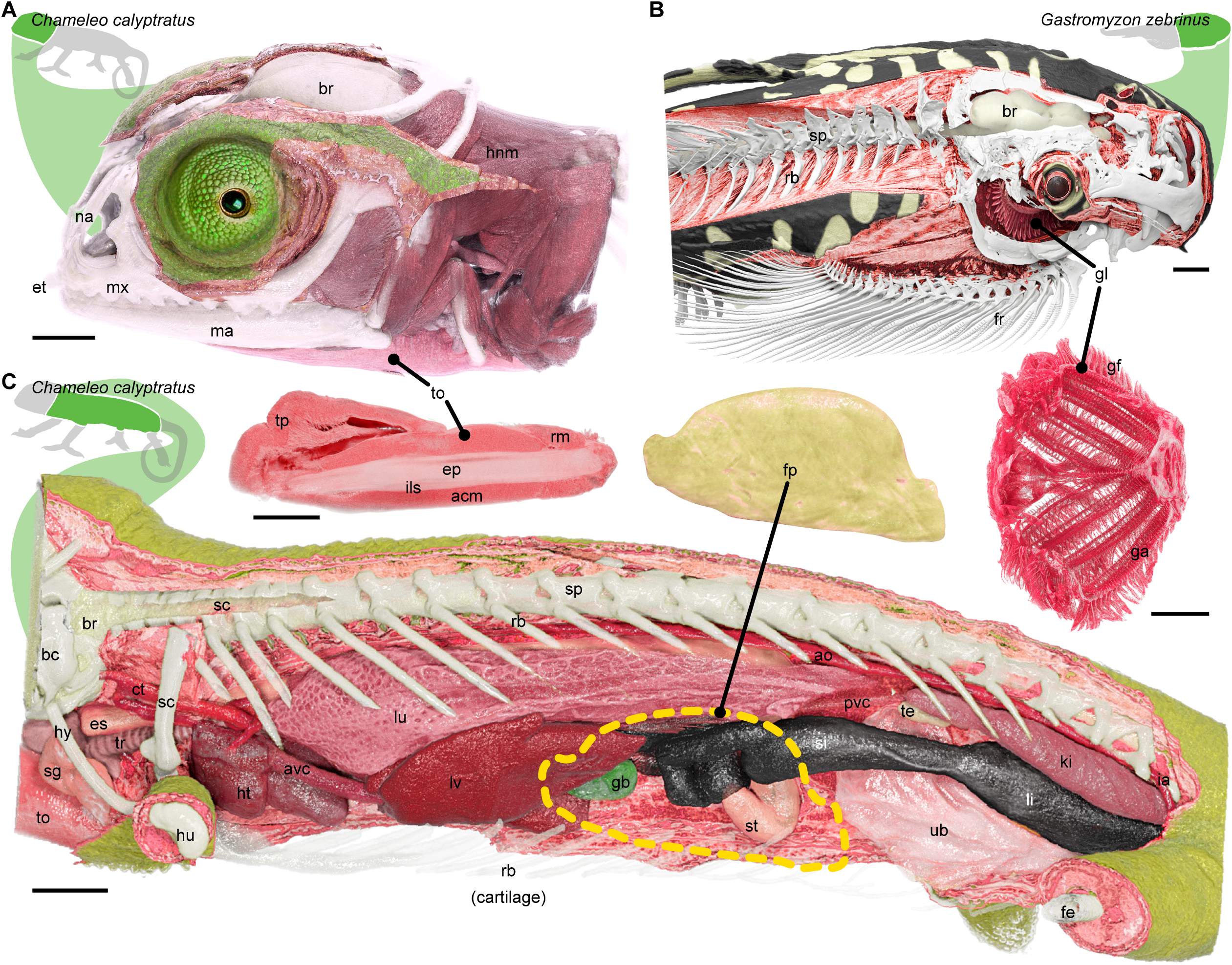
Cinematic cutaway views of vertebrate samples implemented with the SmARTR networks. (A) Lateral cutaway view of the head (top panel) of *C. calyptratus*, revealing the spatial relationships between the skull bones, brain, head and neck muscles. The eye was enhanced by overlaying a photograph using photo editing software. The bottom panel provides a detailed view of the tongue, illustrating its various components. (B) Cutaway view of the head and anterior trunk region (top panel) of *G. zebrinus*, displaying the skeletal elements and muscles. Cuts in the skull expose the brain and gills, with the latter shown in a magnified view in the bottom panel. (C) Details of the internal anatomy of *C. calyptratus*. The ribs (rb) have been cut, and the fat pad (fp, top panel) has been removed from its original position (dashed yellow line) to enhance visualization of organ morphology and spatial configuration within the body cavity. Abbreviations: acm, accelerator muscles; ao, aorta; avc, anterior vena cava; bc, braincase; br, brain; ct, carotid; ep, entoglossal process; es, esophagus; et, egg tooth; fe, femur; fp, fat pad; fr, fin rays; ga, gill arch; gb, gall bladder; gg, gill filaments; gl, gills; hnm, head and neck muscles; ht, heart; hu, humerus; hy, hyoid; ia, iliac artery; ils, intralingual sheats; ki, kidney; li, large intestine; lu, lung; lv, liver; ma, mandible; mx, maxilla; na, nasal; pvc, posterior vena cava; rb, ribs; rm, retractor muscles; sc, spinal cord; sg, salivary gland; si, small intestine; sp, spine; st, stomach; te, testis; to, tongue; tp, tip pad; tr, trachea; ub, urinary bladder. Scale bars, 1 mm.

**Figure 6.**
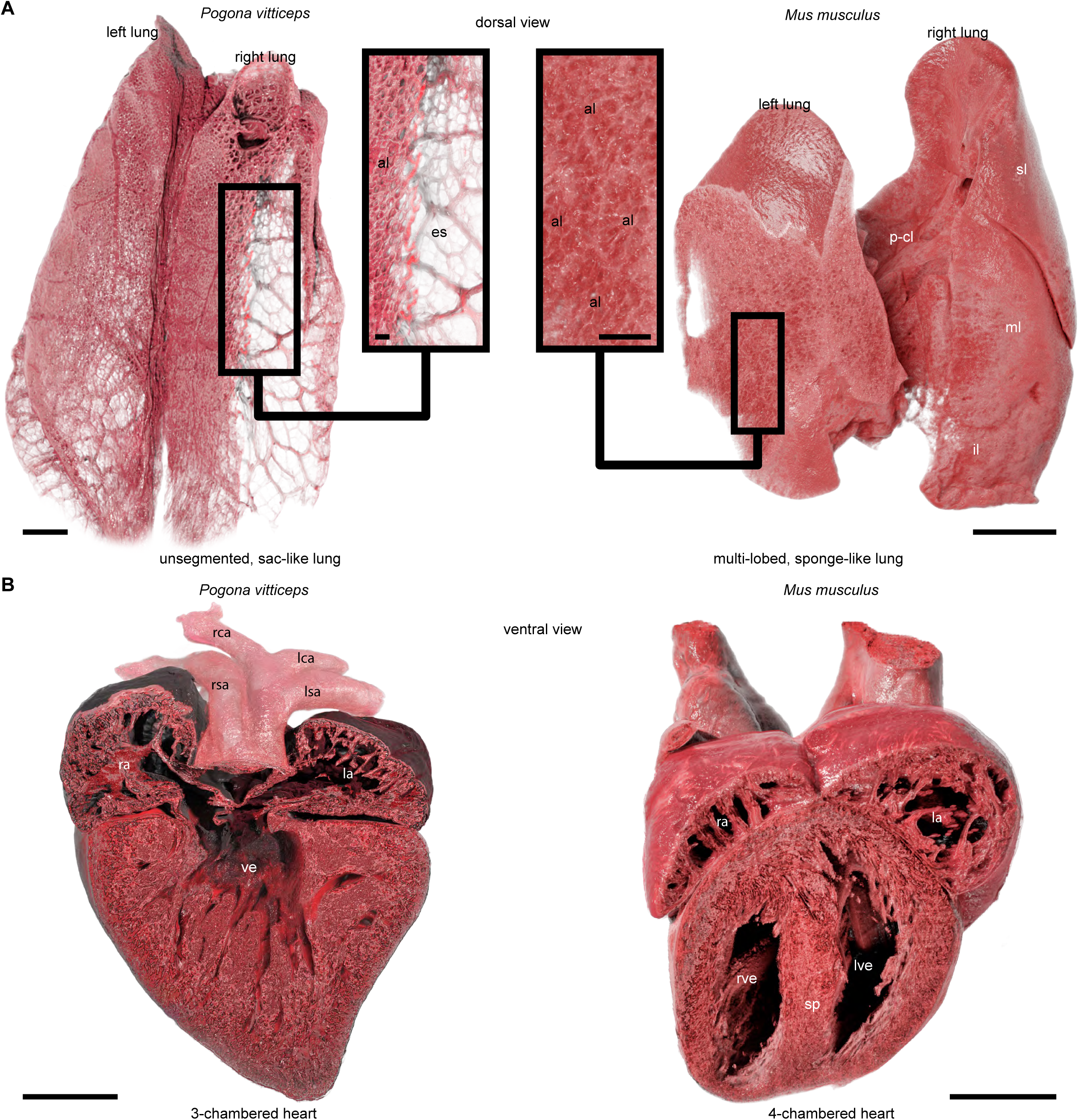
Comparative view of internal organs from different vertebrate groups enabled by the SmARTR pipeline. (A) Cinematic rendering of the lungs in *P. vitticeps* (left) and *M. musculus* (right). The lizard lungs are cut to highlight their unsegmented and saccular organization typical of squamate reptiles, while the mousés multi-lobed lungs are cut to display their spongiform nature. High magnifications reveal alveoli distributed only along the outer surface in lizards or alveoli also distributed in the deep parenchyma in mice. (B) Cinematic rendering of the heart and its main inflow and outflow vasculature in *P. vitticeps* (left panel) and *M. musculu*s (right). The lizard heart is cut to show the three-chambered structure typical of squamate reptiles, characterized by an incomplete septation of the ventricle. Conversely, the mouse heart is cut to reveal the four-chambered configuration characteristic of mammals, resulting from the complete division of the ventricle. Abbreviations: al, alveoli; es, empty space; il, inferior lobe; la, left atrium; lca, left carotid arch; lsa, left systemic arch; lve, left ventricle; ml, middle lobe; p-cl, post-caval lobe; ra, right atrium; rca, right carotid arch; rsa, right systemic arch; rve, right ventricle; sl, superior lobe; sp, septum; ve, ventricle. Scale bars, 1 mm or 0.2 mm (insets).

The enhanced cutaway views enabled by the SmARTR networks, along with their potential for expansion and customization to suit specific scenarios, could serve as a valuable resource for teaching and implementing life science courses. This includes clearer observations of unique and group-specific anatomical features, making it a powerful tool in teaching comparative anatomy and promoting biodiversity awareness among the general public (Figs. 5 and 6). Additionally, virtual dissections are easily performed using a personal computer equipped with the necessary hardware.

SmARTR-derived illustrations can also support tissue and organ dissection, as well as surgical procedure teaching. Although not yet approved for clinical use, medical professionals are increasingly recognizing the potential benefits of cinematic rendering. Despite some practical limitations, this technology significantly enhances diagnosis, disease management, treatment planning, and anatomy/surgical protocols^72,73,78,80,154–157^. Similarly, postmortem and 3D-rendering of animal organs, whether affected by specific diseases or not, can create a comprehensive database illustrating alterations in organ surface and inner compartments from different perspectives (Figs. 5 and 6). This approach would complement traditional postmortem dissection images displayed in animal care and veterinary medicine textbooks, offering a more detailed and versatile educational resource. In a similar manner, SmARTR cutaway renderings could complement macroscopic, histological, or modeling techniques in identifying, characterizing, and illustrating mutant phenotypes and pathologies in life sciences basic research.

### Limitations of the pipeline

Although the SmARTR pipeline is designed to accommodate various rendering scenarios, it cannot cover the entire range of potential visualizations for specific research or educational needs. However, the modular nature of the SmARTR networks allows for modifications, expansions, or combinations to achieve specific types of visualization. Additionally, the reliance of the path tracer on Compute Unified Device Architecture (CUDA) architecture limits the pipelinés use to systems equipped with NVIDIA GPUs. Furthermore, depending on the sample size, acquisition parameters (such as voxel size and rotation step), and the number of volumes shown simultaneously in the scene, manipulating the volume appearance can become challenging, even on machines with high-end compatible GPUs. This difficulty arises from the stochastic and computationally intensive nature of Monte Carlo iterations. In such cases, 3D data can be resampled to reduce the computational burden, though this may come at the expense of rendering quality. Striking the right balance between computational efficiency and visual fidelity is crucial in such scenarios.

## CONCLUDING REMARKS

As a whole, the SmARTR pipeline stands out as a flexible and innovative framework exhibiting a wide range of applications in biological sciences, including education, basic and veterinary research, diagnostics, and outreach. Its modular design, coupled with diverse highly customizable visualization scenarios from the SmARTR network prototypes within a free software environment, positions it as a valuable addition to the toolkits of many users, particularly researchers and scientists working with 3D imaging. Furthermore, the SmARTR pipeline opens up new avenues for interdisciplinary collaboration and creativity, facilitating the merging of disciplines such as art and research and the emerging of “Sci-Art” projects (Figure 7), which have been gaining momentum in recent years and offer unique opportunities for interdisciplinary collaboration, innovation, and public engagement with science. This echoes the masterworks in classical natural science and anatomical history by highly influential scientists such as Vesalius, Merian, Hooke, Haeckel, Boas, and Paulli, replacing the skilled hand of the dissector and illustrator by modern digital tools. Owing to its nature, the SmARTR pipeline is poised to benefit from ongoing advancements in information technology and data processing resources, including improvements in both hardware, with increasingly powerful GPUs, and software, through the continuous development of AI and machine learning algorithms. These advancements, including machine-learned automatic organ segmentation^158,159^ now being implemented in biological sciences^160^ and the creation of free tools allowing the integration of 3D reconstructions into interactive extended reality environments— either augmented, virtual, or mixed reality^161–163^—could be a crucial enhancement to the SmARTR visualization framework, pushing 3D imaging-derived biological data visualizations into a new era.

**Figure 7.**
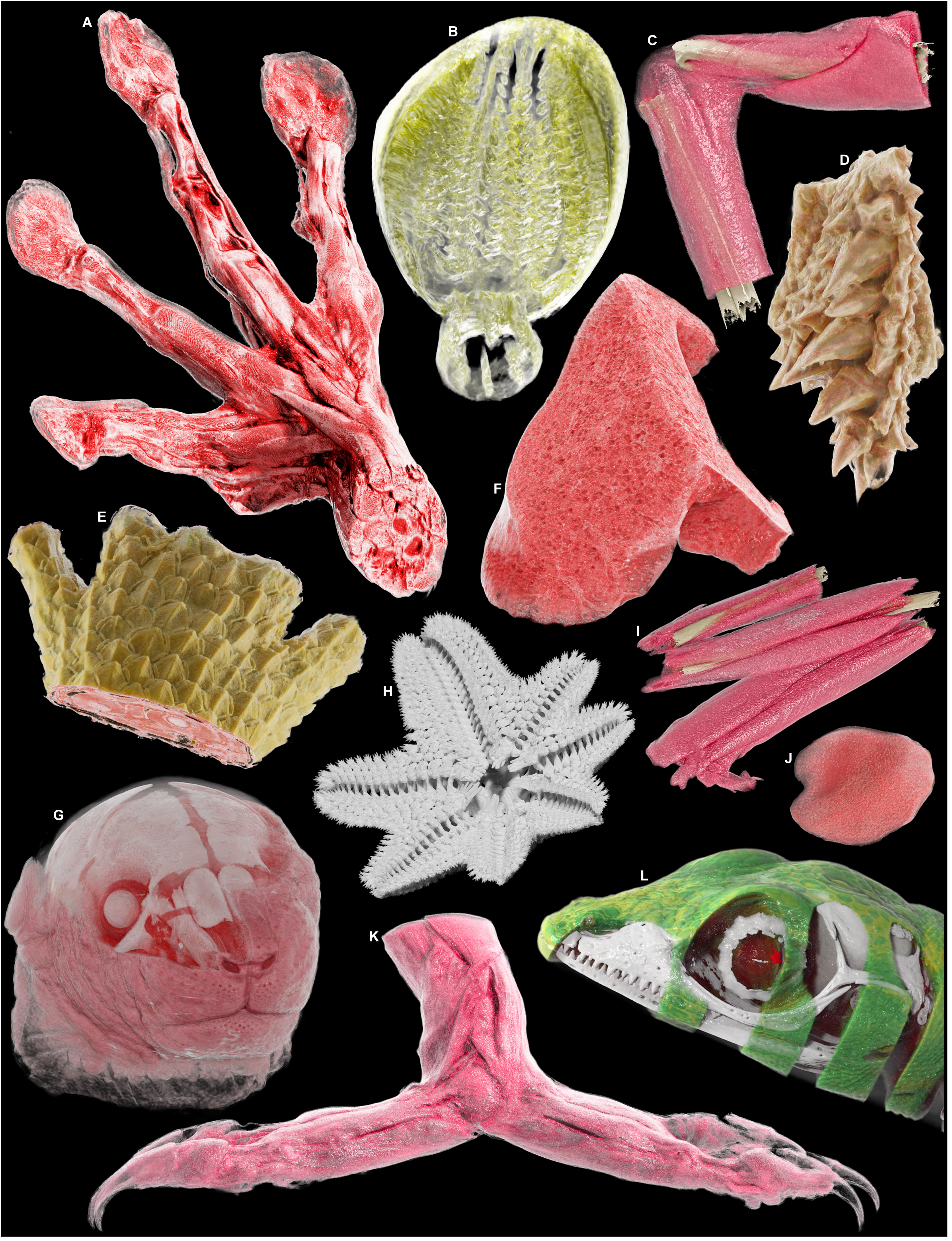
Artistic digital collage of various vertebrate and invertebrate anatomical parts processed with the SmARTR pipeline. (A) Forelimb musculature of *A. callidryas*. (B) Crop of *G. bimaculatus*. (C) Forearm musculature and bones of *A. callidryas*. (D) Head skin detail of *P. vitticeps*. (E) Carpal and metacarpal region of the forelimb of *P. vitticeps*. (F) Detailed view of the internal structure of the lung of *M. musculus*. (G) Semitransparent cutaway view of the head of *M. musculus* revealing cranial bones and internal soft tissues. (H) Skeleton of *A. burtoni* in ventral view. (I) Hindlimb musculature and bones of *A. callidryas*. (J) Tongue of *A.callidryas*. (K) Hindlimb musculature of *C. calyptratus*. (L) Semitransparent cutaway view of the head of *A. carolinensis* revealing cranial bones and the brain.

## METHODS

### Sample collection

Nine samples belonging to different vertebrate and invertebrate groups (Supplemental Table S1) were obtained from various sources, including animal colonies at the University of Helsinki (*Anolis carolinensis, Pogona vitticeps, Mus musculus*), private breeders (*Chameleo calyptratus*), specialized retailers (*Agalychnis callidryas*, *Gastromyzon zebrinus, Gryllus bimaculatus, Pterostichus oblongopunctatus*), and Sea Life Helsinki aquarium (*Aquilonastra burtoni*). Captive breedings and experiments at the University of Helsinki were approved by the Laboratory Animal Centre (LAC) of the University of Helsinki and the National Animal Experiment Board (ELLA) in Finland (license numbers ESAVI/7484/04.10.07/2016 and ESAVI/5416/2021).

### Sample preparation and staining

Euthanized or freshly naturally dead samples were placed in a large volume (15-20 times the sample volume) of 70% ethanol aqueous solution (*G. bimaculatus* and *P. oblongopunctatus*) or 4% paraformaldehyde (PFA) in phosphate buffered saline (PBS) solution (other species) for 24 hours. If needed, samples were repositioned on plastic slides using a flexible cyanoacrylate glue after 30 minutes of fixation. This process facilitates sample mounting prior to CT-scanning and minimizes the occurrence of unwanted sample movements during image acquisition. After fixation, samples were dehydrated through a series of 24/48-hour washes in solutions with increasing ethanol concentrations (30%, 50%, 70%), and kept in 70% ethanol prior either to be scanned to obtain hard tissue reconstructions or stained with suitable agents for soft tissue acquisition. Whole animal bodies were stained in 1% (w/v) solution of iodine dissolved in 70% ethanol (I_2_E_70_), while dissected parts/organs and the intact *A. burtoni* were incubated in 0.7% (w/v) solution of PTA dissolved in 70% ethanol (PTAE_70_; Supplemental Table S1). To minimize the number of animals used in the study, organs were dissected from previously iodine-stained sample bodies and further stained in 0.7% PTA solution whenever possible (Supplemental Table S1). Staining agent concentration and solvent selection was made taking into consideration literature data^37–39,41,55,146,164,165^ and author’s previous experience^33,47,126^ to maximize applicability of the protocol, owing to the high sample heterogeneity. Sample preparation and staining yielded excellent contrast across the wide range of sample types and scales processed in the study, while also minimizing tissue morphological alteration. In less broad research or educational contexts, however, sample- or tissue-specific optimized staining procedures, including methods such as perfusion rather than submersion, may be necessary to achieve ideal levels of tissue contrast.

### Sample mounting and µCT imaging

Maintaining adequate hydration is a critical consideration when performing µCT scans of biological samples. Dehydration during the scan can lead to tissues shrinkage and distortion, resulting in artifacts and potentially permanent damage to sample. Moreover, prolonged scanning times, such as those required to image whole small animal bodies, can further exacerbate this issue. To circumvent this problem, custom wet chambers were developed using standard laboratory equipment (Supplemental Figure S1). The following setup has been devised to simplify sample visualization process across a wide and diversified range of organisms and organs. Although it has proven effective in preserving sample position and minimizing sample shrinkage consistently, even in intact samples undergoing multiple and protracted scans (over 180 hours in total), any mounting strategy that best suits a particular specimen can be adopted prior to scanning. In our approach, following the dehydration steps, the slide holding the sample was attached by its short side to a cut pipette tip that had been glued to a suitable container cap. The container, either a 50 ml or 15 ml tube, was selected to be wider than the sample holder and had its closed end cut to fit the specimen’s length. Its internal walls were lined with absorbent paper, which extended beyond the tube’s height. Several drops of 70% ethanol were used to moisten the absorbent paper, ensuring it adhered to the walls. The sample was then carefully placed inside, and the tube cap secured. To maintain a humid environment, 70% ethanol was added to the container just below the level of the sample, and the edges of the absorbent paper protruding beyond the tube’s height were folded inwards to form a roof over the sample. Finally, the setup was sealed with Parafilm to prevent evaporation (Supplemental Figure S1, top panel). A similar setup was adopted for dissected sample mounting (Supplemental Figure S1, bottom panel), where smaller tubes, such as 1.5 ml, 2 ml, or 5 ml were used. Here, samples were glued to a plastic slide shaped to match the tube’s profile, positioning the sample centrally within the tube to allow adequate space around it. Humidity was preserved by placing strips of 70% ethanol-soaked paper inside the tube, along with a few drops of ethanol below the sample level. Once ready, samples were then scanned at the University of Helsinki imaging facility using Skyscan 1272 (Brucker, Belgium). The same filter type, source voltage, and source current were used for all scans (Al 0.25 mm, 60 kV, 166 μA, respectively). However, the voxel size, rotation steps, and average framing values varied depending on specimen size, ranging from 2.75 to 20 μm, 0.2 to 0.4 degrees, and 8 to 15 frames, respectively. Scans were reconstructed using NRecon 1.7.0.4 software (Bruker) and, when required, segmentation, resampling, and registration were performed using the software Amira 5.5.0 (Thermo Fisher Scientific, U.S.A.).

### MeVisLab network prototyping and generation of cinematic 3D renderings

Six main networks, each offering different visualization opportunities (SmARTR_Single_Volume, SmARTR_Multi-Mask_Volume, SmARTR_Multi-Independent_Volume, SmARTR_Nested_Multi-Volume, SmARTR_Pattern_Drawing and SmARTR_Advanced_Multi-Volume), along with an accessory one (SmARTR_Volume_Extraction), have been designed and prototyped in the platform MeVisLab (MeVis Medical Solutions AG and Germany and Institute Fraunhofer MEVIS, Bremen, Germany). The six prototyped network files, a description of each network, and a detailed step-by-step manual with practical examples to guide users in implementing cinematic rendering of their samples, are provided (Supplemental Methods). MeVisLab (download here: https://www.mevislab.de/download) is a versatile modular application framework compatible with both Mac and Window operating systems, featuring sophisticated algorithms that support detailed morphological and functional image processing and analysis. The free MeVisLab SDK version is distributed for non-commercial use and is designed for medical image processing. While limitations exist in this version, the prototyped networks rely exclusively on MeVisLab standard modules (Supplemental Methods), allowing them to be replicated, modified, improved, and utilized for scientific publications without any license requirements. One of the key components of MeVisLab, the MeVis Path Tracer, enables cinematic rendering of volumes, featuring physically based lighting with area lights and soft shadows, thanks to a Monte Carlo path tracing framework that operates on CUDA-enabled NVIDIA (NVIDIA Corporation, Santa Clara, CA, U.S.A.) GPUs. Path tracing supports interactive, photorealistic 3D environments that showcase dynamic lighting, along with caustics. However, it requires numerous iterations to achieve high-quality images, making it computationally intensive. The renderings in the article were implemented using an NVIDIA RTX A4000 GPU (with 6144 CUDA cores and 16 GB of dedicated random access memory, RAM) installed on a 64-bit Microsoft (Microsoft Corporation, Redmond, WA, U.S.A.) Windows 10 Enterprise workstation, equipped with a12th Gen Intel^R^ (Intel Corporate, Santa Clara, CA, U.S.A.) Core^TM^ I7-2700 and 64 GB of RAM. However, preliminary work on rendering and networks has been conducted on less performing machines featuring an NVIDIA GeForce GTX 750 (512 CUDA cores and 1GB of dedicated RAM).

### Image post-processing

If needed, adjustments to parameters like brightness, contrast, and exposure of the 2D images relative to the rendered volumes acquired in MeVisLab were made using Photoshop software, version 25.1 or later (Adobe Inc., Mountain View, CA, U.S.A.). Furthermore, to enhance realism, photographs of some specimens have been used to replace the eye region in a subset of the rendered species (*A. carolinensis, A. callidryas and C. calyptratus*). Finally, the “generative fill”—a generative artificial intelligence (AI) tool embedded in Photoshop from version 25.0—allows the creation of images based on user text prompts, even for commercial purposes, and was used to reproduce specieś natural habitats in some renderings.

## DATA AVAILABILITY

A step-by-step guide describing the MeVisLab visualization framework and detailing the implementation and use of the different SmARTR networks is provided as Supplemental Methods. Eight folders, each containing one or more relevant network files, the scan and mask files required for the practical examples detailed in the guide, and an additional folder with LUT presets, have been uploaded to Zenodo and are accessible <u>here</u>.

## ACKNOWLEDGEMENTS

We thank Heikki Suhonen (University of Helsinki, Finland) for access to X-ray computed tomography facilities; Jouni Jaakkola (Sea Life Helsinki) for marine specimens (*Aquilonastra burtoni*); Mari Joki, Peetri Joki, and Joni Ollonen for the chameleon sample; Ida-Maria Aalto and Joni Ollonen for SmARTR network testing; and members of the Di-Poï laboratory as well as Sascha Heckmann and Olaf Konrad (MeVisLab Core Team, MeVis Medical Solutions AG, Bremen) for helpful discussions. This work was supported by funds from the Academy of Finland (decision 356867 to N.D.-P.), the Jane and Aatos Erkko Foundation (to N.D.-P.), and the Sigrid Jusélius Foundation (to N.D.-P.).

## AUTHOR CONTRIBUTIONS

S.M. and N.D.-P. designed the overall experimental approach and selected the species sampling. S.M. developed the SmARTR pipeline, created the SmARTR networks, carried out micro-CT scans, and performed all 3D renderings. S.M. and N.D.-P. made the figures and wrote the paper. The two authors approved the final version of the manuscript.

## MATERIALS & CORRESPONDENCE

Correspondence and requests for materials should be addressed to S.M. and/or N.D.-P.

## SUPPLEMENTAL INFORMATION

Supplemental Methods

Supplemental Figure S1

Supplemental Table S1

