## Supplemental Information for "The SmARTR pipeline: a modular workflow for the cinematic rendering of 3D scientific imaging data"

#### Contents:

**Supplemental Methods**

**Supplemental Figure S1**

**Supplemental Table S1**

**Supplemental Methods**

### SmARTR step-by-step cinematic rendering workflow

#### Introduction

The steps to implement the cinematic rendering of 3D scientific data will be detailed in the following sections. The SmARTR visualization framework relies on six main modular networks, the **SmARTR networks,** which are prototyped in the free MeVisLab SDK (MeVis Medical Solutions AG and Fraunhofer MEVIS, Germany), an image processing platform distributed for non-commercial use.

The **SmARTR networks**—which have been uploaded on Zenodo and available here along with all the relevant files needed to replicate the examples detailed in this guide—cover a wide range of visualization scenarios and come into two forms:

- **Example networks**: These networks feature fully pre-configured modules designed to replicate the visualizations described in the various sections. The relevant volume files for these examples are provided in the "**Assets**" folders associated with each network.
- **Empty networks**: These fully functional networks include modules related to volume appearance set to basic (not default) values, which users must modify according to their needs. Ideally, after selecting a suitable network for their visualization needs (see below), users can load their samples, adjust the volume appearance, perform any necessary volume cutting operations, and acquire high-quality images without worrying about the underlying network construction or module configurations for specific visualizations.

However, the purpose of the network sections is not only to provide the basic information on single module configuration but also to illustrate the concepts behind module integration. These principles are fundamental to the specific visualizations each network offers. By understanding these concepts, users can customize, improve, or create new networks that better fit their needs.

##### General considerations about section content and network selection

Though presented with a specific example, the applications of each **SmARTR** network are broad, as illustrated by the **SmARTR_Advanced_Multi-Volume** network which combines concepts individually elaborated in other networks. Below is a list of considerations for the network sections, designed to help users select the network that best fits their visualization needs.

- **Section 1** (pages 3-5) Gives a brief overview of MeVisLab interface and behavior.
- **Section 2** (pages 6-14) Describes the **SmARTR_Single_Volume** network, which applies to the rendering of a single, homogeneous, and non-segmented volume, like a single organ or animal part. "Homogeneous" refers here to both the exterior and interior composition of the volume, which can be well approximated by a single appearance profile (color, light interaction, etc.). This ensures that rendering will still accurately represent the volume´s natural aspect even if volume cutting is performed. In the case of a single volume with a non-homogeneous structure, the **SmARTR_Nested Multi-Volume** network can be selected to obtain photorealistic volume cutaway views (see below for more details). **Section 2** also introduces the majority of the modules used in the various networks and details their configuration. Thus, it may also serve as a starting point independently from the specific network description.
- **Sections 3 and 4** (pages 15-20 and 21-25, respectively) Present two methods to render multiple subvolumes deriving from the same scan volume—such as the multiple lizard internal organs in the example—in the same 3D scene. Both networks utilize segmentation masks, either to "label" the corresponding subvolumes on the whole scan volume, or to extract these subvolumes from the whole scan volume, representing them as independent structures. The first strategy, implemented in the **SmARTR_Multi-Mask_Volume** network (**Section 3**), relies on the possibility to generate multiple instances (copies) of the whole scan volume, each displaying only the subvolume specified by the

corresponding segmentation mask. The configuration of the three modules (the "**SopathTracerVolume**", the "**SopathTracerMaskVolume**" and the "**SoPathTracerVolumeInstance**") required to implement such an approach—which serves as the foundation of other multi-volume networks described later in the text (**Sections** 5, 6 and 7)—is detailed for the first time in this section. The multi-volume rendering strategy proposed in **Section 4** relies, instead, on an accessory and separate network (the **SmARTR_Volume_Extraction** network) to extract the subvolumes of interest from the whole scan volume. These extracted subvolumes are then rendered as individual structures in the **SmARTR_Multi-Independent_Volume** network. The subvolume extraction procedure described in **Section 4** is also an intermediate step in the realization of the **SmARTR_Advanced_Multi-Volume** network renderings (**Section 7**), which also relies on concepts introduced for the first time in **Section 4.2,** step 3*.

- **Section 5** (pages 26-30) illustrates how to achieve photorealistic cutaway views when dealing with volumes nested one within the other, such as the head with its skull and brain, or the body and its visceral organs, using the **SmARTR_Nested_Multi-Volume** network. This network can be selected also when generating cuts through a single non-homogeneous volume (see **Section 2** bullet, above). Moreover, it can be chosen also to provide an additional level of refinement when performing volume cutting operations on a single homogeneous volume. The network takes advantage of the binary mask generated upon volume cutting (see steps 3ab, **Section 2.2**) to automatically create an additional volume superimposed to the outermost volume surface exposed by volume cutting. This new volume can be independently rendered to increase scene realism.
- **Section 6** (pages 31-36) describes how to create color patterns on volumes to approximate natural coloration of samples using the **SmARTR_Pattern_Drawing** network. While complex color schemes are unattainable, as they are typically the prerogative of methodologies other than internal imaging and DVR techniques, the network allows multiple patterns to be generated and rendered as independent structures, enhancing the realism and lifelikeness of the samples. Furthermore, the binary masks generated during pattern drawing can be utilized in the realization of enhanced cutaway views (**Section 7**).
- **Section 7** (pages 37-41) details the **SmARTR_Advanced_Multi-Volume** network, the end point of a workflow involving multiple networks, which also exploits the concepts illustrated in step 3* of **Section 4.2**. The network is a modified version of the **SmARTR_Nested_Multi-Volume** network (**Section 5**) and allows enhanced cutaway views of multiple volumes that share the same internal structure but differ in surface color, such as the atrial and ventricular heart regions or the differently colored parts of a lizard´s body. The binary masks obtained either with the **SmARTR_Pattern_Drawing** network (**Section 6**) or by segmenting the volumes of interest in external software, are used to extract the relative subvolumes in the **SmARTR_Volume_Extraction** network (**Section 4**) which are then loaded, together with the volume they derive from, in the **SmARTR_Advanced_Multi-Volume** network, where multi-volume cutaway views can be created.

### Section 1. MevisLab visualization environment

#### 1.1 Brief overview of MeVisLab interface and behavior

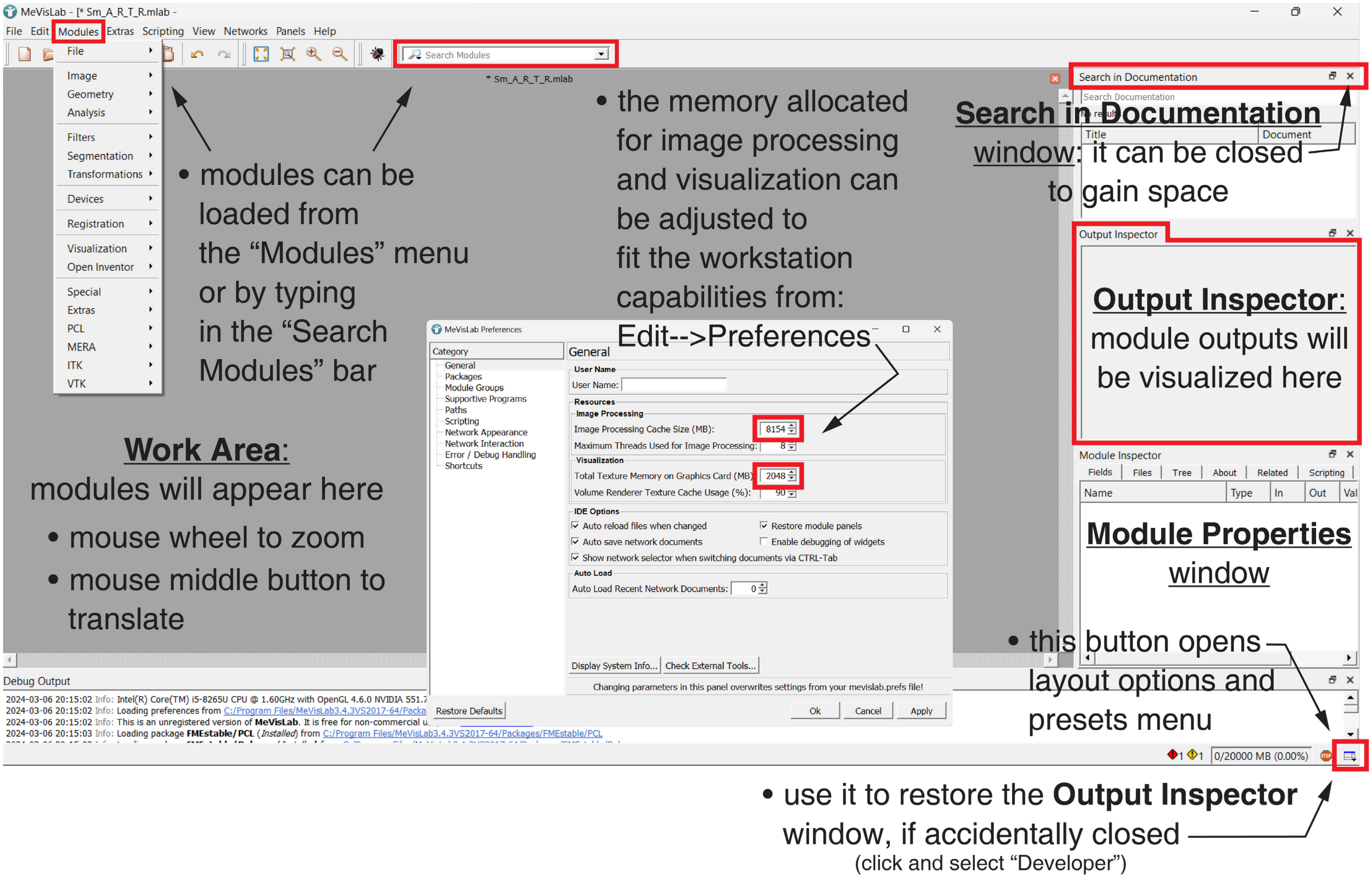


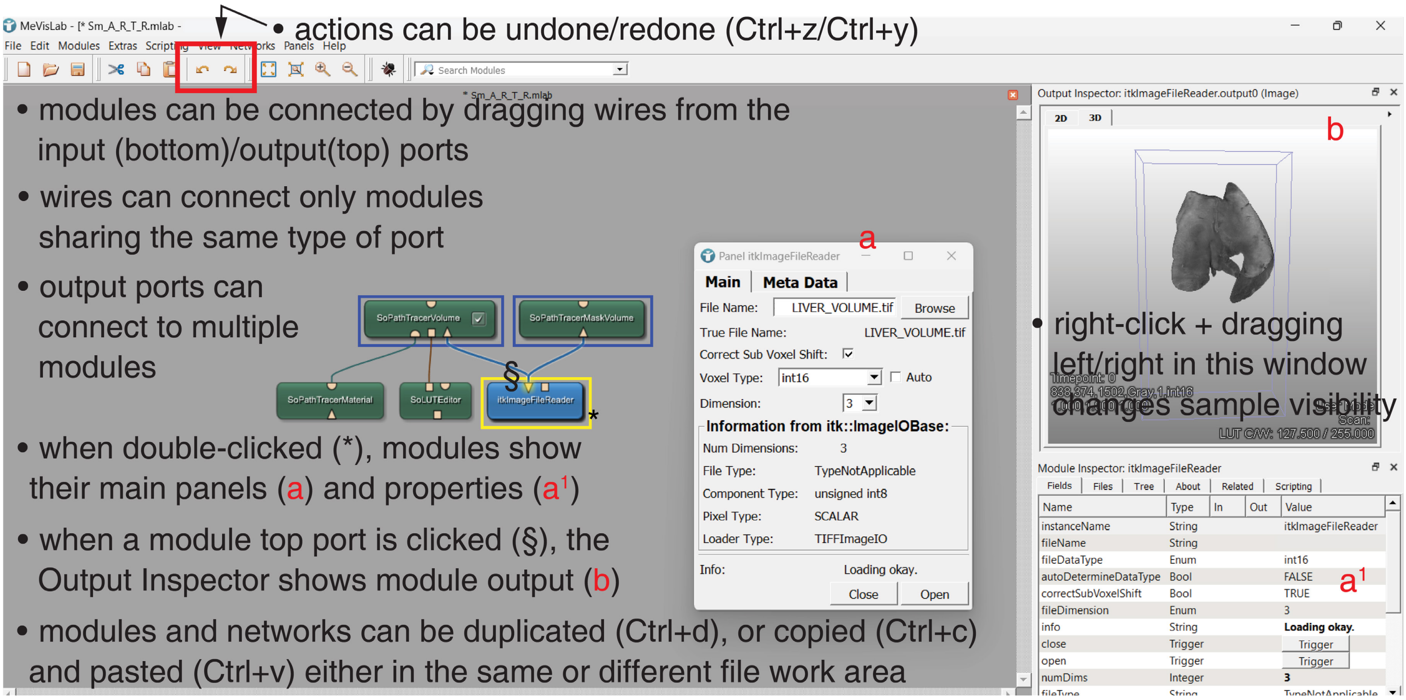

- modules can be automatically rearranged by selecting them and clicking Ctrl+1, and grouped by right-clicking and selecting Grouping-->Add to new group

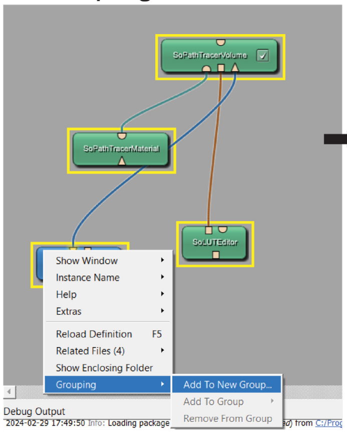


- modules can be renamed by right-clicking and selecting Instance name-->Edit Instance name

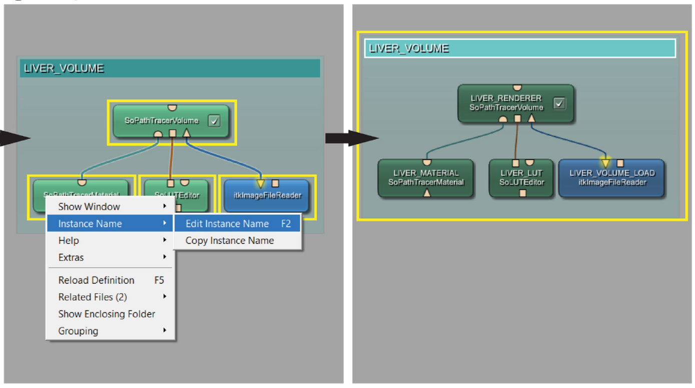


- some modules will display additional ports to allow multiple connections

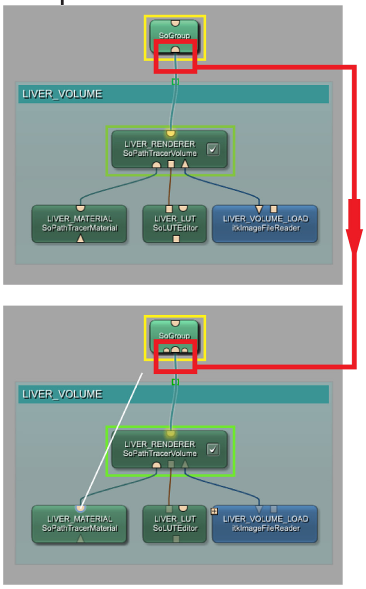


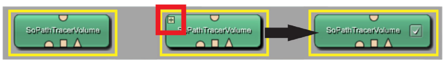


- modules can be be collapsed/expanded by hovering on their top left corner and clicking on the + or - button

#### 1.2 Interaction viewer (SoExaminerViewer) features

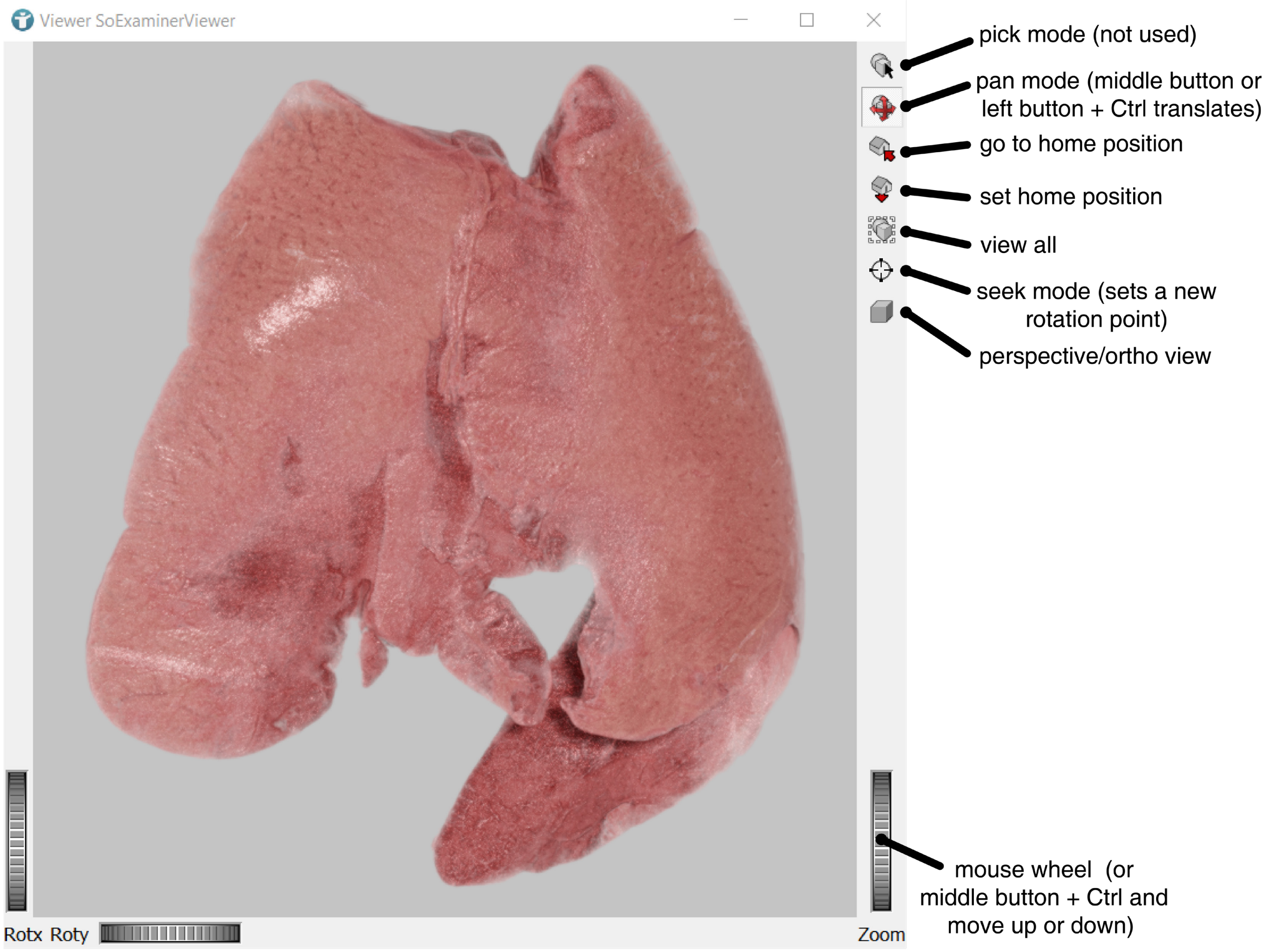


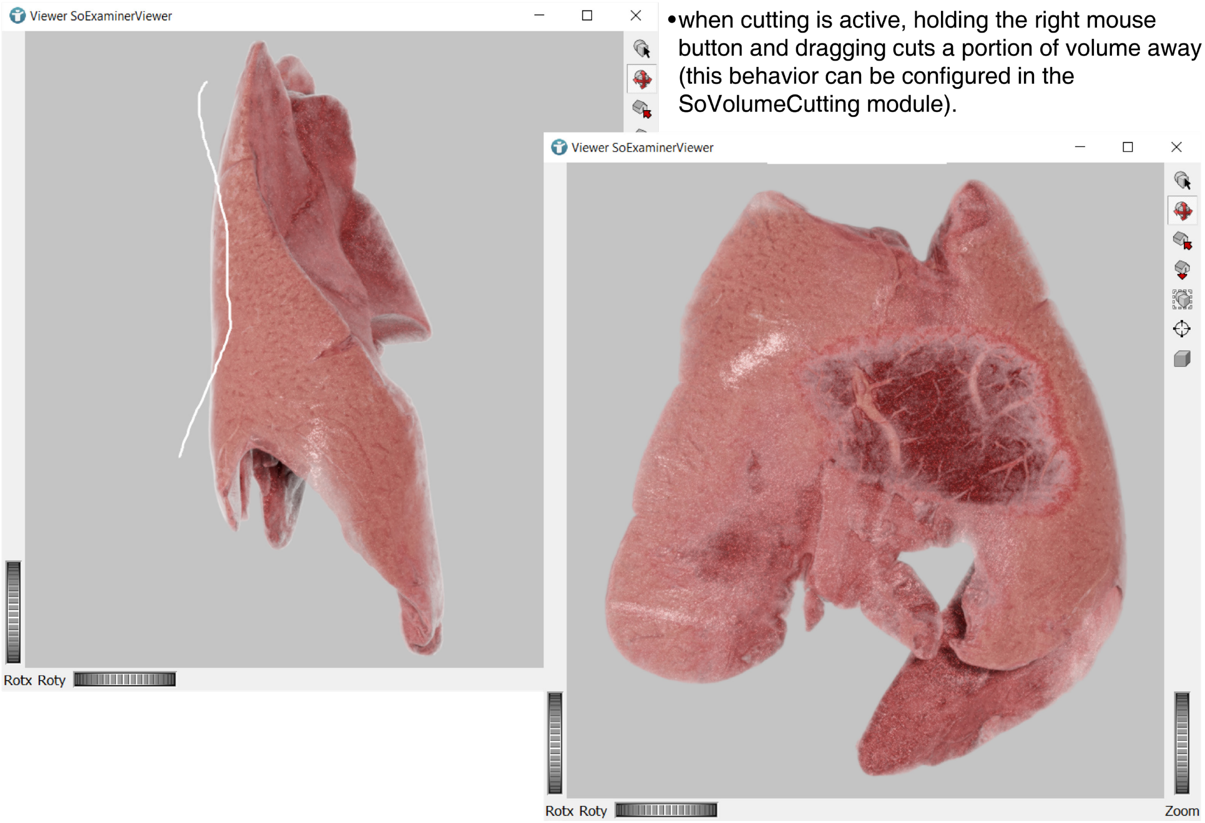

#### Section 2. Rendering of single, homogeneous, and non-segmented volume

This section illustrates the features and configurations of most MeVisLab modules used in the all the prototyped networks. Indeed, though presenting some distinctive modules or module configurations/integrations, the multi-volume networks detailed later in the text are mainly extensions or modifications of the one described in this section.

The rendering of isolated, homogeneous, animal organ/part simply requires the sample scan data—usually in the form of 2D image sequences (.tif or other format)—to be loaded in the provided **SmARTR_Single_Volume** network file (.mlab) and to perform the required steps to modify volume appearance. Briefly, the 2D image sequence is loaded and the corresponding 3D volume created and optionally cropped (to save memory) by the modules in the "**Volume loading and cropping**" group. The volume is, then, fed to the "**Volume appearance**" and, optionally, to the "**Volume cutting**" blocks. Several parameters can be modified in the "**Volume appearance**" modules, each affecting volume representation. In particular, the "**SoLUTEditor**" module allows to set the color and opacity of the volume pixel groups while the "**SoPathTracerMaterial**" properties affect the way the volume surface interacts with light. The effect of parameter changes on volume appearance can be monitored in the "**SoExaminerViewer**" (Interaction viewer), accessed by double-clicking on the corresponding module balloon. The "**SoExaminerViewer**" is also the place where volume cutaways can be performed, according to the "**SoVolumeCutting**" module settings. Modules in the "**Light**" group are used to illuminate the scene and their intensity and orientation can dramatically influence volume appearance. Finally, the "**Image acquisition**" group modules allow to save high quality images of the rendered volume.

##### 2.1 The SmARTR_Single_Volume network overview

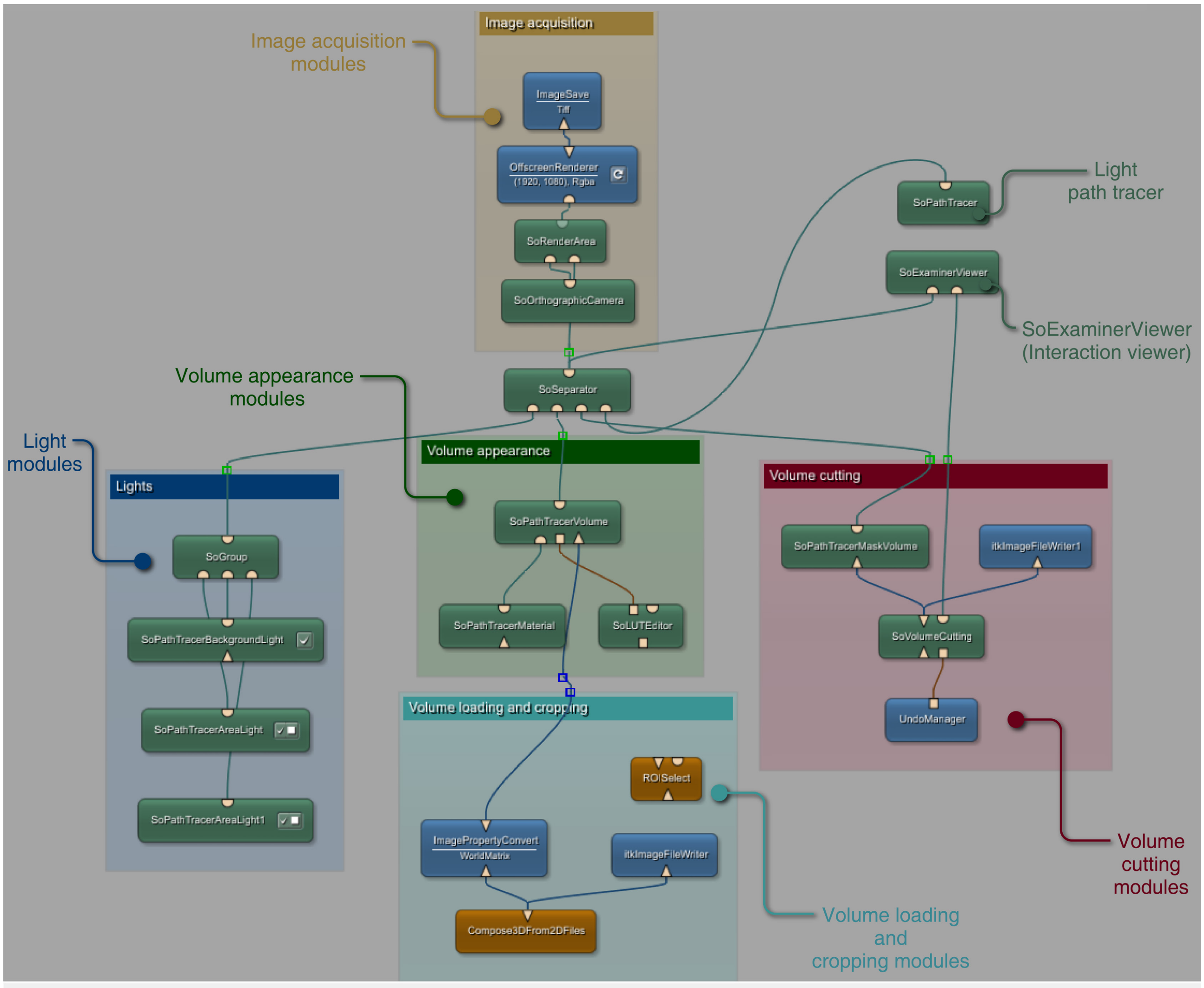

#### 2.2 Step-by-step protocol for single volume rendering and acquisition

##### 1. Load sample (Volume loading and cropping block)

a. Double-click on the "**Compose3DFrom2DFiles**" module to open the settings panel. To load a series of files as a volume, browse to the folder containing the image sequence (be sure that no extra image files, except those belonging to the scan stack, are in the folder), specify the file type in the "**Search Pattern**" field (*.tif, in the example) and hit "**Create 3D**". The "**Status**" field at the bottom of the panel shows the progression in 3D volume reconstruction.

b. Once the 3D volume has been created, open the "**ImagePropertyConvert**" panel and check that the "**X**", "**Y**", and "**Z**" values, in the "**Voxel Size**" field, are set to 1. If not, set them to 1 and click "**Apply**". Specifying the actual scan voxel size is not relevant for visualization purposes. If loading a single 3D volume file (e.g., 3d.tif, .nii, .mha, .nrrd), rather than a 2D image sequence, follow step d3, below.

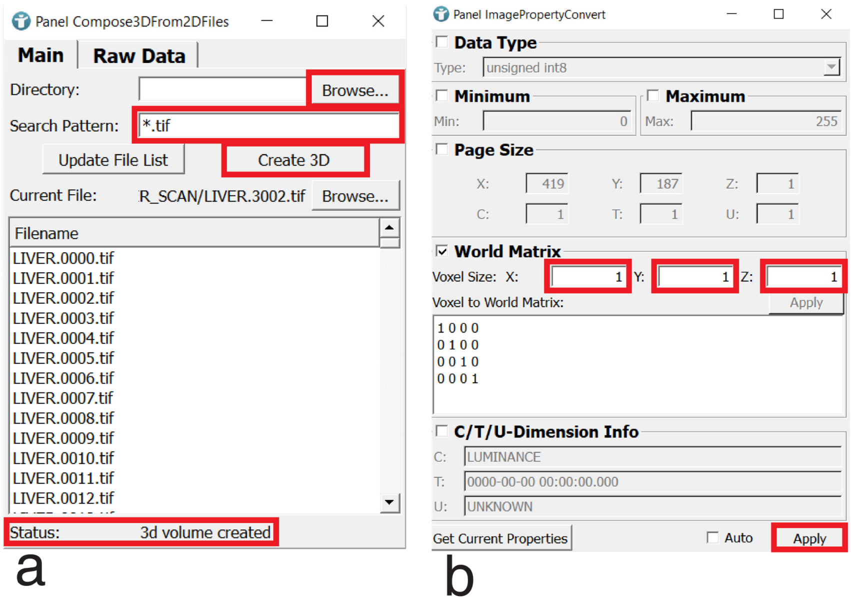


c. **(OPTIONAL)** If needed, the volume can optionally be cropped by connecting the "**ROISelect**" module to the network. (c1) Simply drag it on the wire linking the **ImagePropertyConvert**" and the "**SoPathTracerVolume**" module (both "**ROISelect**" input and output triangular ports will light up and automatically connect upon mouse button release). (c2) In the "**ROISelect**" panel the volume bounding box can be interactively resized, lowering GPU computational burden. Select the "**Navigate**" mode to scroll the slides by clicking the left mouse button, and hold "**Shift**" (cursor will turn to a crosshair) to drag the box sides till they fit the volume. (c3) Real time modifications of the bounding box can be also monitored in the "**3D**" tab of the "**Output Inspector**", after clicking on the "**ROISelect**" triangular output port.

d. **(OPTIONAL)** The reconstructed volume can be saved as a single 3D file (e.g., 3D tiff). (d1) Connect the "**itkImageFileWriter**" module either to the "**Compose3DFrom2DFiles**" (to save the original volume) or to the “**ROISelect**” (to save a cropped version of the volume) output port. (d2) Simply specify the filename, followed by the chosen file type suffix, in the module panel and click “**Save**”. (d3) To load the file, connect an “**itkImageFileReader**” module (type “reader” in the “**Search Modules**” box to find it) to the “**ROISelect**” input port. If volume cropping is not necessary, bypass the “**ROISelect**” module and connect the “**itkImageFileReader**” directly to the “**SoPathTracerVolume**”. Browse to file location, tick the “**Auto**” checkbox and “**Open**” the file.

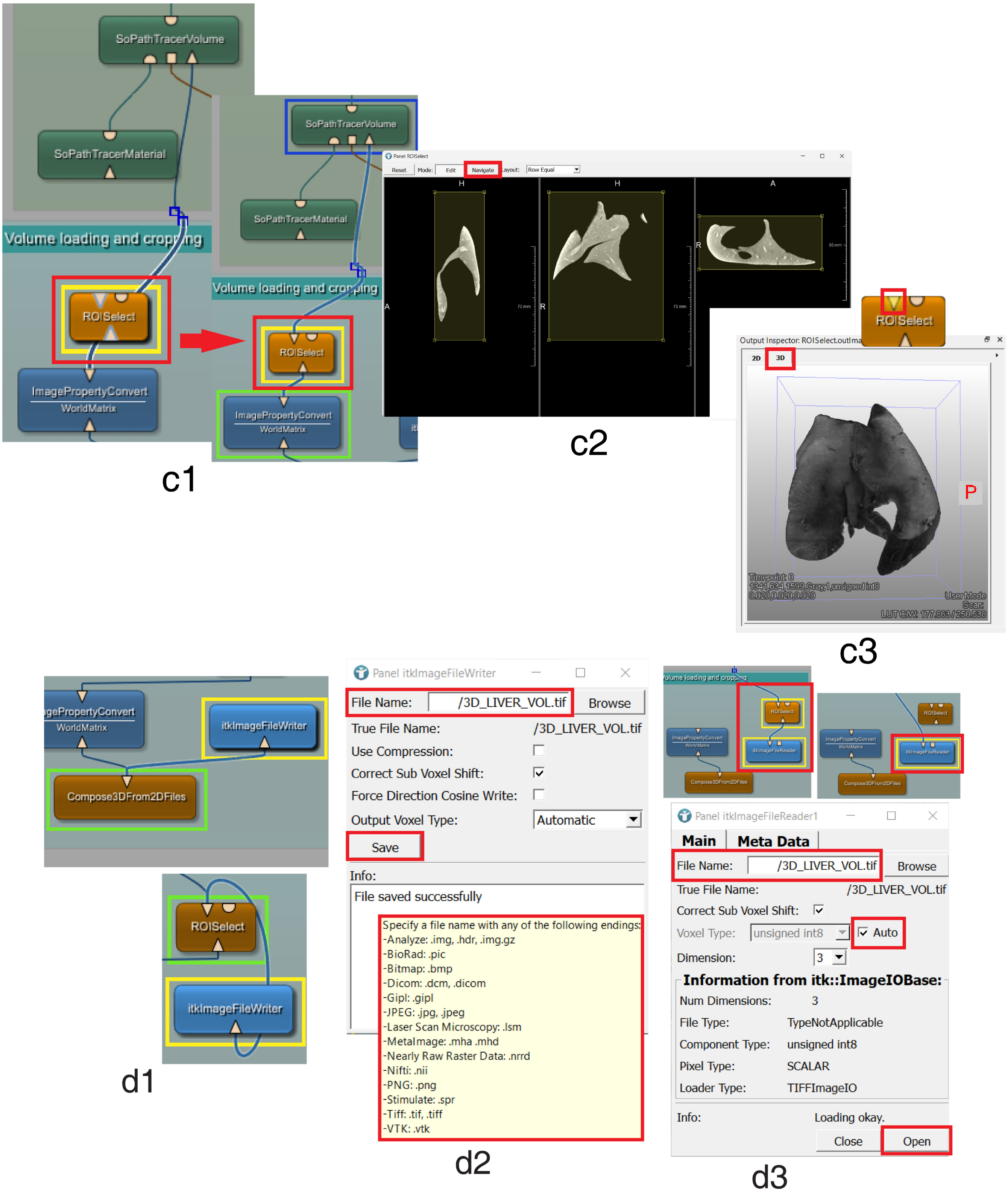

2. **Visualize and modify volume appearance (Volume appearance and Lights blocks, SoPathTracer)**

a. Double-click on the “**SoExaminerViewer”** and hit the “**View all**” (see **section 1.2**) button to visualize the volume. The viewer allows to monitor how parameter modifications in the “**SoPathTracer**”, “**Volume appearance**”, and “**Light**” block modules affect the volume look. “**SoExaminerViewer”** background color (see also next step) and other settings, can be modified in the module panel, accessible by right-clicking on the module balloon and selecting “**Show Window à Panel**”.

b. The “**SoPathTracer**” is the module responsible for MeVis Path Tracer visualization framework. Several parameters such as exposure, denoising, sharpness, camera aperture and background colors can be set, influencing both real time volume rendering and image acquisition (see also step 4). In most cases, the default parameters work well for real time rendering (provided that the “**Denoise**”, “**Denoise Initial**”, and “**Denoise Final**” boxes are unchecked in the module panel “**Resolve**” tab, see **figure b**, bottom). In this phase, modifications to default parameters might, thus, be limited to the background color to (e.g., to have an idea of how rendered volume will appear against a white background, like that of an article/book page) and/or to the “**Exposure**” value. The scene background color can be set by changing the “**Top color**” and “**Bottom color**” in the “**Main**” tab and selecting the preferred “**Blend Mode**”. The color of the “**SoExaminerViewer”** background also influences the scene background. With the “**SoExaminerViewer”** background color set to white/black, by selecting “**Blend**” as “**Blend Mode**” in the “**SoPathTracer**”, the scene background will be set to uniform white/black. The number of iterations used to estimate light path can be set in the “**Stop at Iteration**” field in the “**Main**” tab. This value, set to 1000 by default, can be raised (to 2500-5000) to achieve a better rendering quality (though, after a certain threshold, no relevant improvements can be detected by naked eye). Volume appearance can also improve by increasing the number of light bounces to be traced (1-16) in the “**Num Bounces**” box, though in this case the effects on rendering quality are sample-specific (most of the renderings displayed in the article and in this section have been achieved with the default “**Num Bounces**” value). Moreover, while higher iteration values do not impact real time rendering performance, increasing the number of bounces does. Indeed, when “**Num Bounces**” values are >1, path tracing estimation includes light scattering calculation, thus prolonging the time needed for an iteration to complete. In certain cases, the “**Aperture**” value can be modified to acquire images with depth of field effect to add realism to the scene (see step 4b). The “**Current Iteration**” field displays the iteration progression in real time. In the “**Resolve**” tab, together with a suitable “**Exposure**” value, one of the four available “**Tone**

**Mapping**” presets can be selected to better reproduce sample color shades. “**Sharpness**” can be set to 1, while default values can be used in the “**Sharpness Start**” and “**Sharpness blend**” fields.

c. Modify volume appearance by opening the “**SoLUTEditor”** panel and adjusting pixel range opacity and color in the “**Editor**” tab. A large variety of “**SoLUTEditor**” presets, can be found in the “**LUT_Presets**” file. The modules in the file can be copy-pasted into any network. Though possible, it is advised not to connect them directly to the network, as it is often simpler to start by editing a default color profile (press the 'Reset' button to revert to the default status, if needed). Instead, these modules can be used as guidance in color selection during the creation of new sample color representations. To edit a LUT profile, simply click on the black line to add new points and assign them a color using the color wheel at the bottom of the panel. Existing points’ position and opacity can be modified by dragging them. The opacity of each point corresponds to its height in the “**Editor**”. Points at the top have the maximum opacity while one at the bottom is transparent. The color interpolation between the various points is displayed by the bar below the graph. The LUT **“Alpha Factor”** (the sample global transparency) value (default =1), found in the “**Settings**” tab, can be lowered/raised to give a more vaporous/compact representation of sample texture.

- **(*) NOTE:** The “**SoLUTEditor**” has the “**Relative LUT**” box, found in the “**Settings**” tab, unchecked by default. This makes it difficult to edit volume colors. When creating new instances of the “**SoLUTEditor**” module, remember to flag the “**Relative LUT**” checkbox.

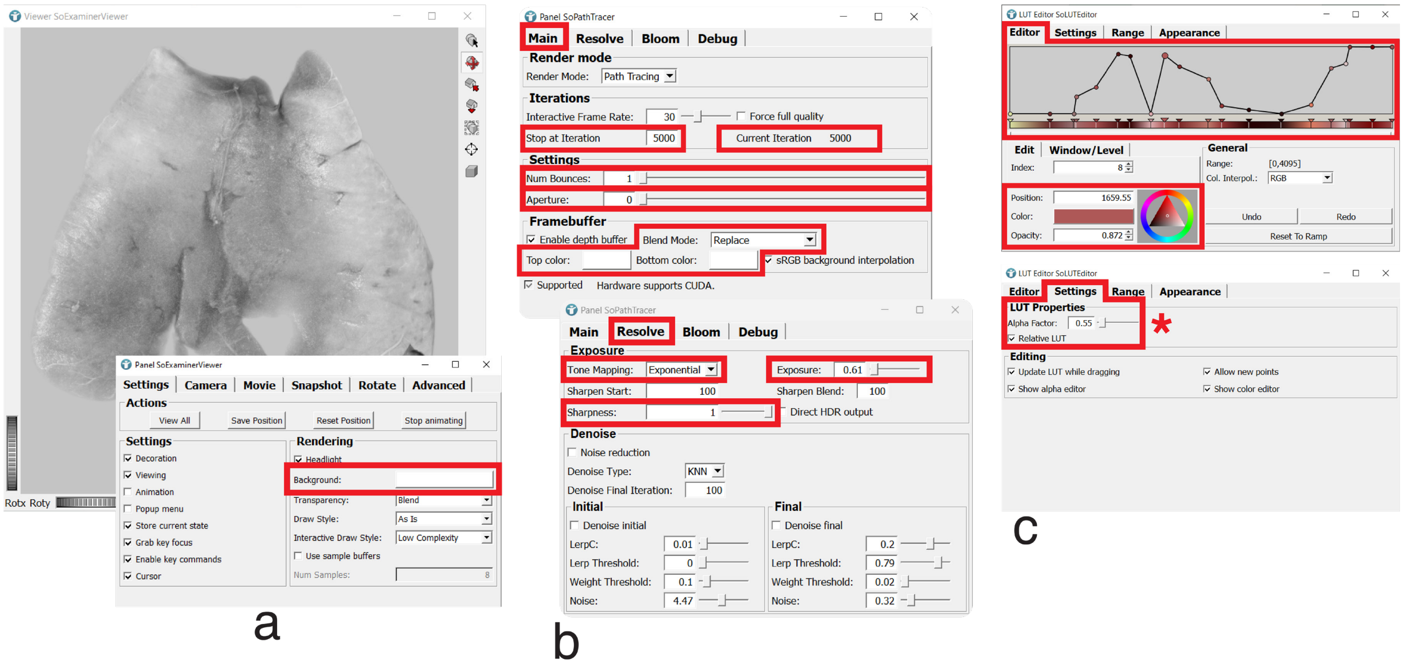


d. Modify light intensity and position in the “**SoPathTracerAreaLight**” module panel. “**Distance**”, “**Altitude**” and “**Azimuth**” can be set at the bottom of the panel by manipulating sliders and knobs to change light source coordinates. Typical “**Intensity**” value range is 2-15 while “**Distance**” can be set to illuminate larger (10) or smaller (0) volume areas. Other parameters, unless needed, can be left with their default values. Each light can be linked to the camera view by flagging the "**Attach light to camera**" box. A "**SoPathTracerBackgroundLight**" light can be added to retro-illuminate the sample. This module allows setting a top, middle, and bottom light color. New instances of the module come with the "**Render Source**" option active, which affects background color. Depending on visualization needs it can be deactivated. The module also features the "**Attach light to camera**" option.

e. The "**SoPathTracerMaterial**" module has four "**Material**" presets, accessible under the "**Surface Brdf**" tab. Each preset allows fine-tuning of volume surface attributes to add realism to the sample. The "**Principled**" preset worked best for most samples (typical settings are: "**Metallic**": 0-0.2; "**Specular**": 0.3-0.6; "**Roughness**":0-0.1, "**Specular Tint**": 0.2-0.6; "**Sheen Tint**": 0.5-0.8; "**Sheen**": 0.5-0.8; "**Clearcoat**":0.5-1; "**Clearcoat Gloss**" :0.5-1; "**Subsurface**": 0,8-1. Under the "**Volume Shader**" tab, tick the "**Enable volume shader**" box and select the "**Shader Type**". The "**Hybrid**" shader, with a "**Gradient Factor**" set in the range of 30-150 is effective for most types of samples.

f. Modify the "**Step Size Factor**", "**Step Size Factor Shadows**", and "**Self Shadowing Offset**" values in the "**SoPathTracerVolume**" panel "**Settings**" tab, by acting on the sliders. These parameters affect the volume rendering and shadowing properties. A "**Volume Name**" can be set to avoid confusion in multi-volume scenes. Switch to the "**Tag/MaskVolume**" tab (if not visible in the panel, click on the black arrowheads in the panel top-right corner). Set a name (liver, in the example) in the "**Mask Volume Name**" field. This step is necessary to enable volume cutaways using the "**SoVolumeCutting**" module (see next **Section**).

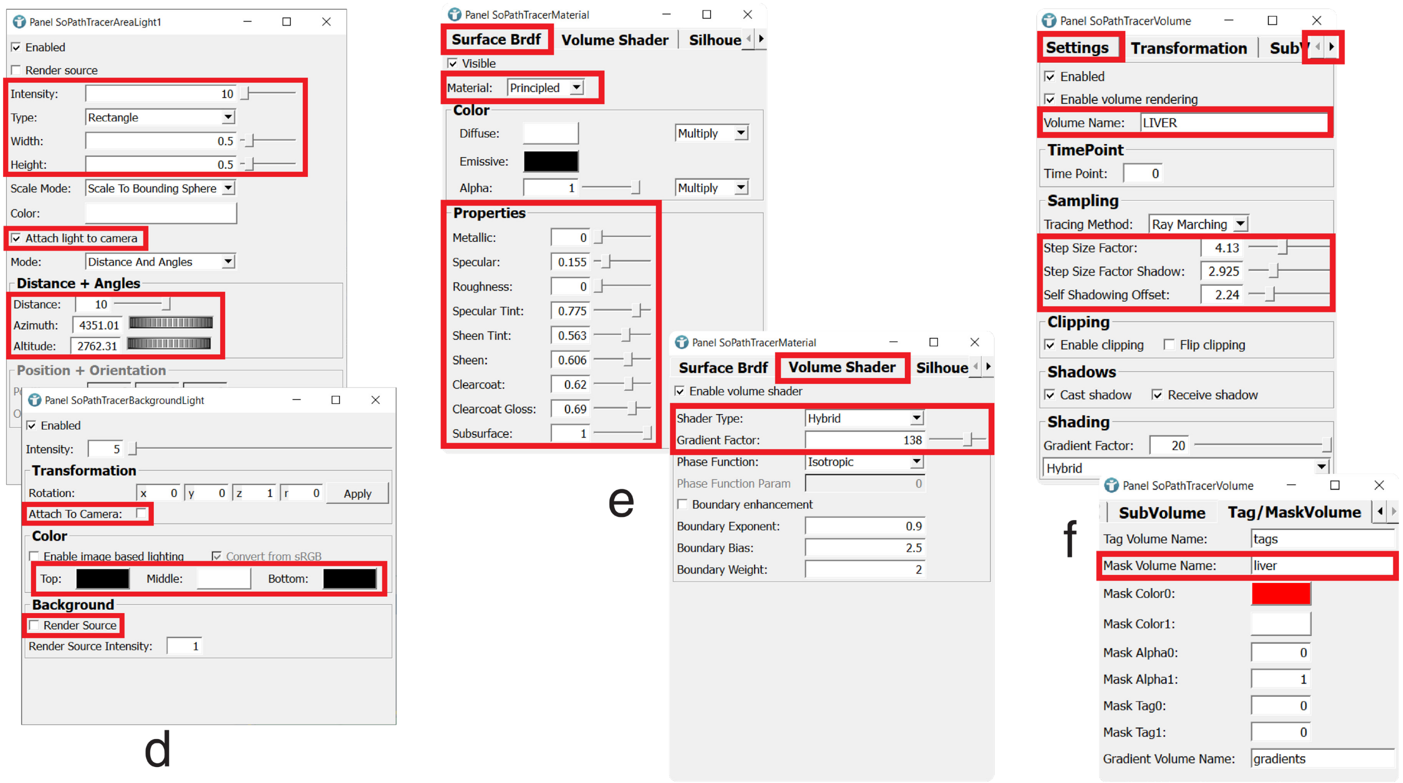

#### 3. Volume cutting (Volume cutting block)

The modules in the Volume Cutting block allow to interactively remove volume portions inside or outside a free hand contour drawn in the "**SoExaminerViewer**" (see **section 1.2**). Before starting the volume cutting operations, connect the output of the "**ImagePropertyConvert**", "**itkImageFileReader**", or "**RoiSelect**" module (depending on whether the volume has been cropped; see step 1d in **Section 2.2**) to the triangular input port of the "**SoVolumeCutting**" module.

a. Open the "**SoVolumeCutting**" panel to configure the module. Check the "**Interior**" or "**Exterior**" radio button to select the cutting behavior, i.e., whether volume parts located inside or outside the free hand contour drawn in the "**SoExaminerViewer**" will be removed (see **section 1.2)**. The options in the "**Add Mode**" either make a new cutaway replace the old one ("**Add with Shift**") or add it to the existing one/ones ("**Always Add**"). It is possible to set the color ("**Line Color**") and the width ("**Line Width**") of the contour line as well as which mouse button to use for cutting. Volume integrity can be restored by pressing the "**Reset**" button. Undoing/redoing the last action(s) can, instead, be performed, in the "**UndoManager**" module panel by clicking the "**Undo**"/"**Redo**" button.

b. To visualize the cutaways, it is necessary to feed the output of the "**SoVolumeCutting**" to a "**SoPathTracerMaskVolume**". In this module, the name set in the "**Volume Name**" field must match the one in the "**Mask Volume Name**" field of the "**SoPathTracerVolume**" (see **Section 2.2**, step 2f). Upon contour drawing, a binary mask is created and applied to the referenced volume by the "**SoPathTracerMaskVolume**" module output relayed to the "**SoSeparator**". Such mask features a value of "0" (black color) for cut parts and a value of 255 (white color) for the rest of the volume, letting only the latter to be visible.

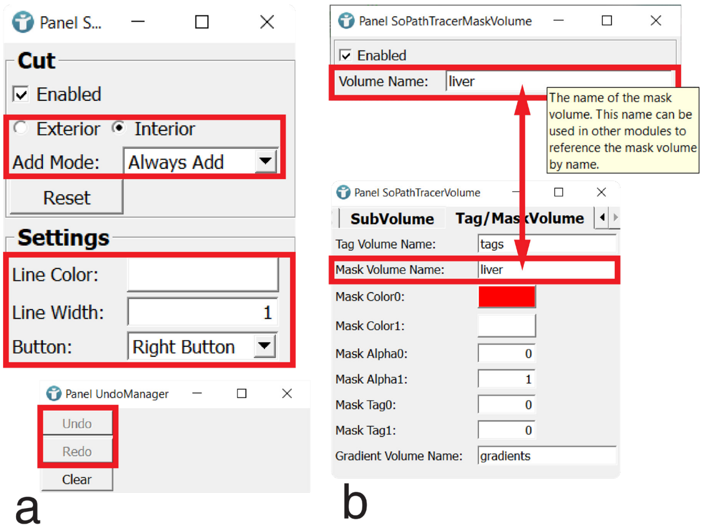


c. **(OPTIONAL)** The binary mask created during volume cutting is not stored in the modules so, upon file exit, any information about the cutaway view is lost. (c1) The mask can be saved in a suitable format (e.g., 3D tiff) via an "**itkImageFileWriter**" module connected to the "**SoVolumeCutting**" module. (c2) To load the file, connect an "**itkImageFileReader**" module directly to the

"**SoPathTracerMaskVolume**" input port, browse to file location, flag the "**Auto**" box, and open the saved mask file. (c3) This will restore the cutaway view in the "**SoExaminerViewer**".

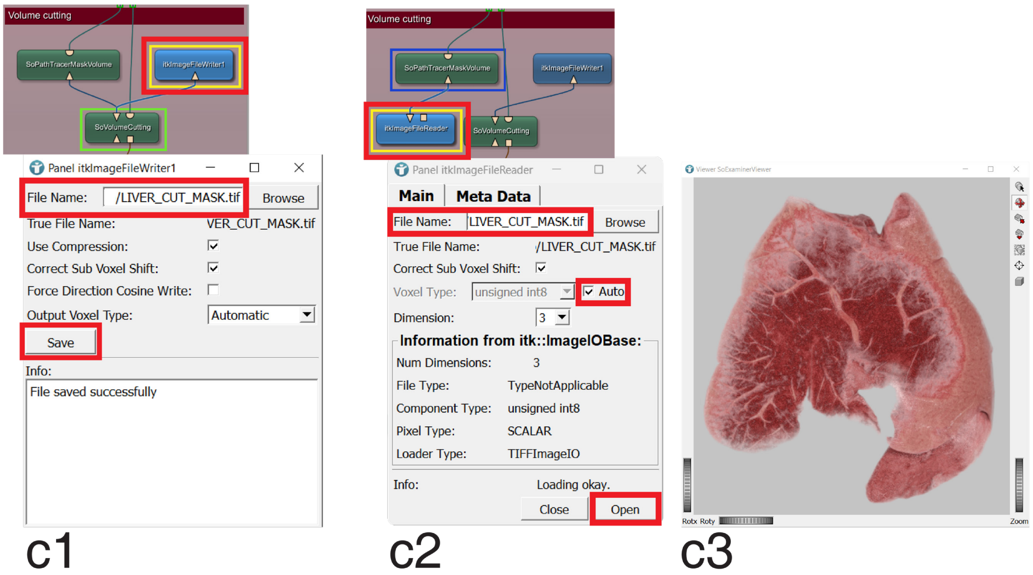


4. **Image acquisition (Image acquisition block)**

The easiest way to save an image of the rendered volume is to take a screenshot of the "**SoExaminerViewer**" by pressing F11. The snapshot will be automatically saved in MeVisLab "**Screenshot gallery**", accessible via the "**Layout**" tab in the "**View**" drop-down menu. The screenshot destination folder can be set from **"Edit à Preferences à Path**". However, screenshot quality is limited by screen resolution, and furthermore, such a method does not allow important parameters, like background transparency or file format, to be set.

a. To save high quality images of the rendered volume, close the "**SoExaminerViewer**" (having multiple viewers open simultaneously causes the rendering iteration process to freeze) and click on the output port of the "**SoRenderArea**" module in the image acquisition block, to visualize the volume in the "**Output Inspector**" (if closed, the "**Output Inspector**" window can be re-opened by clicking on the layout button located in the bottom right corner of the main software window (see **Section 1.1**)—or via the "**Layout**" tab in the "**View**" menu—and selecting "**Developer**". This will be the reference window for image acquisition, and it is independent from the "**SoExaminerViewer**". If needed, increase the size of the "**Output Inspector**" window (it can also be detached from the side panel) and click on the "**View all**" button to center the volume. Move the volume to the desired position and click the "**Set home**" button. As mentioned before, increasing both the iteration and light bounce number in the "**SoPathTracer**" (see step 2b**)** may improve rendering quality. Such changes, together with the output image size, however, impact image acquisition time and must be chosen depending on several factors such as image purpose, time availability, and hardware configuration. In general, 2500-5000 iterations guarantee an optimal rendering quality and, though it must be evaluated for each sample, 2-5 light bounces can also deliver extra quality (however, as stated in step 2, most of the renderings displayed in the article have been obtained with "**Num**

**Bounces**” set to 1). The progression speed of the “**Current Iteration**” counter in the “**Main**” panel (see step 2b**)** can be used to estimate the impact of light bounces and iteration number on image acquisition time and to set appropriate values to maximize quality/time ratio. However, such estimation must take into account the difference between the “**Output Inspector**” window size and the final image output size, which could be set to be several times bigger (see b, below). In certain cases, the use of the “**Seek**” function (see **Section 1.2**) in conjunction with “**Aperture**” value modification (see step 2b) can contribute to scene realism by generating images with depth of field effect. This opportunity, however, depends on sample type and rendering purpose, and it has to be evaluated case by case.

b. Open the “**OffscreenRenderer**” module panel to choose the desired parameters for the output image. A transparent background is set by selecting “**Rgba**” in the “**Type**” field, while the output image size (in pixels) can be specified in the “**Size**” field (**4K**: 3840x2160; **HD**:1920x1080; however, any resolution that fits volume geometry can be chosen). Enable the use of stencil buffer and multisampling by flagging the “**Use stencil**” and “**Enable multisampling**” checkboxes. Default values can be kept for the other settings. Define output image filename and format in the corresponding fields of the “**ImageSave**” module’s “**Main**” tab. In the “**Options**” tab, select “**None**” in the TIFF “**Compression**” field. Once all the parameters in the different modules have been set, simply click on the update button on the “**OffscreenRenderer**” balloon or on the “**Update**” button in the module panel. This will trigger the offscreen volume rendering. Any open viewer will freeze, and the software will not respond until the value set in “**Stop at Iteration**” is reached. Once the rendering process is complete, the “**Status**” field at the bottom of the “**ImageSave**” panel will change from “**No image**” to “**Image not saved**”. Click the “**Save**” button in the module panel.

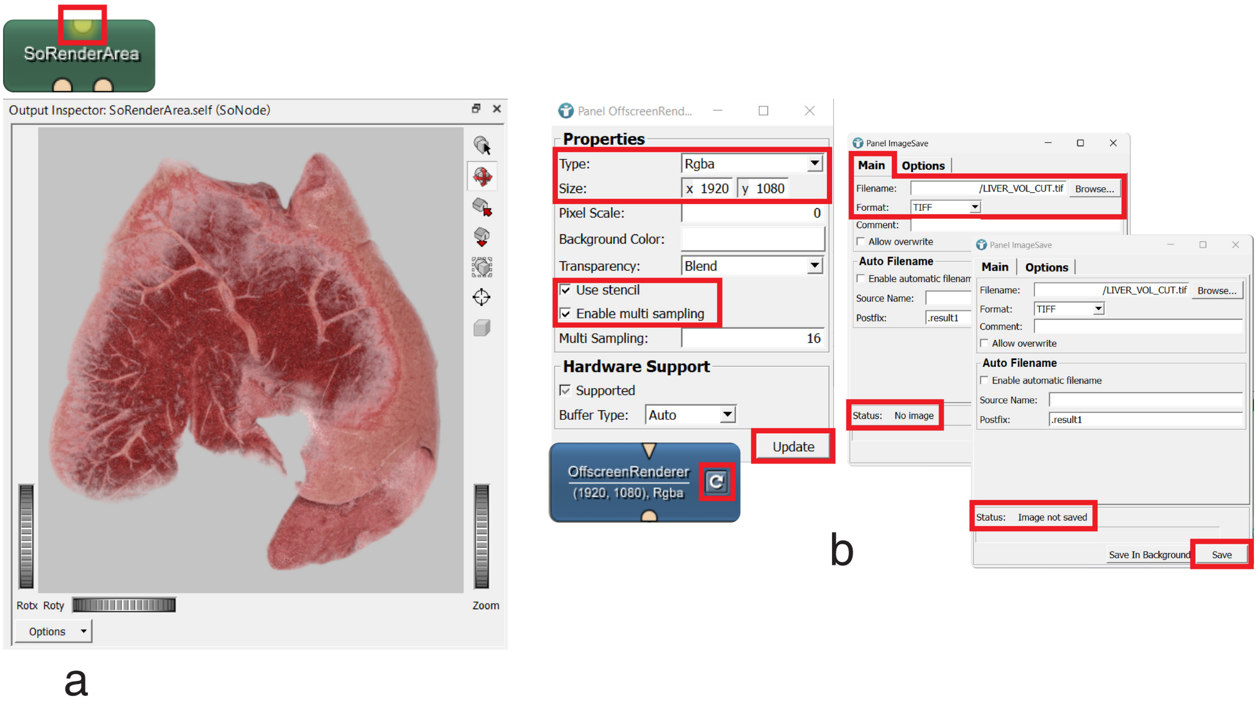

#### Section 3. Rendering of multiple volumes deriving from the same scan

Some additional steps, along with minor modifications to the single volume network, are required to visualize different volumes, all originating from the same scan, in a 3D scene. Each volume of interest, has, indeed, first to be segmented as a separate structure to be, then, represented with distinct appearance properties and features. Though segmentation could be carried out in MeVisLab environment, this step will not be described here, since any software capable of generating binary segmentation masks in a file format compatible with MeVisLab image reading modules (e.g., 3D tiff: .tif, .Nifti: .nii; see figure panel d2 in **Section 2.2**, step 1, and/or check MeVisLab documentation) can be used for such purpose. Segmentation masks will be used in MeVisLab to identify the corresponding subvolumes in the whole CT-scan and, thus, to individually render them. Such an approach allows multiple “secondary” volumes (subvolumes) to be rendered as instances of the “primary” CT-scan volume, significantly reducing computational burden. Real time rendering and image acquisition time are further accelerated by the small internal circuit (MeVisLab offers the possibility to selectively feed in the network only specific information extracted by modules) between the “**itkImageFileReader**”, “**BoundingBox**”, “**SubImage**”, and “**SoPathTracerVolumeInstance**” modules (see inset), aimed to obtain and pass relevant data to automatically crop the subvolumes based on segmentation mask configuration. In some cases, however, the subvolumes can be extracted from the whole volume and each loaded as an independent object. This second strategy may offer more flexibility and will be described in **Section 4**.

##### 3.1 The SmARTR_Multi-Mask_Volume network overview (*)

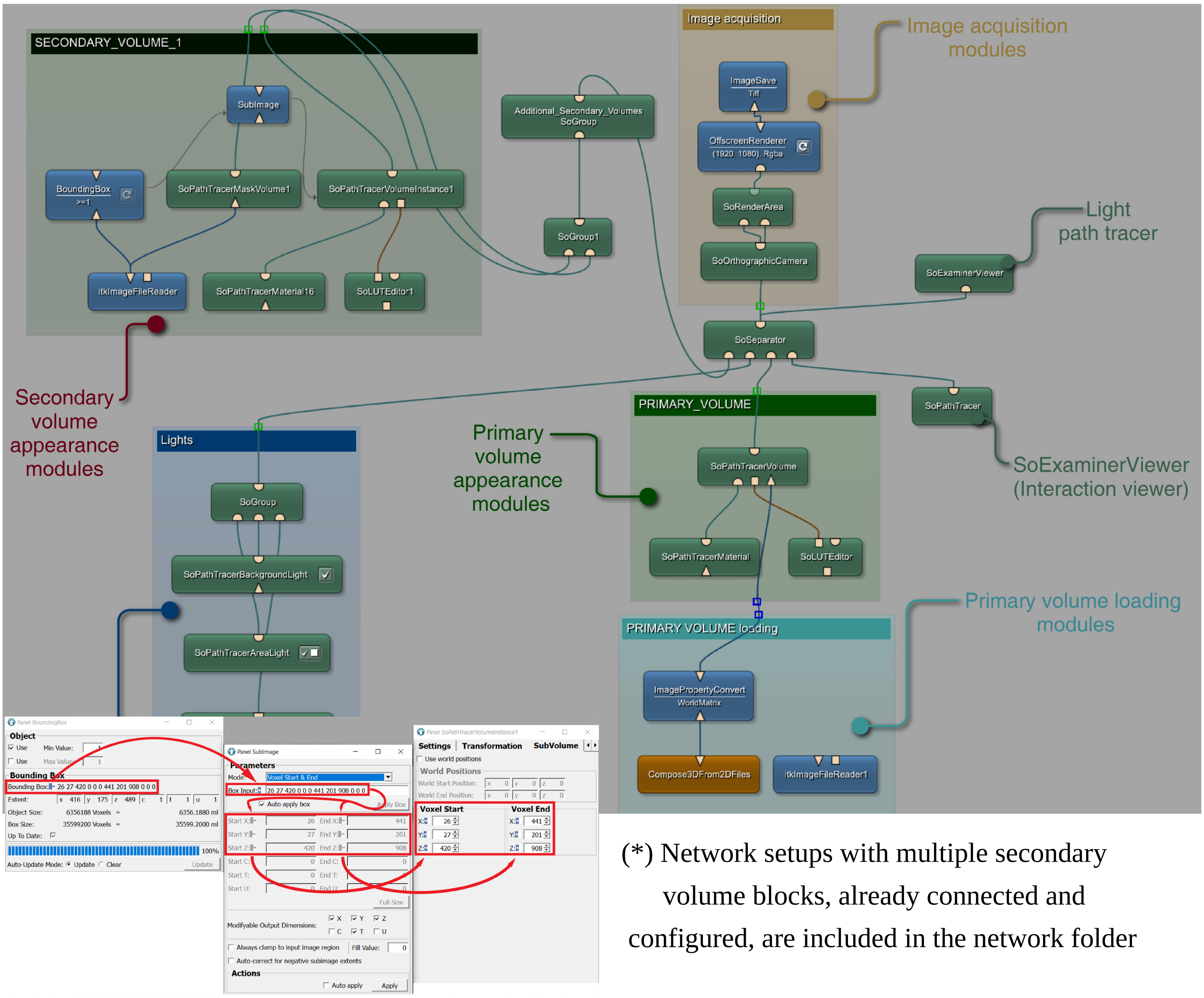


(*) Network setups with multiple secondary volume blocks, already connected and configured, are included in the network folder

##### 3.2 Step-by-step protocol for multiple-volume rendering and acquisition

1. **Load the primary volume (PRIMARY VOLUME loading block)**

Depending on the file type (2D image sequence or single 3D compatible file) follow either steps 1a and b, or step 1d3 in **Section 2.2** to load the sample (in this example, a 2D image sequence file is used).

2. **Configure and visualize the primary volume (PRIMARY_VOLUME block)**

a. Open the "**SoPathTracerVolume**" panel and set the primary "**Volume Name**" (WHOLE, in the example) and the "**Mask Volume Name**" (whole, in the example; see step 2f in **Section 2.2**). The primary "**Volume Name**" will be referenced by the secondary volume modules.

b. Open the "**SoExaminerviewer**" and adjust primary volume LUT (**Section 2.2**, step 2c) just enough to make it visible in the scene. Detailed configuration of the primary volume's appearance is not necessary. In the example, the primary volume corresponds to multiple lizard organs but only a few of them will be rendered as secondary volumes, thanks to the relative segmentation masks.

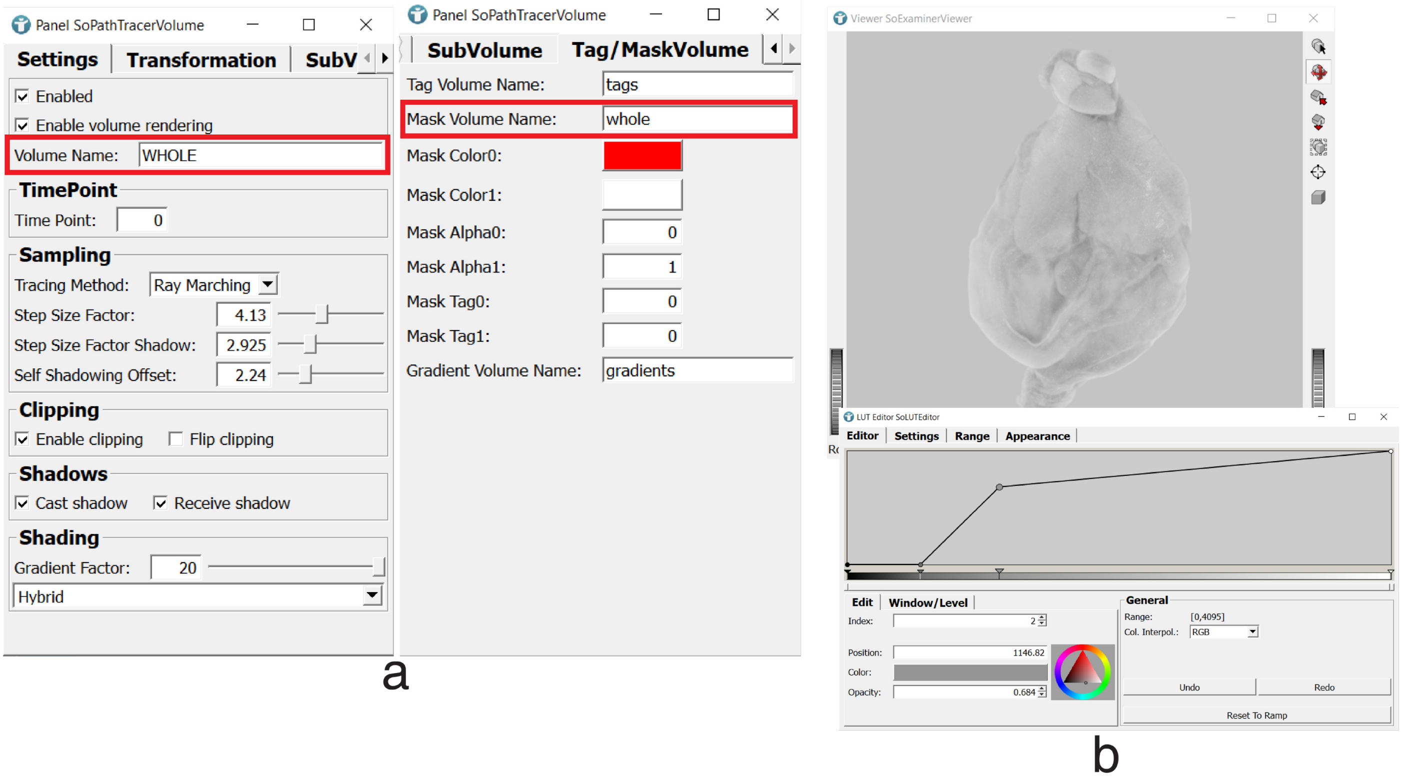


3. **Load the secondary volume segmentation masks and visualize the corresponding subvolume (SECONDARY_VOLUME_1 block)**

Load the segmentation mask compatible file using the "**itkImageFileReader**" belonging to the "**SECONDARY_VOLUME_1**" block (see step1d3 in **Section 2.2**).

- **(*) NOTE:** Old versions of the 3D data analysis and visualization software Amira and Avizo (ThermoFisher Scientific, Thermo Fisher Scientific Inc., USA), by default generate segmentation masks (labelfields) having the "Exterior" material color set to gray rather than black, like in the newer versions. When saved as 3D tiff, such masks are not properly interpreted as binary masks by the "**itkImageFileReader**" module. If working with these platforms for segmentation steps, simply change the "**Exterior**" material color to black in the "**Segmentation Editor**" before saving/exporting the labelfield as a 3D tiff. Alternatively, labelfields can be saved/exported in other formats like Nifti (.nii) or Analyze 7.5 (.hdr, .img) which, though resulting in larger file size, are properly interpreted by MeVisLab reader module.

a. Set the "**Source Volume Name**" and the "**Volume Name**" of the secondary volume in the "**SoPathTracerVolumeInstance**" panel. The "**Source Volume Name**" must match the name of the primary volume ("WHOLE", in the example, see step 2a, above). Switch, then, to the "**Tag/Mask Volume**" tab and specify a name in the "**Mask Volume Name**" field ("liver", in the example). Use the same name (case sensitive) in the "**Volume Name**" field of the "**SopathTracerMaskVolume**" belonging to the same block ("**SECONDARY_VOLUME_1**").

b. Unflag the "**Enable Volume Rendering**" checkbox in the primary volume "**SoPathTracerVolume**" panel to hide the primary volume from the scene, then modify the secondary volume aspect by adjusting the parameters in the relevant modules (see steps 2a-f in **Section 2.2;** the "**SoPathTracerVolumeInstance**" is only a subtype of the **"SoPathTracerVolume"** and displays the same type of adjustable parameters in its main panel).

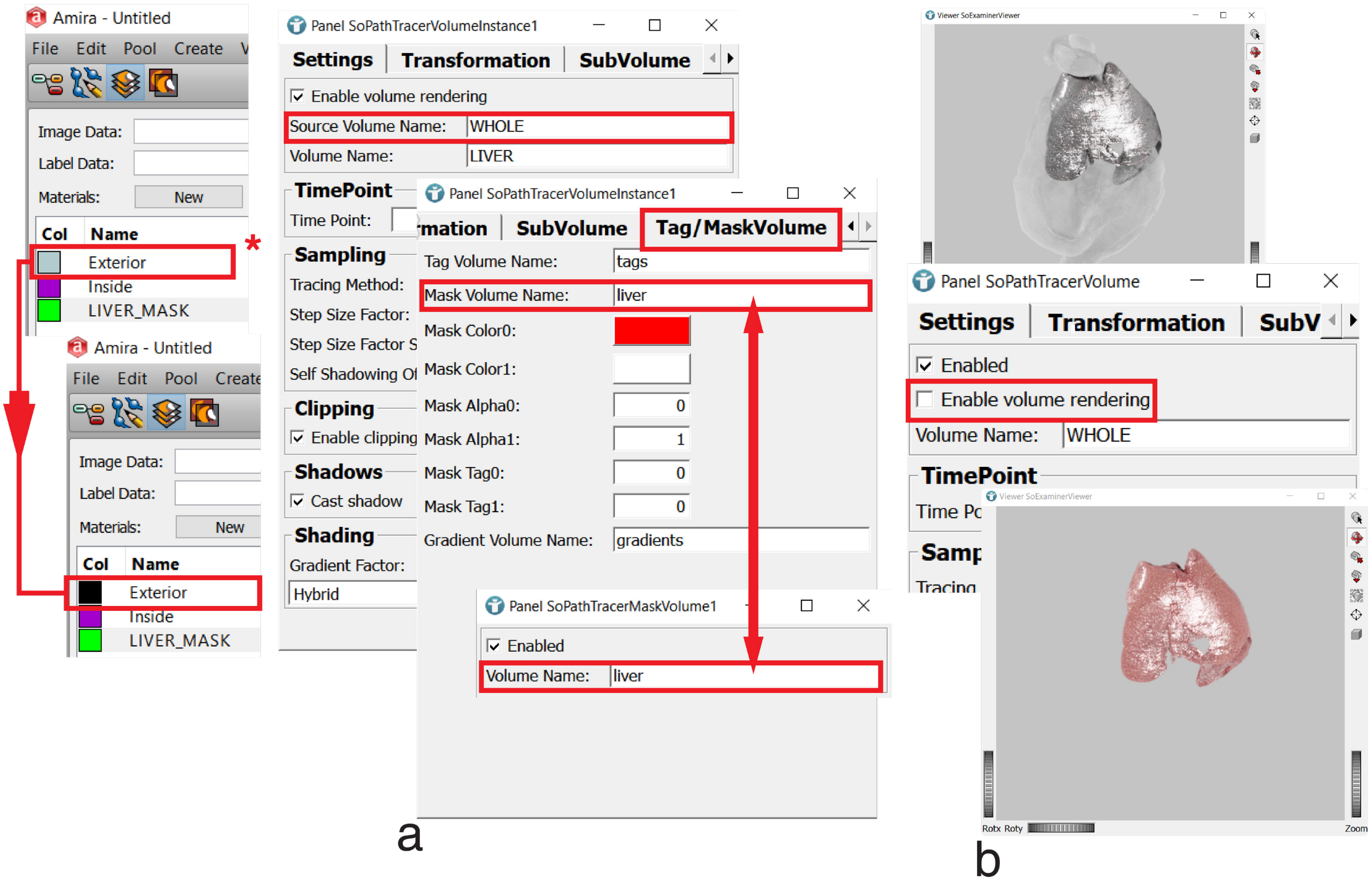

4. **Generate and visualize additional subvolumes of interest**

a. To add an extra secondary subvolume of interest, simply copy-paste the modules of the "**SECONDARY_VOLUME_1**" block together with the "**SoGroup**" module they are connected to. Group the newly generated modules (as SECONDARY_VOLUME_2 - RIGHT_ATRIUM, in the example), wire up the duplicated "**SoGroup**"—which relays the new group outputs—to the "**Additional_Secondary_Volumes**" "**SoGroup**" module, and repeat the steps described in the previous section by consistently specifying names in the relevant module fields, according to the properties of new subvolumes.

b. Use the same procedure to extend the network to all the subvolumes of interest and visualize them.

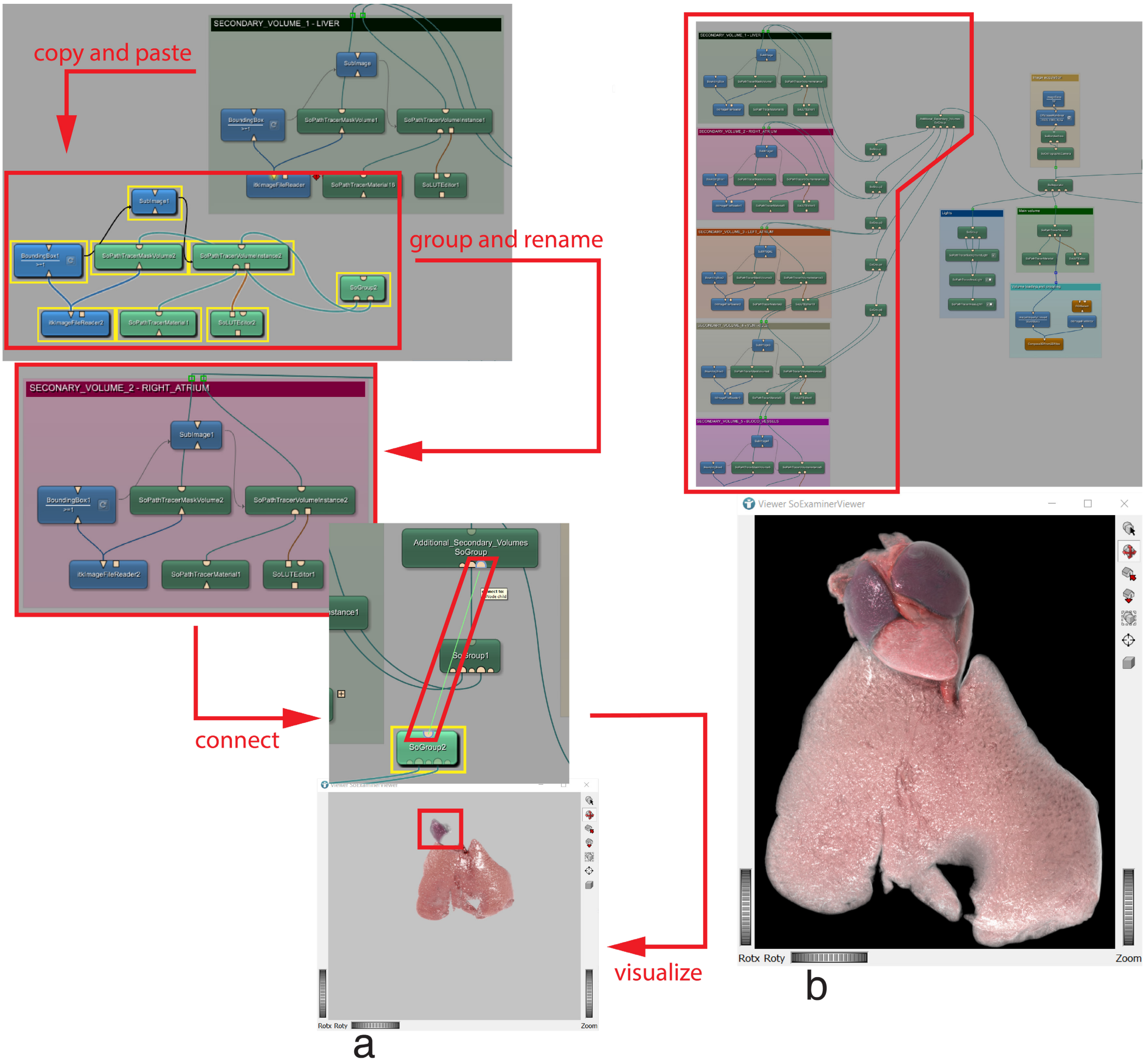


c. **(OPTIONAL)** Cutaways can be generated in the secondary volumes by (c1) connecting an "**UndoManager**" to a "**SoVolumeCutting**", and the latter to the second input (the right port) of a

"**Mask**" module. (c2) Connect the three loaded modules to the network as shown in figure c2 and configure the "**SoVolumeCutting**" module (see **Section 2.2**, step 3a). The "**Mask**" module combines two inputs (two binary masks) from different sources and integrates them—according to some presets ("masked original", in this case)—in a single output. (c3) Since the binary mask associated with the "**SoVolumeCutting**" is only generated upon volume cutting, to create it and achieve a coherent mask integration, it is necessary to draw a contour anywhere in the "**SoExaminerViewer**" window. This will restore the volume appearance, which had been modified upon "**Mask**" module connection to the "**SoPathTracerMaskVolume"**. After this action, cutaways can be performed on the subvolume.

d. (d1) The steps in "c" can be repeated for other groups in the network to display cutaway views of different subvolumes. Multiple "**SoVolumeCutting**" modules can, indeed, be configured (see steps 3a,b in **Section 2.3**) and connected as described in step c2, above. (d2) To facilitate cutting operations on several subvolumes, a "**SoSwitch**" module can be loaded and connected to the "**SoExaminerViewer**". Multiple "**SoVolumeCutting"** outputs (the rounded top port) can be, then, fed to different "**SoSwitch**" input ports in order to select—by pressing the arrow buttons on the module balloon—which one of the various subvolumes will be affected, at any time, by the cutting**.** (d3) The integrated mask can be saved by connecting an "**itkImageFileWriter**" to the "**Mask**" module output to restore, at any time, the volume cutaway when loaded through and "**itkImageFileReader**" directly connected to the "**SoPathTracerMaskVolume**".

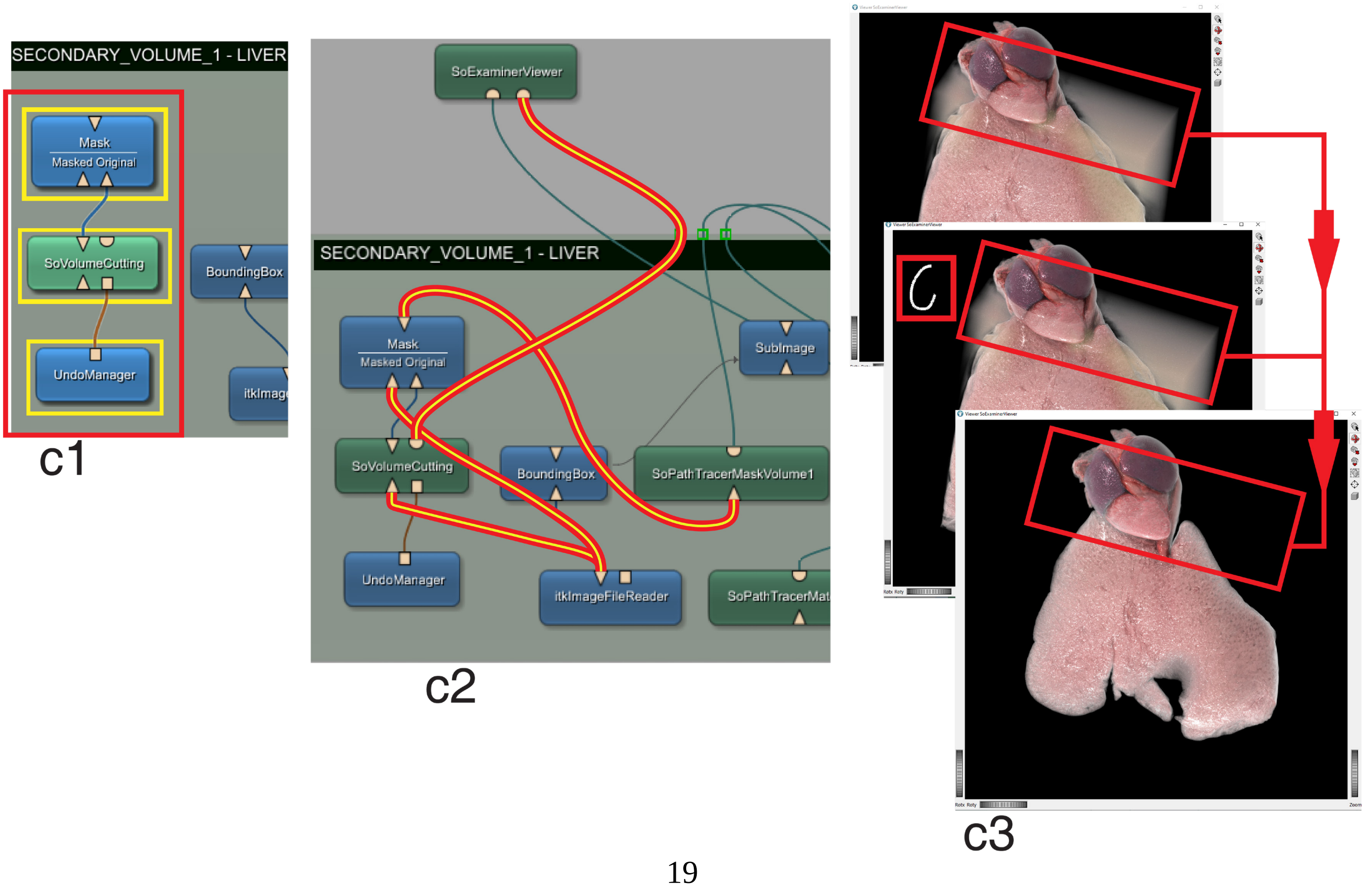

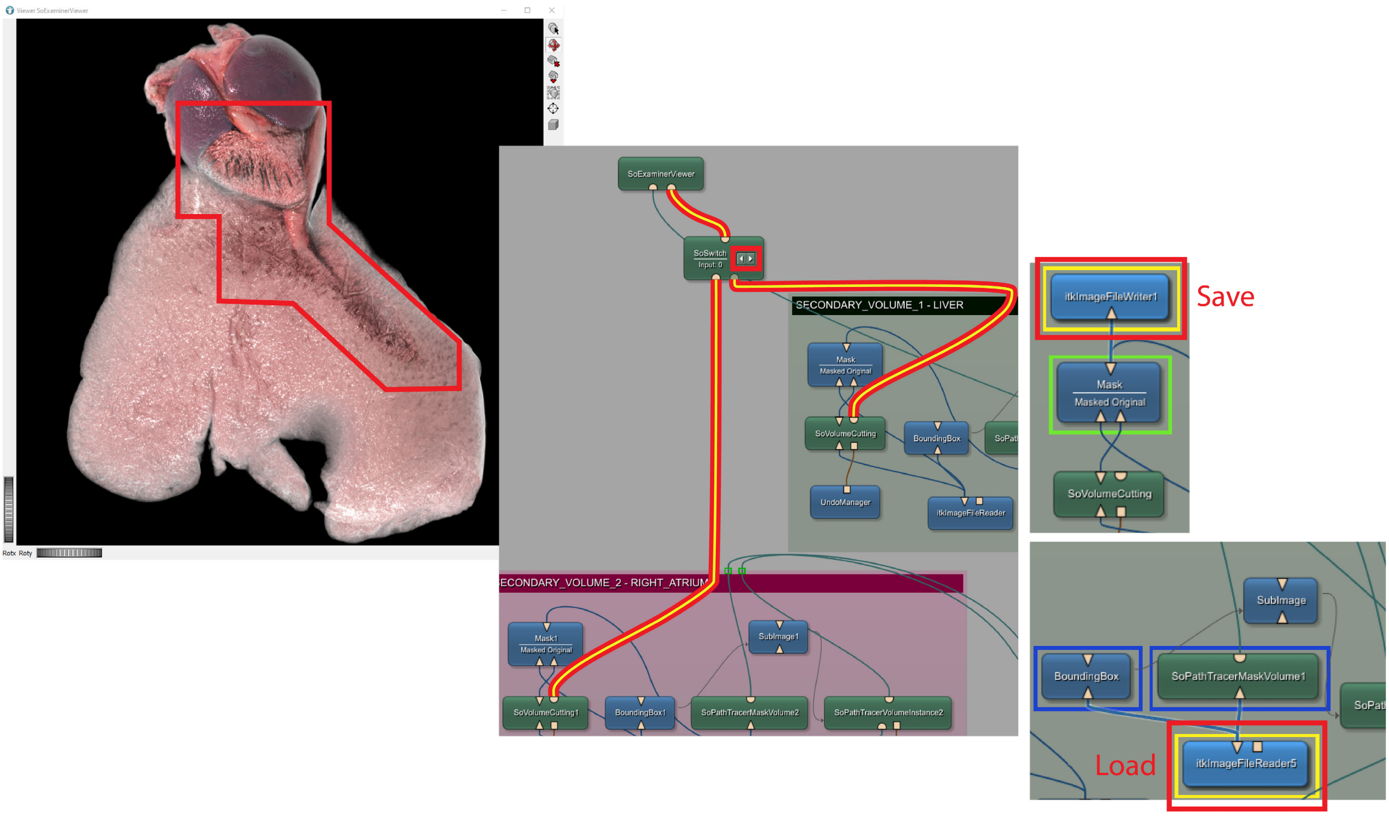


**5. Image acquisition (Image acquisition block)**

Follow step 4a,b in **Section 2.2.**

#### Section 4. Extraction and rendering of multiple independent subvolumes deriving from the same scan

Multi-volume visualization can be achieved also by rendering different subvolumes originating from the same scan as independent entities, unrelated to the full, original volume they derive from. Each subvolume of interest can, in fact, be extracted from the full scan volume and then loaded in the rendering network. Volume extraction is an arithmetic operation allowing regions of a binary segmentation mask to be cut out from the volume they derive from. Such operation can be carried out in the provided **SmARTR_Volume _Extraction** network by simply loading the full scan file and a segmentation mask in the network. The automatically extracted volume can then be saved in a 3D file format (e.g., 3D tiff). However, any software capable of generating volumes in a file format compatible with MeVisLab image reading modules (e.g., 3D tiff: .tif; Nifti: .nii; see figure panel d2 in **Section 2.2**, step 1, and/or check MeVisLab documentation) can be used for volume extraction. The extracted subvolume(s) can be loaded in the visualization network, the **SmARTR_Multi-Independent_Volume** network, which is a slightly modified and extended version of the **SmARTR_Single_Volume** network described in **Section 2**. The network also features a small internal circuit similar to the one characterizing the **SmARTR_Multi-Mask_Volume** network (see **Section 3.1**), aimed to automatically crop the loaded volumes to size and reducing real time rendering and image acquisition time.

##### 4.1 The SmARTR_Multi-Independent_Volume network overview (*)

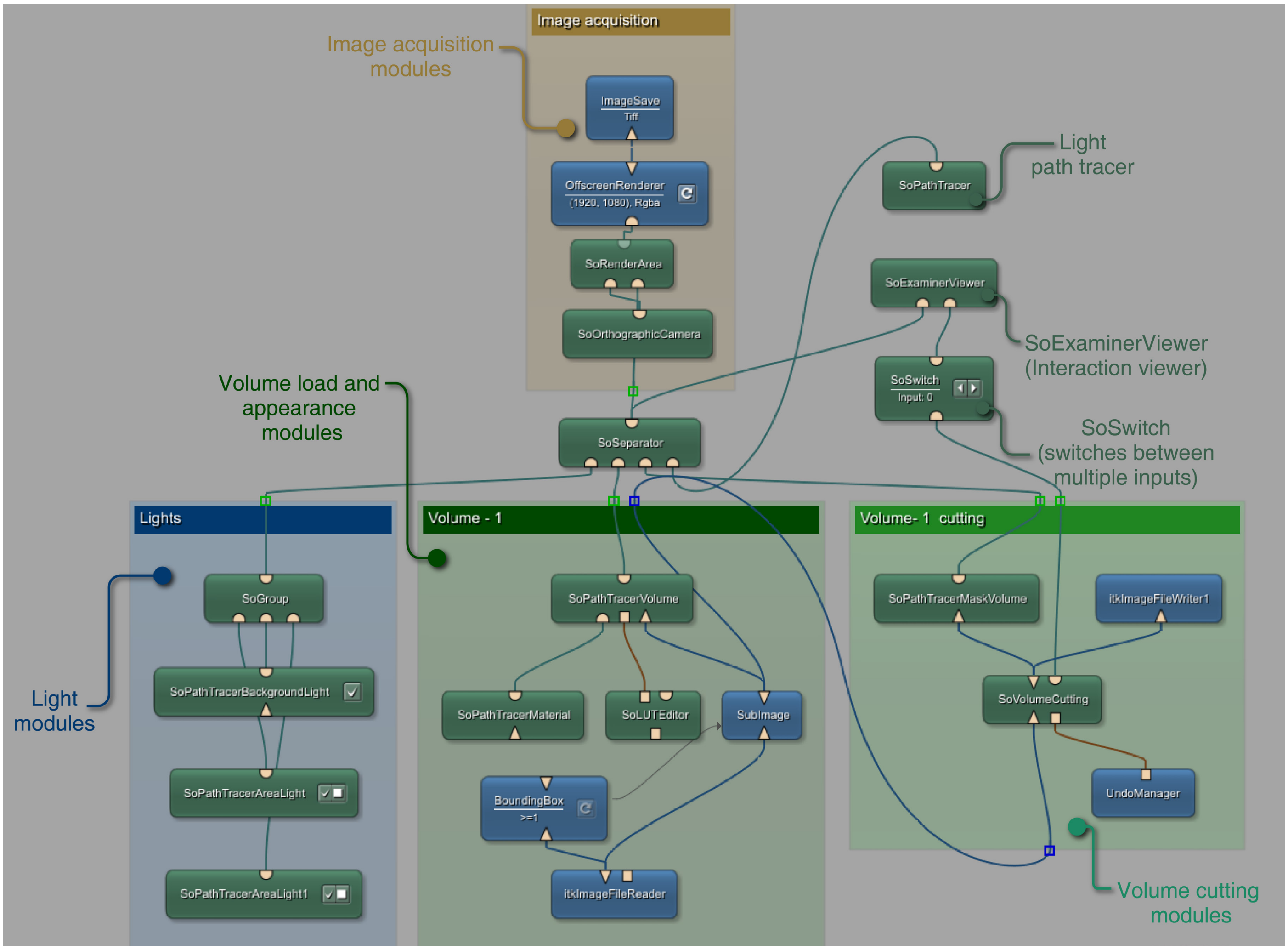


(*) Network setups with multiple independent volume and cutting blocks, already connected and configured, are included in the network folder

#### 4.2 Step-by-step protocol for multi-independent volume rendering and acquisition

##### 1. Use the SmARTR_Volume_Extraction network and obtain the volume(s) of interest

(this step can be skipped if volumes are extracted in external software)

a. Open the provided **SmARTR_Volume_Extraction** network and load the scan data. Depending on the scan file type (2D image sequence or 3d file) use either the “**Compose3DFrom2DFiles**” or “**itkImageFileReader**” (see steps 1a,b or 1d3 in **Section 2.2**). If needed, connect the output of the loading module(s) to the left input port of the “**Arithmetic**” module. Then, load the segmentation mask of the subvolume of interest in the dedicated reader module.

b. The subvolume is automatically extracted. The inputs and output of the “**Arithmetic**” module can be visualized and compared in the “**Output Inspector**” window by clicking on the relative module ports to check the results of volume loading and extraction.

c. Save the extracted subvolume in a suitable format using the “**itkImageFileWriter**” (see **Section 2.2**, step 1d2). If needed, load the masks associated to other subvolumes and save the resulting output files.

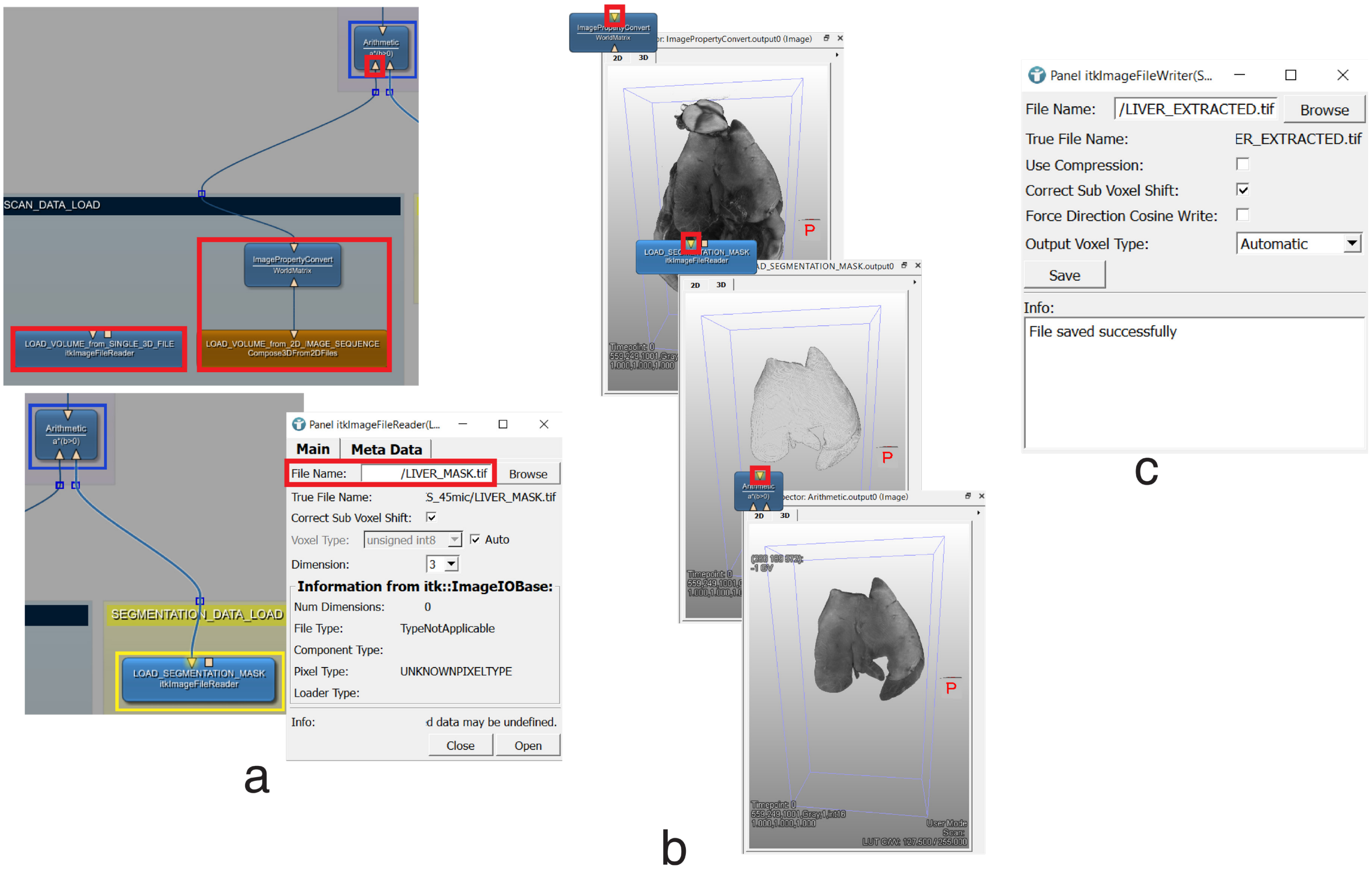

2. **Load and visualize the volume(s) in the SmARTR_Multi-Independent_Volume network (Volume – 1 block)**
   a. Use the "**itkImageFileReader**" in the "**Volume – 1**" block to load the (first) volume in the scene and then, open the "**SoExaminerViewer**". Follow steps 2a-f in **Section 2.2** to modify the appearance of the volume.
   b. To load additional volumes, simply copy-paste the modules belonging to the "**Volume – 1**" block and group them. Alternatively, the "**Volume – 1 cutting**" block modules can be copy-pasted together with the "**Volume – 1**" block if cutaways on the additional volume are planned to be performed (see next subsection). Load the relevant volumes and connect the relative "**SoPathTracerVolume**" outputs to a "**SoSeparator**" module free port. Then, modify each volume's appearance following steps 2a-f in **Section 2.2**.

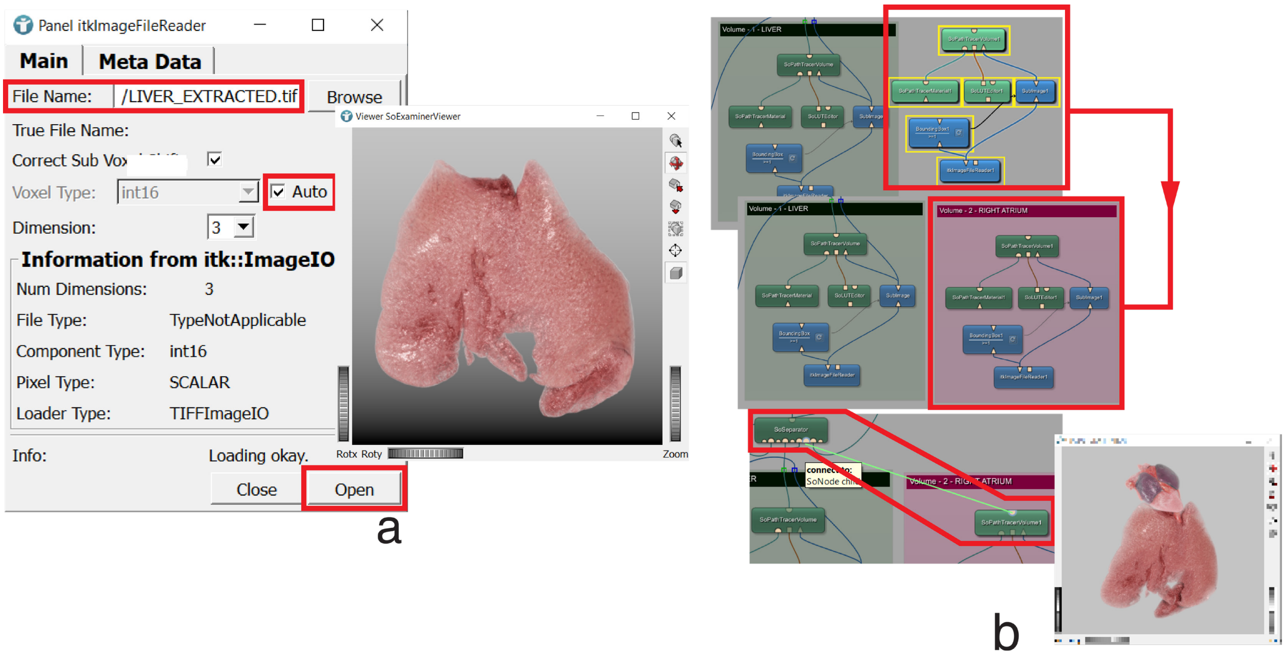


3. **Perform cutaways in one or multiple subvolumes (Volume cutting blocks)**

Cuts in "**Volume – 1**" can be performed thanks to the connected "**Volume – 1 cutting**" block modules. See steps a-c in **Section 2.3** for volume cutting block module configuration. If needed, cutaways can also be performed on several/all volumes present in the 3D scene.

   a. Duplicate the modules of the "**Volume – 1 cutting**" block and configure them according to steps a-c in **Section 2.3**. Then, group and connect them to the relevant modules in the network following the same wiring pattern of the "**Volume – 1 cutting**" modules.
   b. To facilitate cutting operations on multiple volumes, the rounded output port of the additional "**SoVolumeCutting**" modules is connected to one of the "**SoSwitch**" input ports (extra input ports will appear when a wire is drawn towards the module). The "**SoSwitch**" buttons can be used to select which one among the different volumes will be affected by the cutting.

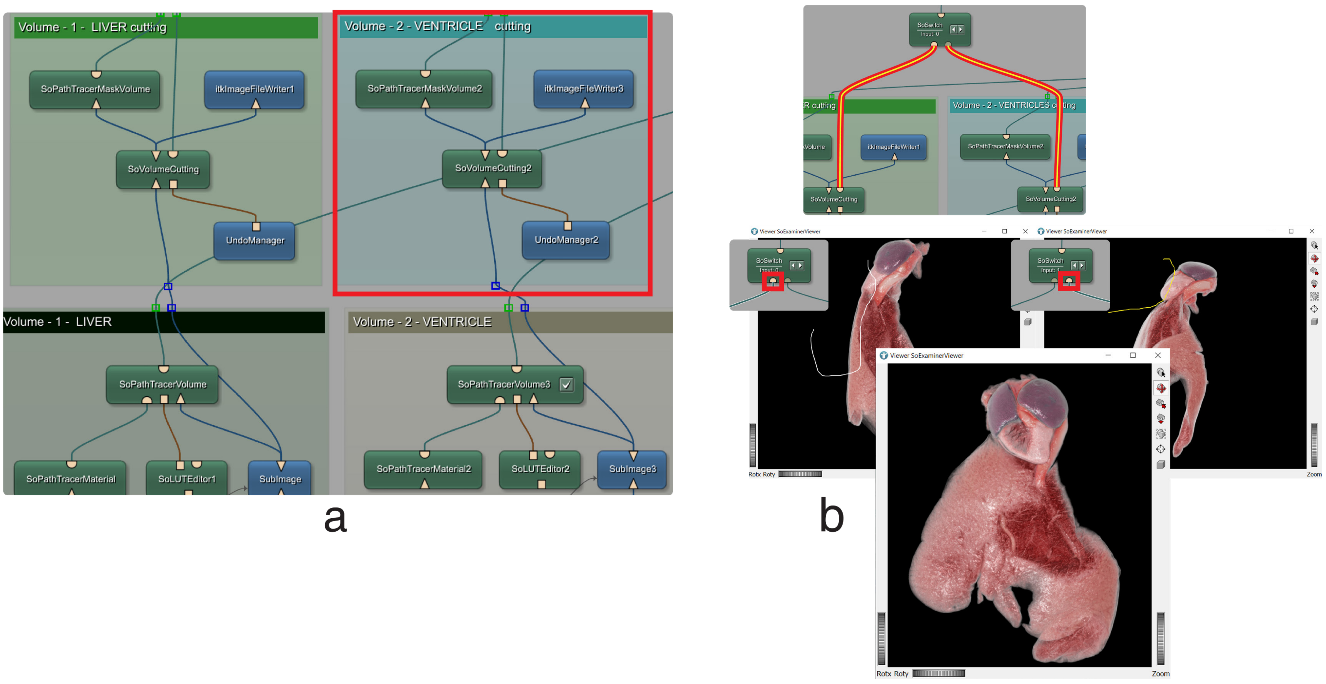


**3*. Perform cutaways in one or multiple subvolumes using a single "SoVolumeCutting" module**

Depending on the needs, multiple extracted volumes can also be cut, simultaneously, by a single "**SoVolumeCutting**" module. This is possible since all the volumes share the same native dimensions, having been extracted from the same scan. However, such an approach comes at the expense of an increased rendering time because to make a single cutting mask fit all the volumes in the scene, they cannot be individually cropped to size.

a. Feed the output of each "**itkImageFileReader**" directly to the corresponding "**SoPathTracerVolume**" module, bypassing the "**BoundingBox**" and "**SubImage**" modules, which can be deleted together with all but one "**Volume Cutting**" blocks. Connect the output of one of the "**itkImageFileReader**" (the one of the liver volume, in the example) directly to the input port of the only "**SoVolumeCutting**" module now present in the network.

b. Specify the same "**Mask Volume Name**" in the "**Tag/Mask Volume**" tab of each of the "**SoPathTracerVolume**" modules and in the "**Volume Name**" field of the "**SoPathTracerMaskVolume**" of the "**Volume Cutting**" block.

c. Ensure the "**SoSwitch**" input port is active. If it is not, use the buttons on the module balloon to activate it. Then, perform volume cutting operations, which will now affect all volumes in the scene that share the same mask properties.

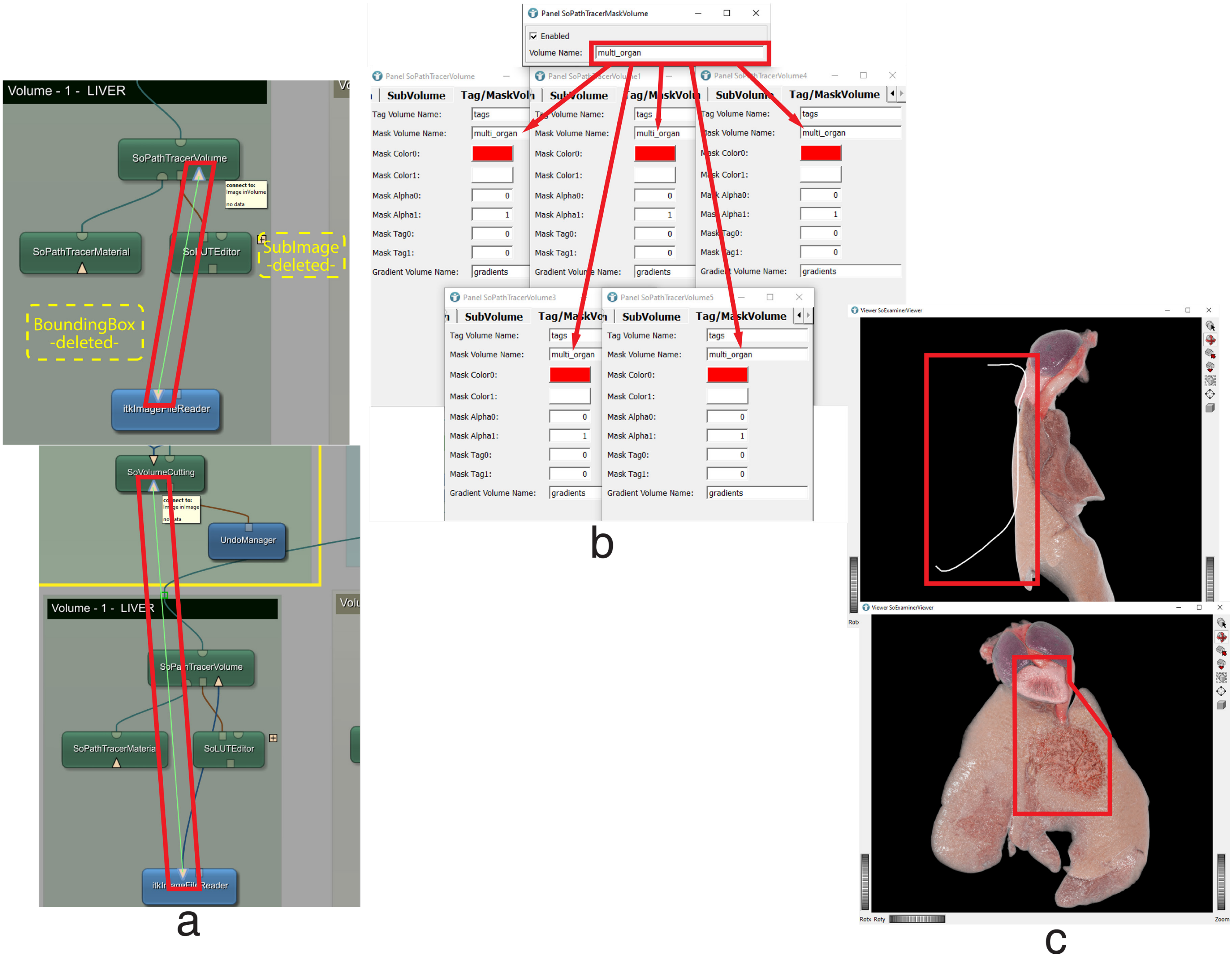


#### 4. Image acquisition (Image acquisition block)

Follow steps 4 a,b in **Section 2.2**

#### Section 5. Generation of cutaway views of multiple nested volumes and non-homogeneous single volumes

In some cases, the volumes of interest can be nested—i.e., located one inside another—like the brain, the skull, and the head soft tissues. In such circumstances, cutaway views represent a suitable way to appreciate the spatial relationships between the different volumes. However, unless all the structures exposed by the outer volume cutting are individually segmented and rendered, the realism of the scene is significantly limited since only one appearance profile can be assigned to each volume. In most cases, in fact, the aspect (color, light interaction features, etc.) of the outermost-volume external surface (e.g., the skin of head) does not coincide with the look of its internal structures (head and neck muscles, see **Fig a**, below). This can also occur when volume cutting is performed on single volumes (such as an organ) that are non-homogeneous, i.e., their internal and external appearance properties cannot be accurately represented by a single appearance profile (see **Fig. b,** below). To get round this issue, the network described in this section, the **SmARTR_Nested_Multi-Volume** network, takes advantage of the output mask generated by the "**SoVolumeCutting**" module connected to the "**Outer volume**" to automatically—and in real time—generate an additional volume (the "**Interposed volume**"), superimposed to the "**Outer volume**" surface exposed following volume cutting. The appearance of this additional volume, rendered by a "**SoPathTracerVolumeInstance**" module can, thus, be configured according to the tissue properties, allowing a realistic representation of the cutaway view (**Figs c,d,** below). Of note, this network can also be selected to provide an additional level of refinement when generating cuts through a single homogeneous volume. Furthermore, when dealing with multiple nested structures, additional cuts can be performed in the "**Intermediate volume**" to better visualize the spatial relationships between the different volumes. Finally, modifying a few values in the automatically generated "**Interposed volume**" appearance properties allows the creation of semitransparent cutaway views **(Figs. e,f,** below). This approach offers an alternative visualization style for displaying multiple nested volumes.

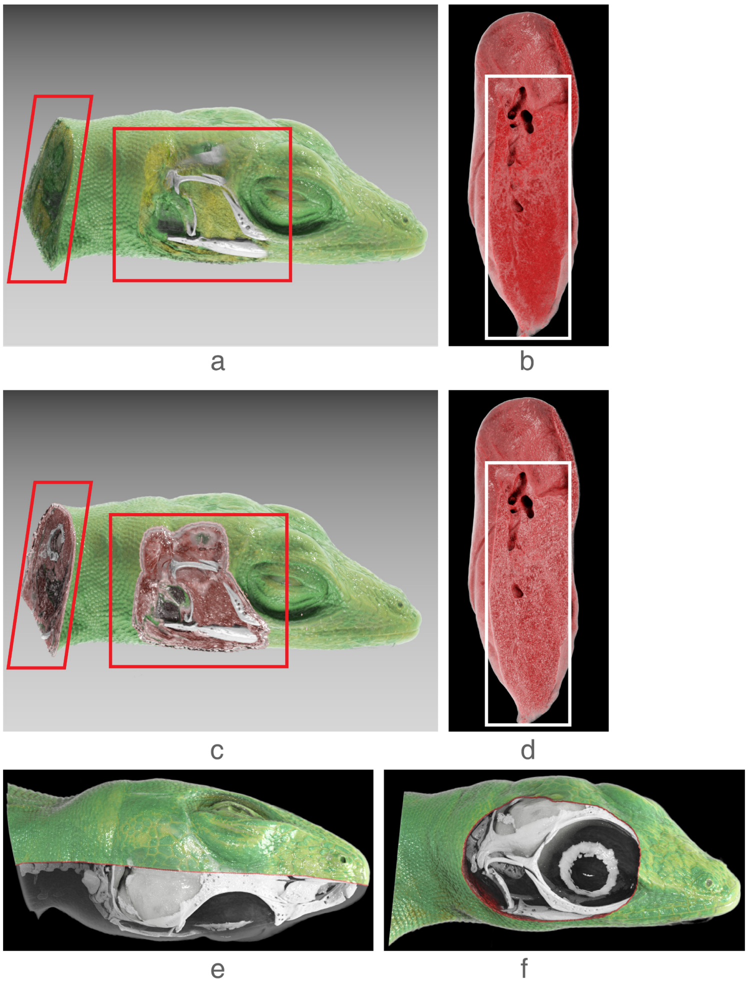

#### 5.1 The SmARTR_Nested_Multi-Volume network overview

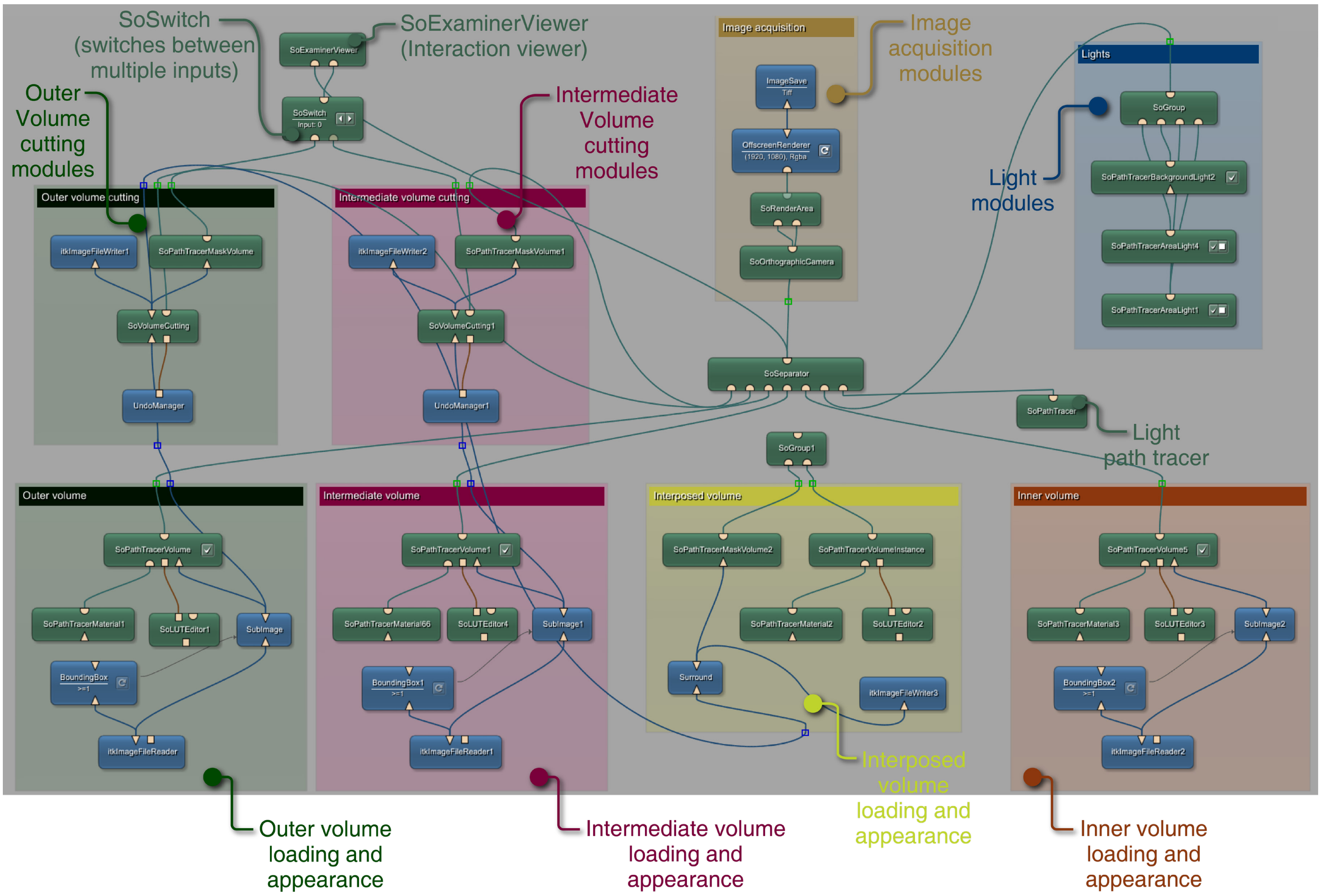

##### 5.2 Step-by-step protocol for cutaway views of multiple nested volumes and non-homogeneous single volumes

1. **Load the nested volumes (Outer, Intermediate, and Inner volume blocks)**
   Use the "**itkImageFileReader**" modules in each block to open the volume files/file (see step 1d3 in **Section 2.2**). If dealing with 2D image sequences, load and connect the relevant modules to open such a file type and create a 3D volume (follow steps 1a and b, in **Section 2.2**). In this practical example, which illustrates only the multiple nested volume scenario, the lizard head is the "**Outer volume**", the skull is the "**Intermediate volume**", and the brain is the "**Inner volume**". The "**Interposed volume**" will be automatically generated and rendered upon cutting the head volume. If dealing with a single, non-homogeneous volume, instead, simply load it using the "**itkImageFileReader**" in the "**Outer volume**" block or, if needed, use other file loading modules (see steps 1a and b, in **Section 2.2**). Then, follow steps 2a,b and 3, below.

2. **Adjust volume appearance, cut the outer and the intermediate volume to reveal the inner volume (Outer, Intermediate, and Inner volume blocks; Outer and Inner cutting blocks)**

a. Change the relevant module parameters (see steps 2a-f in **Section 2.2**) to modify the appearance of the different volumes in the scene.

b. Connect the "**SoGroup**" module from the "**Interposed volume**" block to the network. Similarly to what described in **Section 3.1**, step 3c, upon connection of the "**SoGroup**" to the network, the appearance of the "**Outer volume**" will change. Simply draw a contour anywhere in the "**SoExaminerViewer**" to restore the "**Outer volume**" appearance (be sure that the left input is highlighted in the "**SoSwitch**" module). After this action, the necessary cutting operations can be performed. The "**Interposed volume**" is automatically generated upon cutting and updated thanks to the combined action of the "**Surround**" module, which erodes to its borders the "**SoVolumeCutting**" mask generated upon cutting, and by that of the "**SoPathTracerVolumeInstance**" which exclusively renders the portion of volume specified by the modified cutting mask (see steps 2a and 3a in **Section 3** for more info about "**SoPathTracerVolumeInstance**" module configuration). Modify the appearance of the automatically generated "**Interposed volume**" by adjusting the parameters in the relevant modules to improve the realism of the cutaway view (see steps a-f in **Section 2.2**; the "**SoPathTracerVolumeInstance**" is a subtype of the "**SoPathTracerVolume**" and displays the same type of adjustable parameters in its main panel).

**(*) NOTE:** to realize a semitransparent cutaway view, first switch to a black background. This can be achieved by setting black as background color in the "**SoExaminerViewer**" property panel (see step 2a in **Section 2.2**) and by selecting "**Blend**" as the "**Blend Mode**" in the "**Main**" tab of the

"**SoPathTracer**" panel (see step 2b in **Section 2.2**). Then, open the panel of the "**SoPathTracerVolumeInstance**" in the "**Interposed volume**" block and navigate to the "**Tag/MaskVolume**" tab (if not visible in the panel, click on the black arrowheads in the panel top-right corner; see step 2f, **Section 2.2**). Set the "**Mask Color0**" to black for the highest level of chromatic neutrality of the cut region (if required for specific visualization purposes, other colors can be selected) and assign to "**Mask Alpha0**" a suitable value between 0 (full transparency) and 1 (full opacity). Values in the range of 0.01 and 0.05 generally give the best result for this kind of visualization. The "**Interposed volume**" appearance can be further modified by adjusting the parameters in the relevant modules (see **Section 2.2,** steps 2c-f). Refining light orientation and "**SoPathTracerVolumeInstance**" properties have a strong impact in reaching a suitable transparency effect. Furthermore, the "**SoLUTEditor**" can be configured to highlight in color the border between the semitransparent and the opaque portions of the sample.

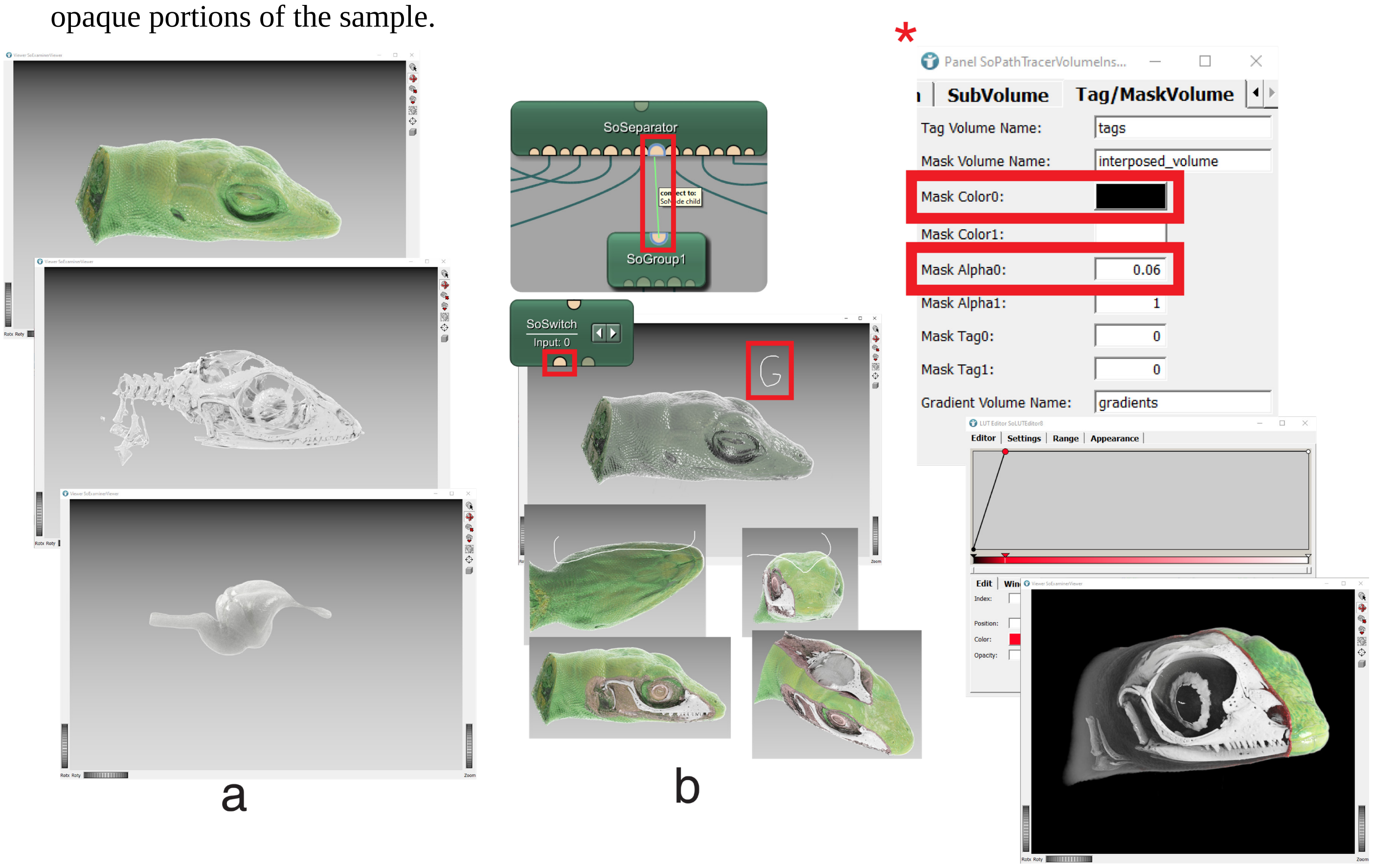


c. **(OPTIONAL)** In rare cases, the configuration of the "**Interposed volume**" may generate inconsistencies resulting in extra material appearing outside the area of interest during cutting operations. This is due to the presence of dense material surrounding the cut volume ("**Outer volume**"). This dense material often derives from procedures designed to prevent sample movements or dehydration during imaging, such as wrapping the sample in tissue paper moistened with contrast agent solvent. This extra material often lies outside the sample and can be deleted after image acquisition in a photo-editing software. However, if needed, it can also be directly eliminated from the 3D scene. To remove it, (c1) load an "**UndoManager**", a "**SoVolumeCutting**" and a "**Mask**" module. Connect the "**UndoManager**" to the "**SoVolumeCutting**", and the latter to the left input (the left port) of the "**Mask**" module. Wire up the modules to the network as shown in the bottom panel of fig. c1, below. Then, (c2) connect the rounded output port of the "**SoVolumeCutting**" to the "**SoSwitch**" module already present in the network. Configure the "**SoVolumeCutting**" properties (see **Section 2.2**, step 3a,) and use the buttons on the "**SoSwitch**" module balloon to select the "**Interposed Volume**" as the target of the cutting operations (see step d2 in **Section 3.4**). Remove the extra material from the scene using the same technique employed for cutting away portions of the volume of interest. This method ensures that the unwanted material is effectively eliminated, maintaining the integrity of the desired sample area.

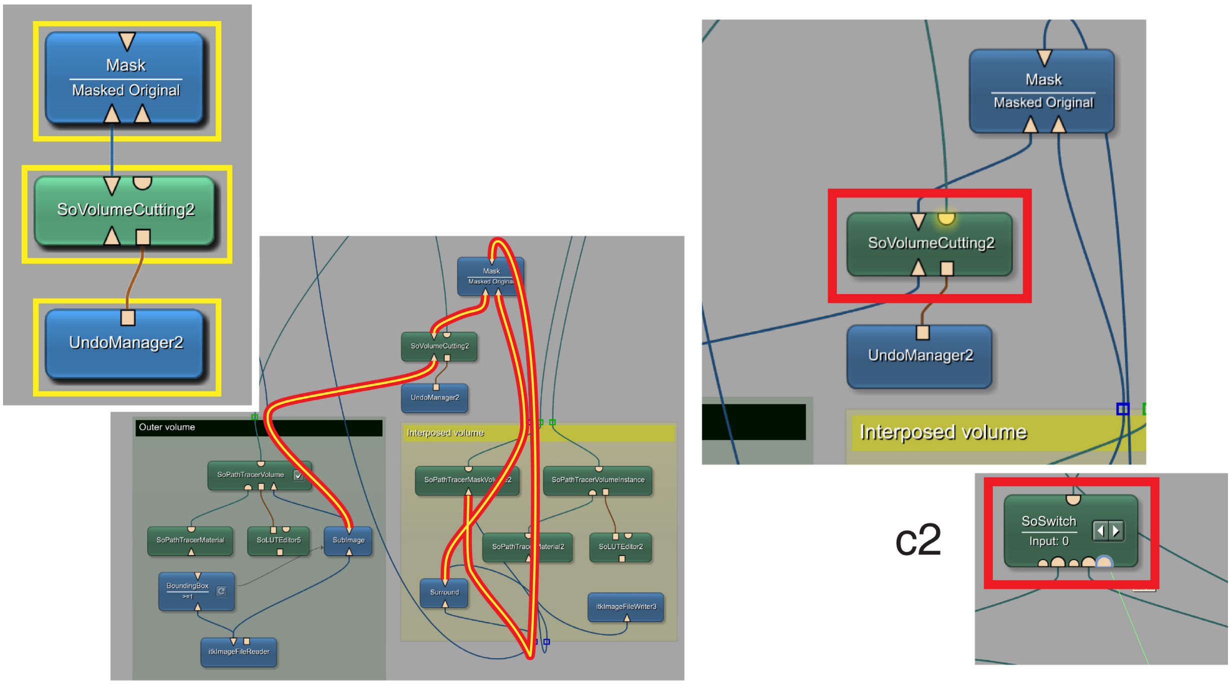


c1 c2

d. Use the "**SoSwitch**" buttons to select the "**Intermediate volume**" and generate cuts at relevant locations to better reveal the "**Inner volume**". Save all the generated cutting masks using the different "**itkImageFileWriter**" located in the various volume blocks (see step d3 in **Section 2.2**).

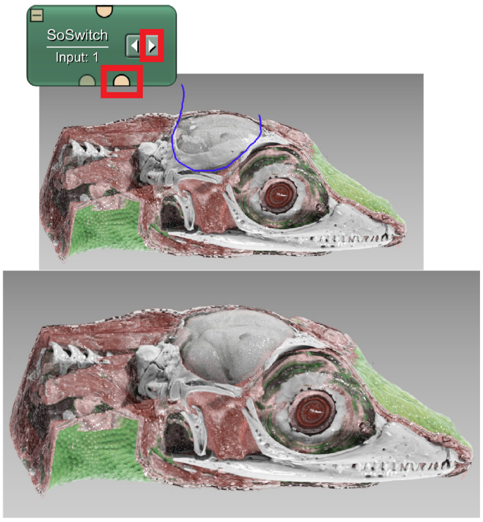


d

#### 3. Image acquisition (Image acquisition block)

Follow steps 4 a,b in **Section 2.2**.

#### Section 6. Skin pattern drawing and rendering

Skin color patterns are a distinctive animal trait, sometimes reaching a bewildering level of complexity. Though it would dramatically enhance rendering lifelikeness, the accurate reproduction of intricate skin color schemes on directly rendered animal volumes derived from internal imaging methodologies is, for technical reasons, unfeasible. The acquisition of detailed information on sample coloration is, in fact, one of the key features of other imaging techniques, like 3D photogrammetry or surface scanning. However, the network described in this section allows simple skin patterns, such as multi-colored stripes and/or patches, to be recreated on volume data, thus increasing rendering realism by approximating animal natural coloration. Modules usually employed to produce cutaway views are here combined and configured (“**PATTERN_DESIGN**” block) to draw and visualize color schemes on the 3D volume in real time. Importantly, each generated pattern can be rendered with its own specific color and appearance profile, independent from those of the other volumes in the scene.

##### 6.1 The SmARTR_Pattern_Drawing network overview

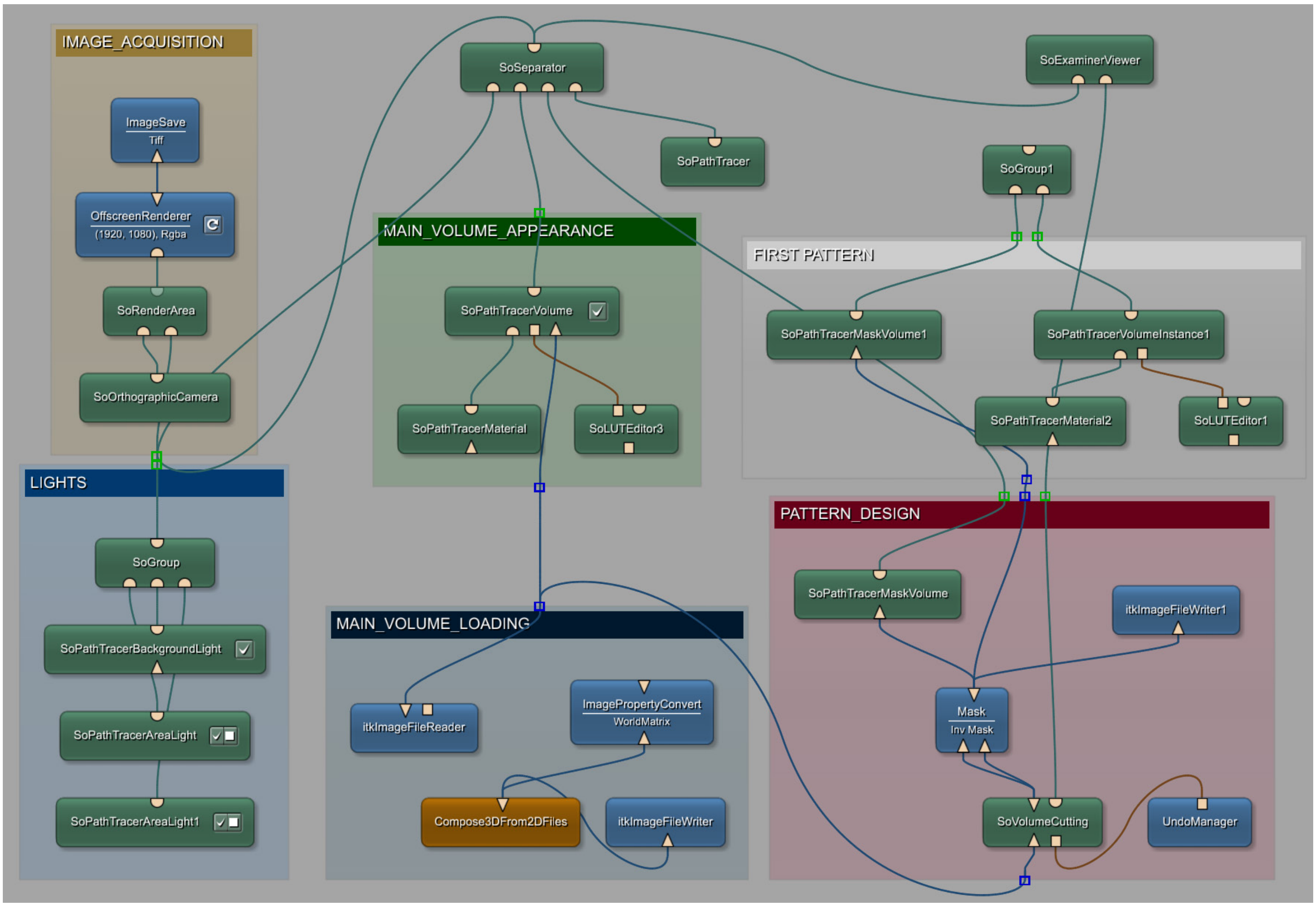

##### 6.2 Step-by-step protocol for color pattern drawing on 3D volumes

In the practical example, the simple green-white (dorso-ventral) pattern of the green anole lizard will be reproduced. The steps needed to create additional color patterns will also be detailed, though in this specific case, the procedure will be used to give the lizard eye a more realistic appearance rather than to generate a second skin color pattern. The focus of the practical example will be on the side chosen for image acquisition, although simple color patterns can be created and refined to be displayed on both sides of the volume (see steps 3*).

1. **Load the volume (MAIN_VOLUME_LOADING block)**

   Use the "**itkImageFileReader**" module in the "**MAIN_VOLUME_LOADING**" block to load the example volume file (see step 1d3 in **Section 2.2**). If dealing with 2D image sequences, connect the relevant modules to open such a file type and create a 3D volume (follow steps 1a and b, in **Section 2.2**).

2. **Adjust main volume appearance and create the first color pattern (MAIN_VOLUME_APPEARANCE and PATTER_DESIGN block)**

a. Use the modules in the "**MAIN_VOLUME_APPEARANCE"** block to set the appearance of the loaded volume (see steps 2a-f in **Section 2.2**), then, connect the "**SoGroup**" receiving the outputs from the "**FIRST_PATTERN**" group to the network. Similarly to what described in **Section 3.1**, step 4c, upon connection of the "**SoGroup**" to the network, the appearance of the main volume will change since the binary mask associated with the "**SoPathTracerMaskVolume**" module in the "**FIRST_PATTERN**" block will be created only upon pattern drawing. Simply draw a contour anywhere in the "**SoExaminerViewer**" to restore the "**MAIN VOLUME**" appearance.

b. The possibility to draw color patterns on the main volume depends on the correct specification of volume and mask names in the modules belonging to the "**MAIN_VOLUME_APPEARANCE"** and "**FIRST_PATTERN**" blocks. The "**Source Volume Name**" field of the "**SoPathTracerVolumeInstance**" module in the "**FIRST_PATTERN**" block must match the "**Volume Name**" of the "**SoPathTracerVolume**" in the "**MAIN_VOLUME_APPEARANCE"** group ("**ANOLIS_HEAD**", in the example). Furthermore, the name specified in the "**Mask Volume Name**" field of this latter module (under the "**Tag/Mask Volume**" tab, see **Section 2.2**, step 2f) and the "**Volume Name**" field of the "**SoPathTracerMaskVolume**" in the "**PATTERN_DESIGN**" block must be identical. In a similar manner, the name indicated in the "**Mask Volume Name"** field of the "**SoPathTracerVolumeInstance**" in the "**FIRST_PATTERN**" group must be identical to the one in "**Volume Name**" field of the "**SoPathTracerMaskVolume**" belonging to the same block.

c. With the "**SoVolumeCutting**" set to "**Interior**" (see steps 3a,b in **Section 2.2**) draw a contour on the region of interest of the main volume (the ventral part of the lizard head, in the example). The area encircled by the contour will acquire a different coloration than the rest of the volume, according to the "**SoLUTEditor**" properties (see step 2c in **Section 2.2**) in the "**FIRST_PATTERN**" group. Setting the "**SoVolumeCutting**" to "**Exterior**" allows the shape of the colored region to be refined as unwanted colored areas can be removed. Once refined, save the generated mask using the "**itkImageFileWriter**" (see step d3 in **Section 3.4**) connected to the "**Mask**" module in the "**PATTERN_DESIGN**" block.

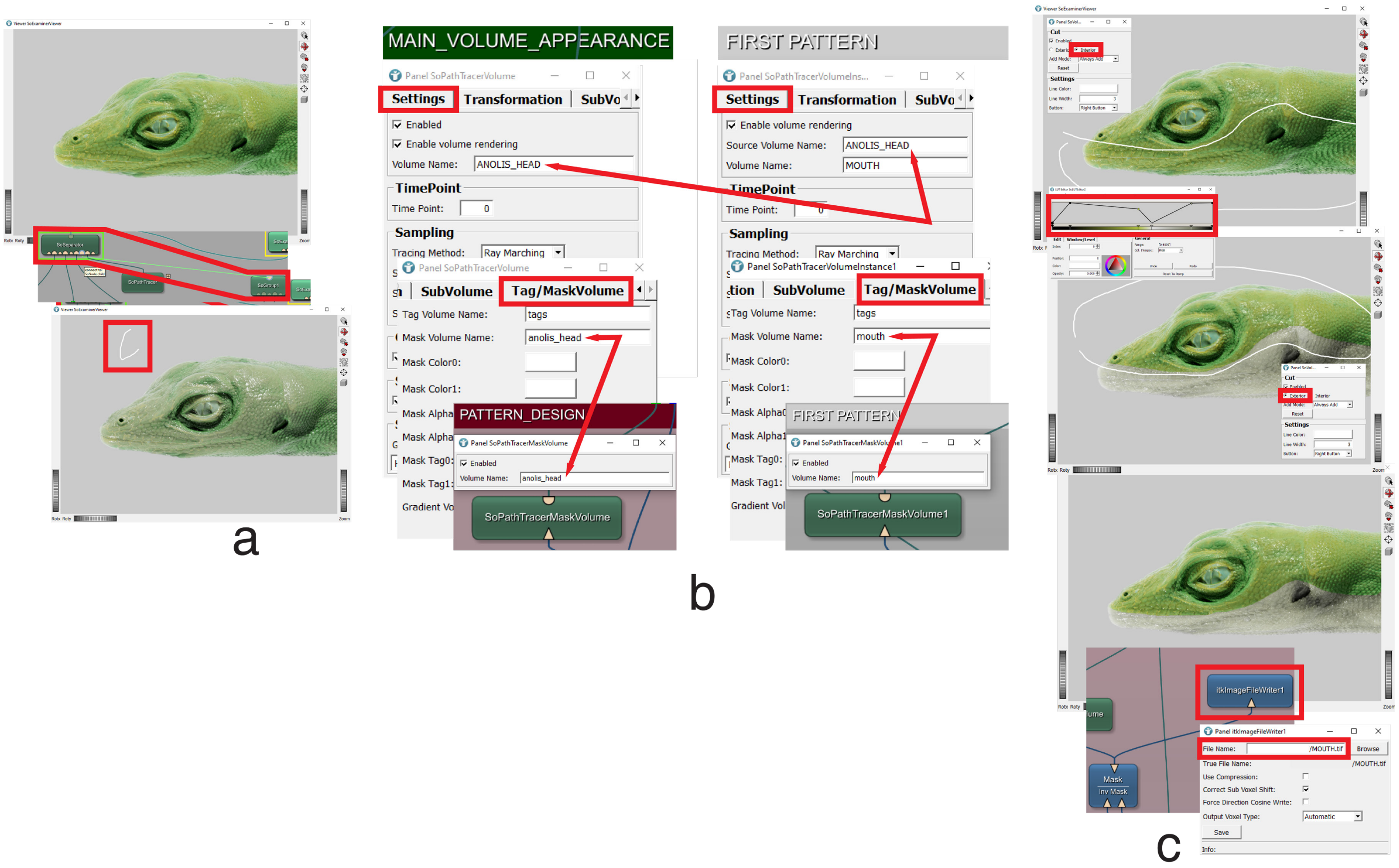


3. **Create the second pattern**

a. Reset the "**SoVolumeCutting**" (see **Section 2.2**, step 3a) before proceeding with the creation of the second color pattern. Then, load an "**itkImageFileReader**" module, connect it directly to the "**SoPathTracerMaskVolume**" in the "**FIRST_PATTERN**" group and open the saved first color pattern mask. Copy-paste the 5 "**FIRST_PATTERN**" modules (including the "**SoGroup**" module) and group them ("**SECOND_PATTERN**", in the example). Connect the output of the "**Mask**" module from the "**PATTERN_DESIGN**" block to the "**SoPathTracerMaskVolume**" of the "**SECOND_PATTERN**" group. Then, connect the "**SoGroup**" of the newly created group to an empty input port of the "**SoSeparator**".

b. In the "**SECOND_PATTERN**" block, specify the same name (eye, in the example) in the "**Mask Volume**" field of the "**SoPathTracerMaskVolume**" and in the "**Mask Volume Name**" field of the "**SoPathTracerVolumeInstance**" (in the "**Tag/Mask Volume**" tab). No additional modifications to other modules are required. Draw the second color pattern (in the example, the procedure will be used to change the eye appearance) and adjust the relative appearance properties (see step 2c,e,f in **Section 2.2**).

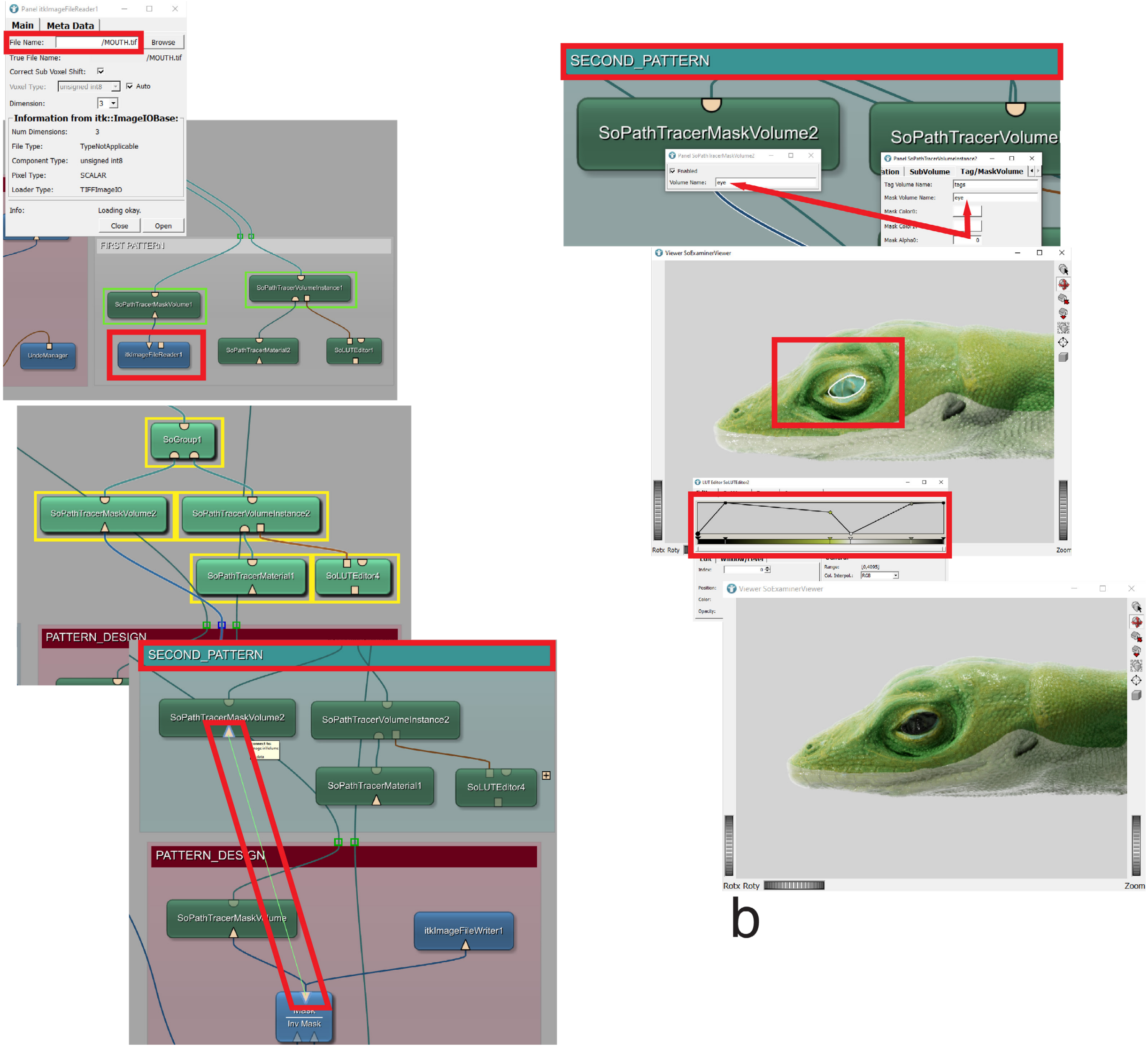


a

b

#### 3*. Creation of bilateral and symmetric skin patterns

Owing to the nature of the methodology involved in the color pattern drawing on the 3D volume, achieving coherent patterns on both sides of a sample requires a multi-step process. To create patterns that share the same appearance on both sides of the volume (e.g., bilateral skin markings, lines, or patches), or patterns that extend symmetrically from one side to the other (e.g., the ventral coloration of a lizard), a pattern is first drawn and refined on one side of the volume.

a. Draw and refine the pattern on one side of the sample as described in step 2c, above. Once done, set the "**SoVolumeCutting**" to "**Exterior**" (see steps 3a in **Section 2.2**) and draw a contour along the sample sagittal mid-line or any other contour that effectively removes only the unwanted markings appeared on the opposite side.

Once ready, the pattern can be saved (see step 2c, above) and loaded (see step 3a, above). New pattern modules can be generated, grouped, renamed (FIRST_PATTERN_OTHER_SIDE, in the example), configured, and connected to the network (see step 3a,b above) to draw the pattern on the other side of the sample.

b. Before drawing the pattern on the opposite side, copy-paste the “**SoLUTEditor**” module from the newly generated pattern block and use a different color for the new pattern to differentiate it from the one on the opposite side. Draw the pattern to achieve the desired symmetry. If any unwanted markings appear on the pattern already drawn on the other side (identifiable by their different color), use the “**SoVolumeCutting**” set to “**Exterior**” and draw a suitable contour to remove them. Once done, save (see step 2c, above) and load (see step 3a, above) the newly created pattern and restore its color by connecting the original “**SoLUTEditor**” (the one which had been copy-pasted before) to the “**SoPathTracerVolumeInstance**”. The 3D volume will now feature seamlessly and symmetrically integrated patterns on both sides. Similar approaches can be used when dealing with bilateral patterns.

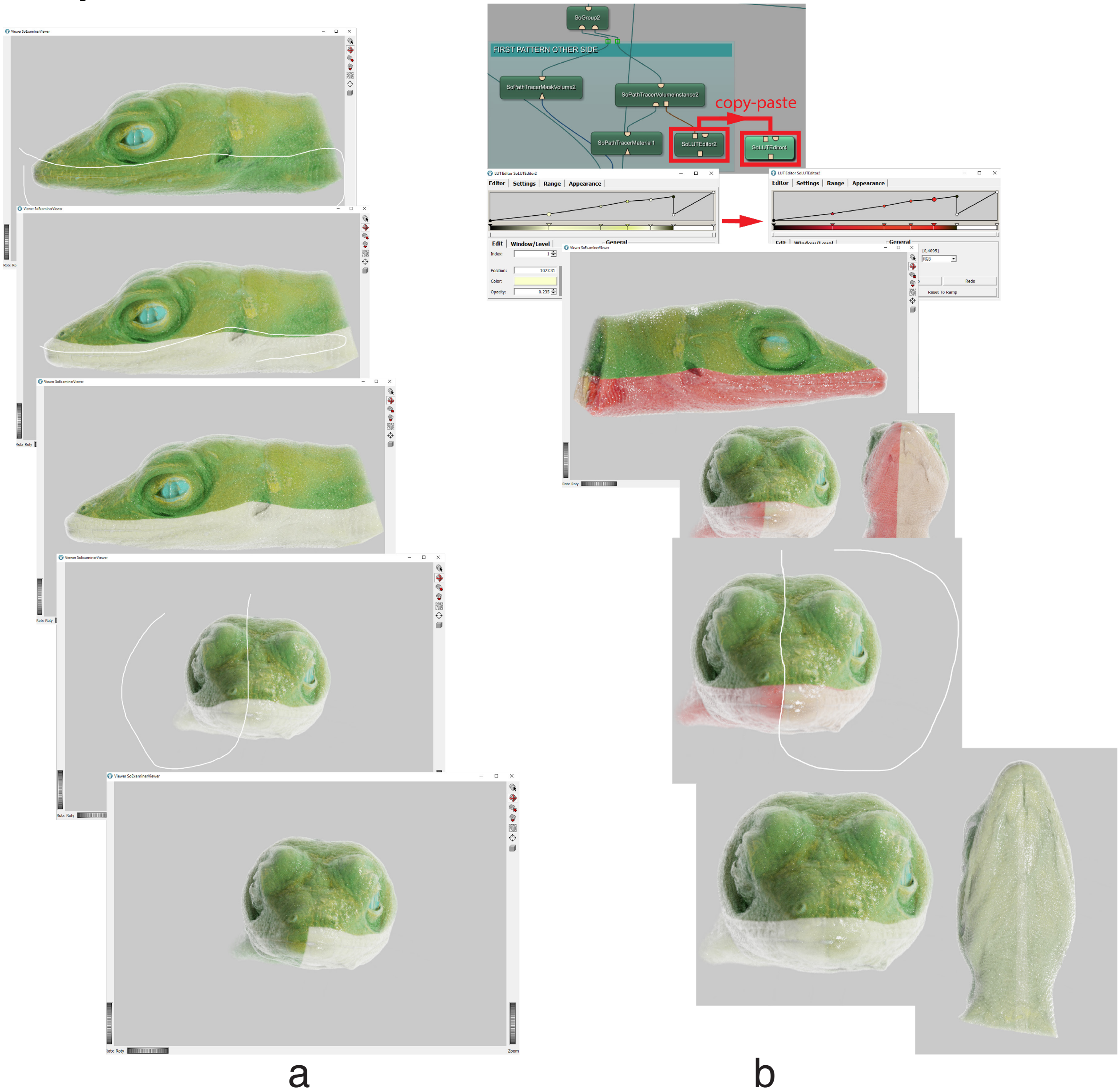

4. **Image acquisition**
   a. If required, save the masks corresponding to the generated patterns and adjust scene properties (see steps 2b,d in **Section 2.2**) before acquiring a high quality image of the scene following steps 4 a,b in in **Section 2.2**.

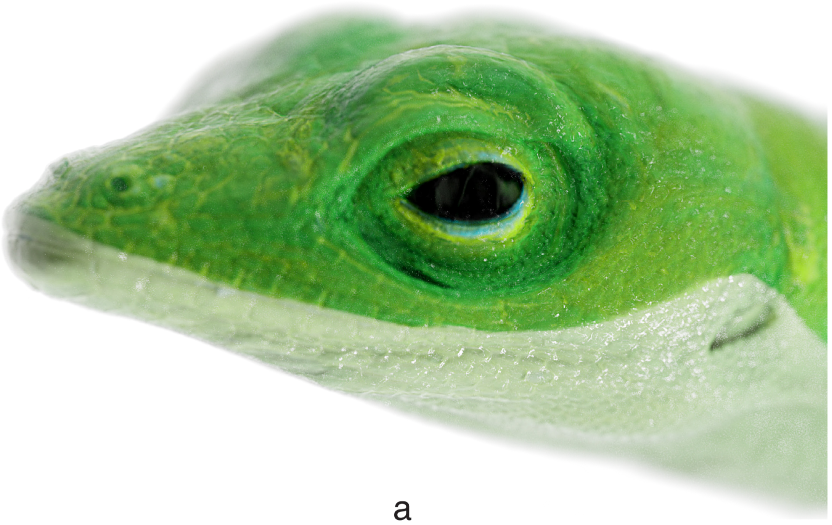

a

#### Section 7. Generation of cutaway views of multiple volumes with uniform internal structure and varied surface colors

As discussed in **Section 5**, the surface of anatomical parts often varies significantly from their internal appearance. This disparity can deeply affect the realism of visualizations, especially when creating cutaway views of these structures (**Figs. a,b,** below). The **SmARTR_Nested_Multi-Volume** network detailed in **Section 5** utilized a specifically designed module integration to independently render the internal and external regions of a volume, thereby enhancing the realism of single- or multi-volume cutaway views. Building on this concept, and on those expressed in steps 3*a-c of **Section 4.2**, the **SmARTR_Advanced_Multi-Volume** network described here extends these capabilities to multiple volumes. It enables the creation of enhanced cutaway views for volumes that share the same internal structure but have different surface colors. Examples include the differently colored parts of a lizard's body (**Fig. a,c**, below) or the distinct tissues or parts forming complex organs, such as the atrial and ventricular regions of the heart (**Figs. b,d**, below).

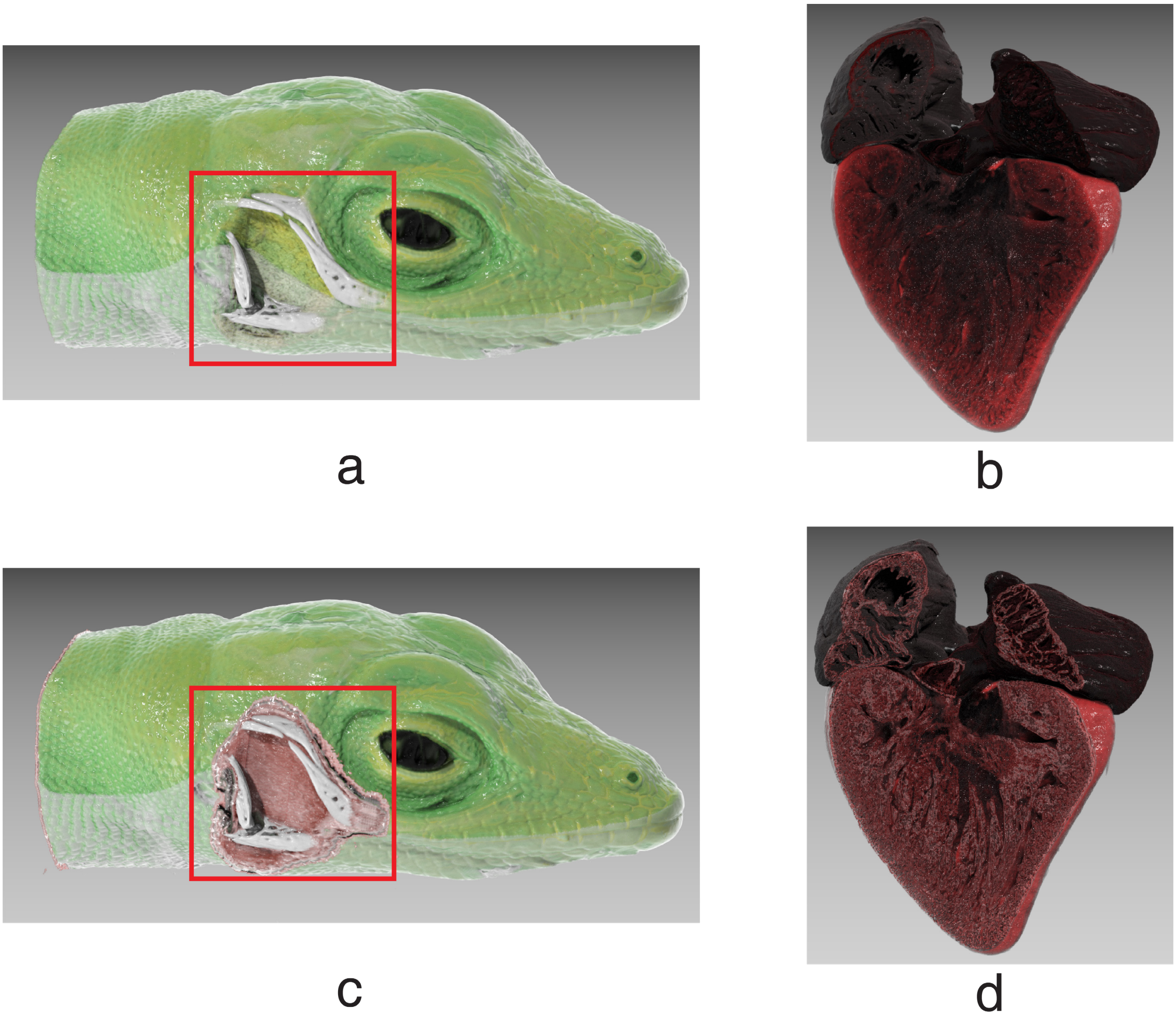

#### 7.1 The SmARTR_Advanced_Multi-Volume network overview

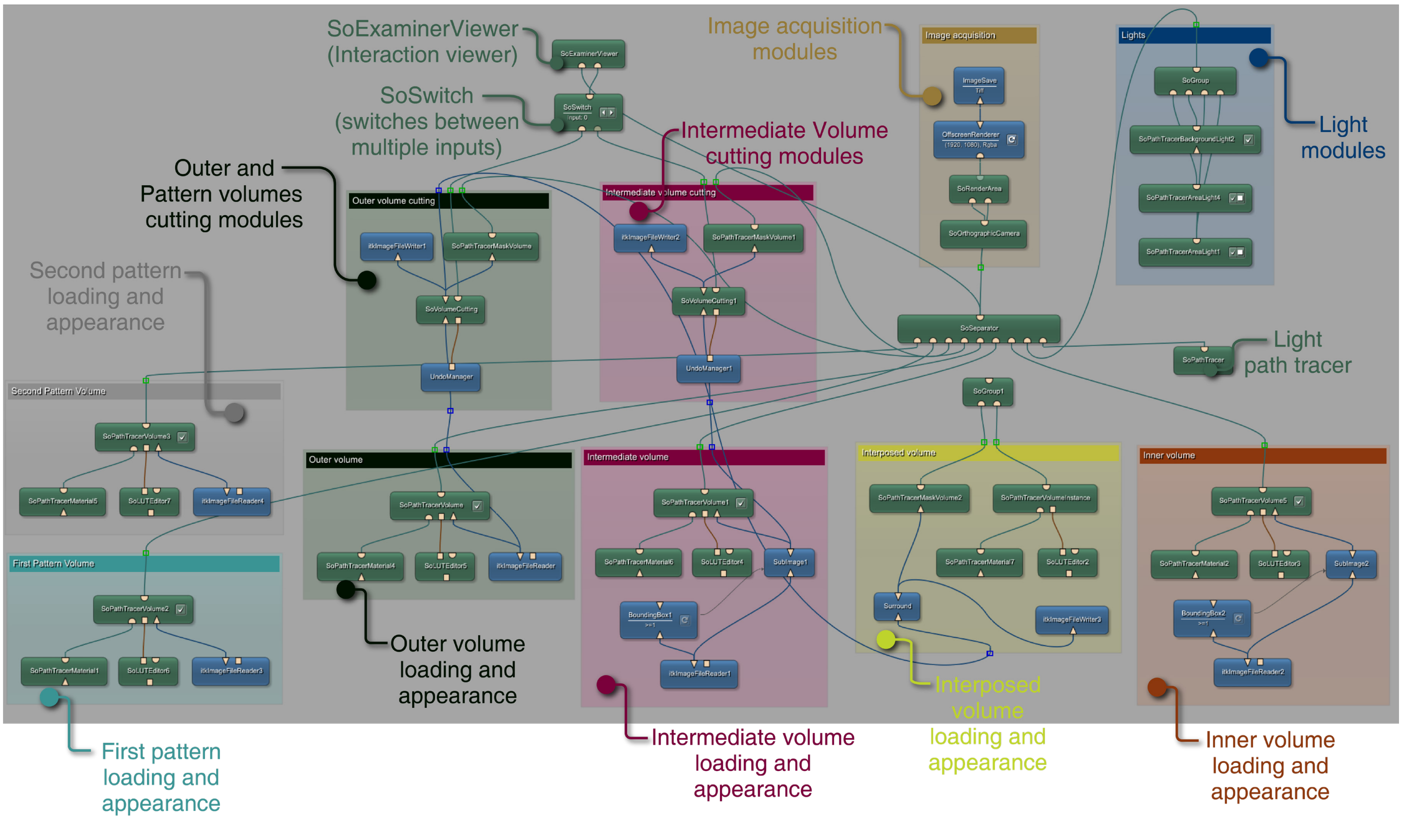

##### 7.2 Step-by-step protocol for advanced multi-volume rendering and acquisition

This section provides a practical example of generating photorealistic cutaway views of a lizard's head, displaying a dorso-ventral skin color pattern. Along with the specific network developed to generate the enhanced cutaway views, the workflow involves the use of the **SmARTR_Pattern_Drawing** network to create color pattern masks (see **Section 6**) and the **SmARTR_Volume_Extraction** network to extract the volumes corresponding to these color patterns **(**see **Section 4.2**, steps a-c). This approach is versatile and can also be applied to samples different from the one presented in the example, such as individual organs. In this case, the **SmARTR_Pattern_Drawing** network (see **Section 6**) can be used to create a mask that isolates one or more parts of an organ with distinct surface properties (**Figs. e,f**, below). Alternatively, binary masks for different organ or volume parts can be generated using external software. These masks can then be imported into the **SmARTR_Volume_Extraction** network to obtain the corresponding volumes (as described in step 2a, below). Once extracted, the volumes are loaded in one (or more, if needed) "**Pattern volume**" blocks in the **SmARTR_Advanced_Multi-Volume** network (see step 2b, below) while the original volume they were extracted from is loaded in "**Outer volume**" block.  Then, each volume appearance can be configured before proceeding with volume cutting operations (see step c, below).

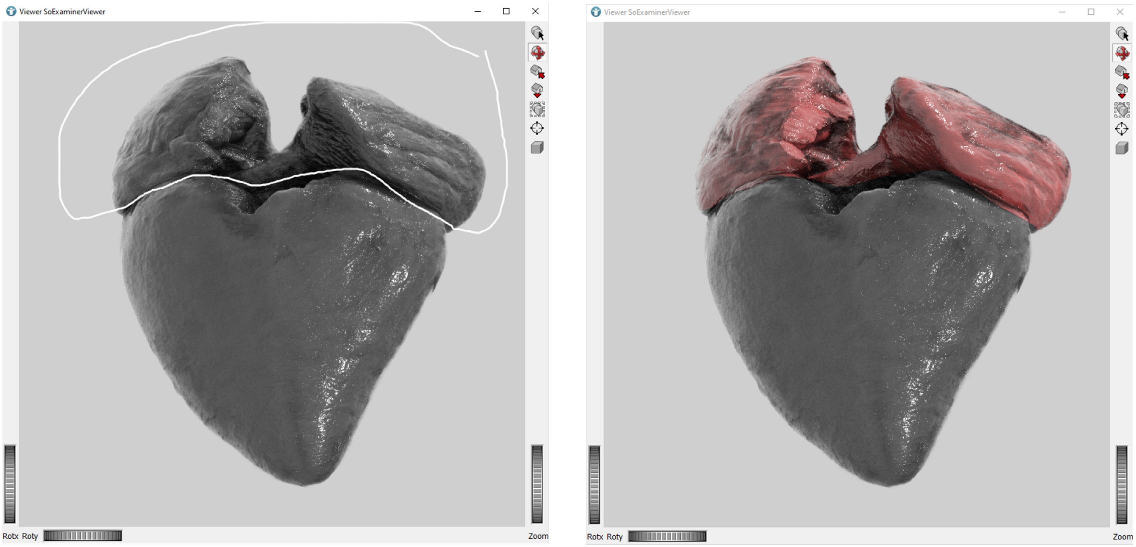


e f

1. **Draw patterns on the volume of interest following instructions in Section 6 and save them.**
2. **Extract the subvolumes corresponding to the generated pattern masks, adjust volume appearance and perform volume cutting operations**

a. Use the saved pattern masks to extract the corresponding volumes from the scan they derive from in the **SmARTR_Volume_Extraction** network.  Simply load the scan and the pattern mask files (the Anolis lizard head scan file and the mouth pattern mask, in the example) in the dedicated reading modules, check the results in the "**Output Inspector**" by clicking on the output of the "**Arithmetic**" module and save the extracted volume (for more details, see steps 1a-c in **Section 4.2**). Repeat these steps to extract additional pattern-related subvolumes (the Anolis lizard eye, in the example). When dealing with bilateral patterns that display the same appearance on both sides, as outlined in **Section 6**, step 3*, both masks corresponding to each side pattern can be loaded simultaneously and fed to the "**Arithmetic**" module input. This allows the extraction of the complete bilateral pattern as a single volume. Such a strategy simplifies downstream rendering and volume cutting operations. An additional "**itkImageFileReader**" and a "**Mask**" module must be first added to the workspace. Then, the two binary masks (loaded with the two "**itkImageFileReader**" modules now in the workspace) are fed into the "**Mask**" module configured to "**Mask In Original**" (this mask integration mode can be selected by double-clicking on the module balloon). The output from the "**Mask**" module is then connected to the right input of the "**Arithmetic**" module. If bilateral patterns consist of more than two binary masks, multiple "**itkImageFileReader**" and "**Mask**" (set to "**Mask In Original**") modules can be nested to feed all the bilateral pattern masks to the "**Arithmetic**" module simultaneously. Extracted volumes can then be saved as described in **Section 4.2**, step c).

b. Open the **SmARTR_Advanced_Multi-Volume** network and load the three nested volumes (the lizard head, skull, and brain, see **Section 5**). Load the extracted pattern volumes in the "**First Pattern Volume**" and "**Second Pattern Volume**" blocks and adjust the appearance of the volumes in the scene using the relevant modules (see steps 2a-f, **Section 2.2**). Similarly to what has been described in step 3*a **Section 4.2**, to allow multiple volume cutting by a single "**Volume Cutting**" block, the "**Mask Volume Name**" in the "**Tag/Mask Volume**" tab of the "**SoPathTracerVolume**" module relative to the volumes to be simultaneously cut (the "**Outer Volume**" and the "**First Pattern Volume**", in the example), must be identical ("**outer_and_pattern**", in the example). The same name must be specified in the "**Volume Name**" field of the "**SoPathTracerMaskVolume**" of the "**Outer Volume Cutting**" block.

c. Connect the "**SoGroup**" module from the "**Interposed volume**" block to the network. The appearance of the volumes in the scene will change upon connection of the "**SoGroup**" to the network. Simply draw a contour anywhere in the "**SoExaminerViewer**" to restore volume appearance **(**see step 2b, **Section 5.2)**. After this action, the necessary cutting operations can be performed and will affect all the volumes sharing the same "**Mask Volume Name**". Finally, modify the appearance of the "**Interposed volume**" to achieve a realistic representation of the parts exposed following volume cutting. In rare cases where extra material appears outside the area affected by cutting operations during the configuration of the 'Interposed volume,' refer to the instructions in **Section 5.2**, Step 2c. Enhanced semitransparent cutaway views can be obtained following the same steps detailed in **Section 5.**

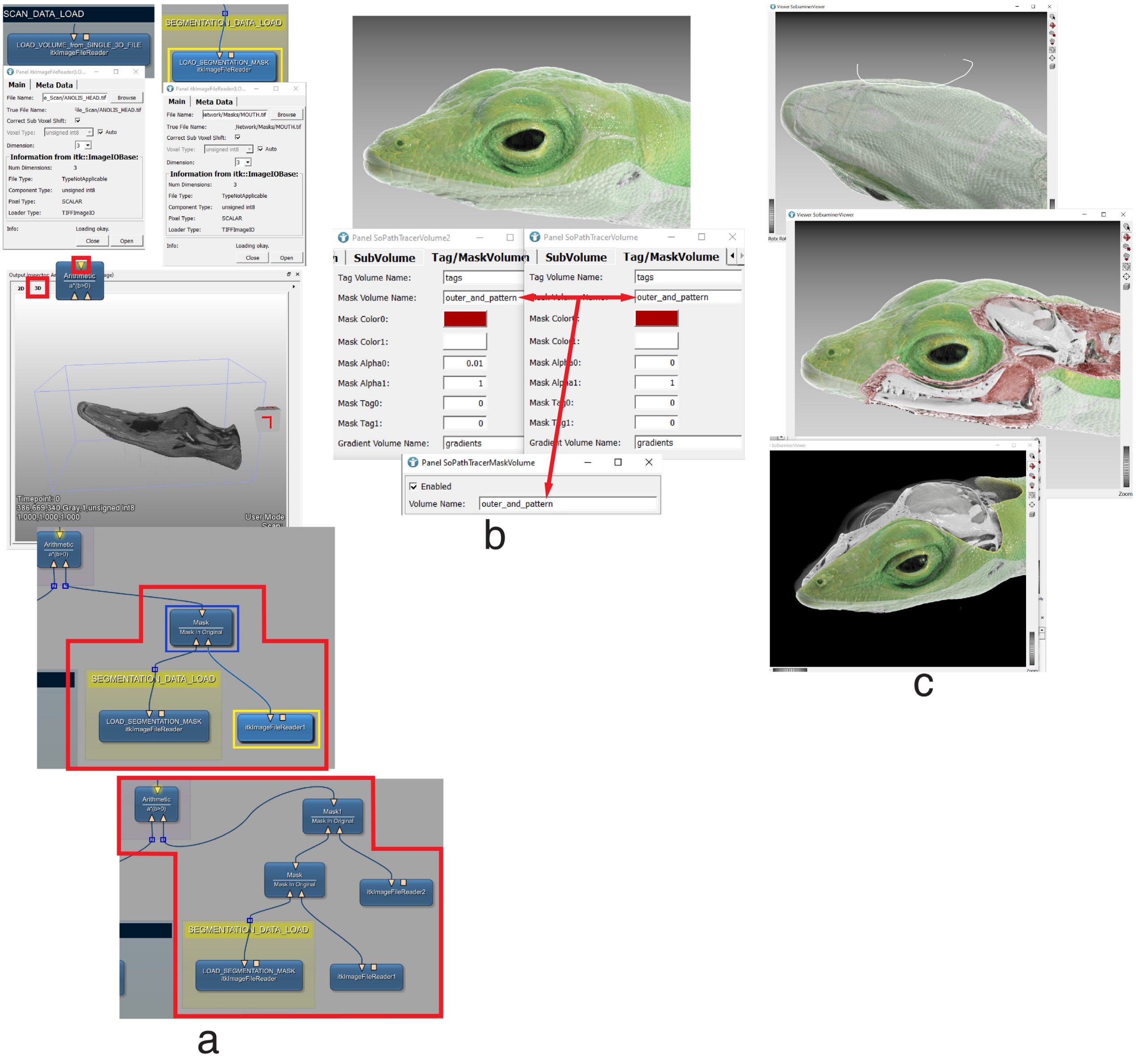
SCAN_DATA_LOAD
LOAD_VOLUME_from_SINGLE_3D_FILE
itkImageFileReader
SEGMENTATION_DATA_LOAD
LOAD_SEGMENTATION_MASK
itkImageFileReader
Arithmetic
a*(b>0)
Mask
Mask In Original
itkImageFileReader1
Mask1
itkImageFileReader2
a
Panel SoPathTracerVolume2
Panel SoPathTracerVolume
SubVolume
Tag/MaskVolume
Tag Volume Name:
tags
Mask Volume Name:
outer_and_pattern
Mask Color0:
Mask Color1:
Mask Alpha0:
0.01
Mask Alpha1:
1
Mask Tag0:
0
Mask Tag1:
0
Gradient Volume Name:
gradients
Panel SoPathTracerMaskVolume
Enabled
Volume Name:
outer_and_pattern
b
c

**Supplemental Figure S1**

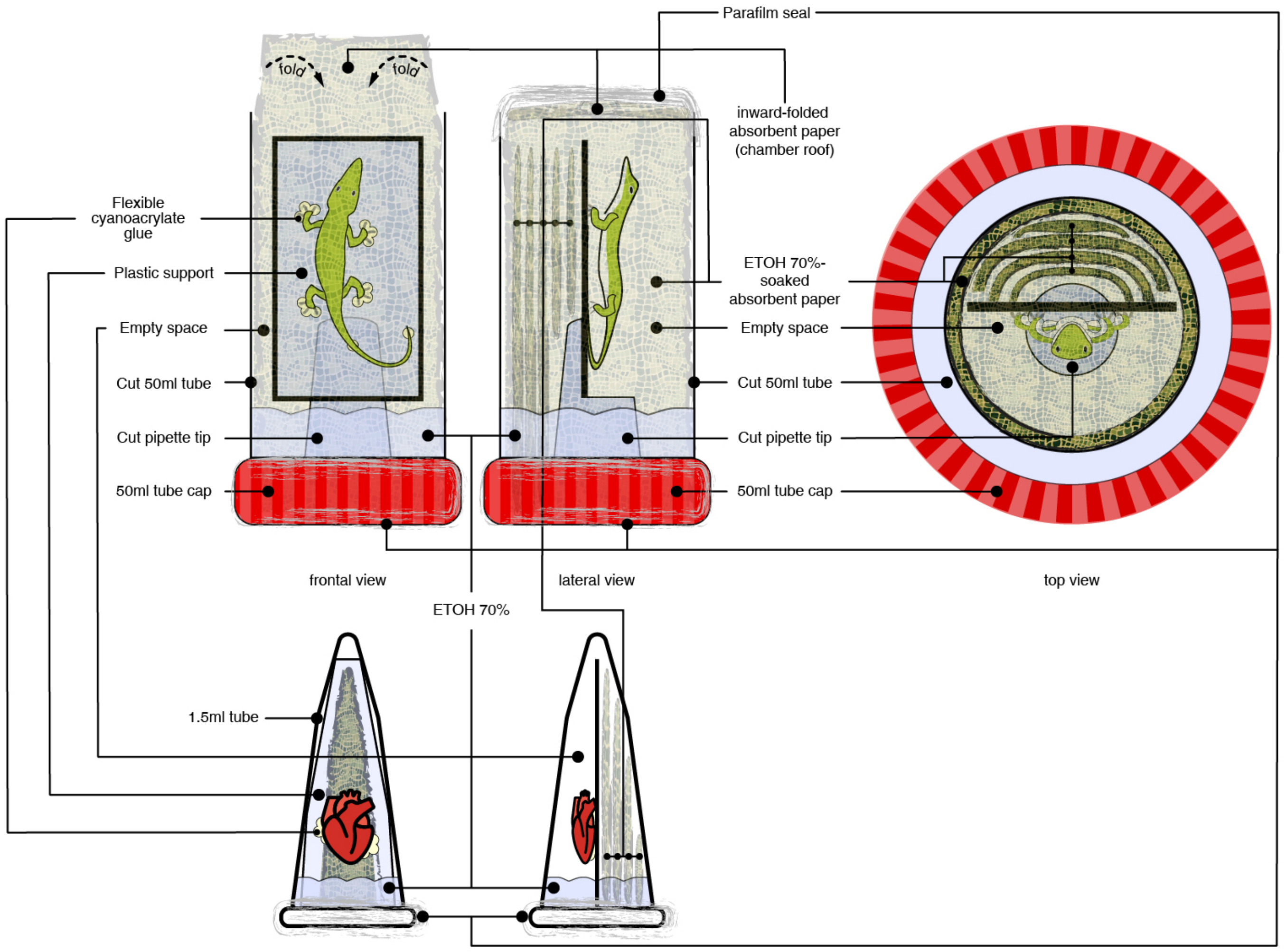


**Supplemental Figure S1. Sample mounting approach for intact specimens and dissected organs.** Schematic drawing summarizing humid chamber components and specimen mounting strategy for intact specimens (top panel) and isolated organs or animal parts (bottom panel) used in this study.

**Supplemental Table S1. List of specimens, staining, imaging and rendering parameters used in the study**

| Species | Intact specimen size* (cm) | Sample type | Contrast agent used | Approximate staining duration (days) | Scan voxel size (μm) | Rendering voxel size (μm) |
|---|---|---|---|---|---|---|
| ***Agalychnis callidryas*** | 2.2 | whole body | $I_2E_{70}$ | 14 | 5.75 | 20/15 |
| | | forelimb | $I_2E_{70}$ + $PTAE_{70}$ | 21 (14+7) | 2.75 | 6 |
| | | hindlimb | $I_2E_{70}$ + $PTAE_{70}$ | 21 (14+7) | 2.75 | 6 |
| | | tongue | $I_2E_{70}$ + $PTAE_{70}$ | 21 (14+7) | 2.15 | 5 |
| | | skeleton | none | x | 5.5 | 15/6 |
| ***Anolis carolinensis*** | 2.5 | whole body | $I_2E_{70}$ | 14 | 5.25 | 15 |
| | | skeleton | none | x | 5.25 | 15 |
| ***Aquilonastra burtoni*** | 1.1 | whole body | $PTAE_{70}$ | 14 | 3 | 10 |
| | | skeleton | none | x | 3 | 10 |
| ***Chameleo calyptratus*** | 3 | whole body | $I_2E_{70}$ | 14 | 20 | 25 |
| | | head | $I_2E_{70}$ | 14 | 4 | 10 |
| | | trunk | $I_2E_{70}$ | 14 | 9.5 | 10 |
| | | tongue | $I_2E_{70}$ + $PTAE_{70}$ | 21 (14+7) | 2.75 | 4 |
| | | hindlimb | $I_2E_{70}$ + $PTAE_{70}$ | 21 (14+7) | 2.75 | 5 |
| | | skeleton | none | x | 20 | 10 |
| ***Gastromyzon zebrinus*** | 4.2 | whole body | $I_2E_{70}$ | 14 | 20 | 21 |
| | | trunk | $I_2E_{70}$ | 14 | 3.75 | 10 |
| | | gills | $I_2E_{70}$ + $PTAE_{70}$ | 21 (14+7) | 2 | 5 |
| | | skeleton | none | x | 4.5 | 21 |
| ***Gryllus bimaculatus*** | 1.9 | whole body | $I_2E_{70}$ | 28 | 5 | 15 |
| | | digestive system | $PTAE_{70}$ | 14 | 3.15 | 15 |
| ***Mus musculus*** | 2.8 | whole body | $I_2E_{70}$ | 14 | 10 | 16 |
| | | head | $I_2E_{70}$ | 14 | 3.75 | 20 |
| | | trunk | $I_2E_{70}$ | 14 | 6.5 | 12 |
| | | heart | $I_2E_{70}$ + $PTAE_{70}$ | 28 (14+14) | 3 | 5 |
| | | lungs | $I_2E_{70}$ + $PTAE_{70}$ | 21 (14+7) | 3 | 8 |
| | | embryo 17.5 | $I_2E_{70}$ | 14 | 3 | 7 |
| | | skeleton | none | x | 10 | 12/16/20 |
| ***Pogona vitticeps*** | x | heart | $PTAE_{70}$ | 14 | 2.75 | 10 |
| | | forelimb | $PTAE_{70}$ | 21 | 2.75 | 6 |
| | | lungs | $PTAE_{70}$ | 14 | 5 | 5 |
| | | skin | $PTAE_{70}$ | 14 | 3 | 6 |
| ***Pterostichus oblongopunctatus*** | 1.1 | whole body | $I_2E_{70}$ | 30 | 6.5 | 8 |

** A. callidryas, A. carolinensis, C. calyptratus, M. musculus: snout to vent length; A. burtoni, G. zebrinus, G. bimaculatus, P. oblongopunctatus: total length*
